# Adversarial domain translation networks for fast and accurate integration of large-scale atlas-level single-cell datasets

**DOI:** 10.1101/2021.11.16.468892

**Authors:** Jia Zhao, Gefei Wang, Jingsi Ming, Zhixiang Lin, Yang Wang, The Tabula Microcebus Consortium, Angela Ruohao Wu, Can Yang

**Author notes:** These authors contributed to this work equally (Jia Zhao, Gefei Wang).

## Abstract

The rapid emergence of large-scale atlas-level single-cell RNA-seq datasets presents remarkable opportunities for broad and deep biological investigations through integrative analyses. However, harmonizing such datasets requires integration approaches to be not only computationally scalable, but also capable of preserving a wide range of fine-grained cell populations. We created Portal, a unified framework of adversarial domain translation to learn harmonized representations of datasets. With innovation in model and algorithm designs, Portal achieves superior performance in preserving biological variation during integration, while achieving integration of millions of cells in minutes with low memory consumption. We show that Portal is widely applicable to integrating datasets across samples, platforms and data types (including scRNA-seq, snRNA-seq and scATAC-seq). Finally, we demonstrate the power of Portal by applying it to the integration of cross-species datasets with limited shared information among them, elucidating biological insights into the similarities and divergences in the spermatogenesis process among mouse, macaque and human.

## Introduction

Advances in single-cell sequencing have enabled identification of novel cell types [1, 2], investigation of gene regulation networks [3, 4], and understanding of cellular differentiation processes [5, 6]. As single-cell technologies rapidly evolved over recent years, its experimental throughput substantially increased, allowing researchers to profile increasingly complex and diverse samples, and accelerating the accumulation of vast numbers of rich datasets over time [7, 8, 9]. Integrative and comparative analyses of such large-scale datasets originating from various samples, different platforms and data modalities, as well as across multiple species, offer unprecedented opportunities to establish a comprehensive picture of diverse cellular behaviors. Integration is a critical step, to account for heterogeneity of different data sources when taking advantage of single-cell data from different studies [10]. Thus, integration methods that can efficiently and accurately harmonize a wide range of data sources are essential for accelerating life sciences research [11].

Although integration methods for single-cell transcriptomics analysis have evolved along with single-cell sequencing technologies, the rapid accumulation of new and diverse single-cell datasets has introduced three major challenges to the integration task. First, as the sample size of each single-cell dataset grows dramatically, numerous extensive datasets with hundreds of thousands or even millions of cells have been produced [8, 9, 12]. The emergence of large-scale datasets requires integration methods to be fast, memory-efficient, and scalable to millions of cells. Second, technology now allows effective, comprehensive characterization of complex organs, containing rare subpopulations of cells that can now be captured, albeit in small numbers, thanks to the scale of profiling that is now possible [7, 13]. Investigation into high-level heterogeneity among cell populations is essential for understanding the mechanism of complex biological systems. Hence, the ideal integration method needs to carefully preserve fine-grained cell populations from each atlas-level dataset. Third, the biological origins of datasets has expanded in diversity, with data now spanning across not only different technological platforms and data types, different individual donors, but even across different species, which can be especially interesting for evolutionary studies [14, 15, 16]. Integrative analysis of such diverse datasets would allow researchers to unify resources to address a wider range of biological questions. Recent single-cell atlasing efforts are a primary example of these challenges – various human tissue atlases [12, 17], mouse multi-tissue atlases [7, 18], and non-human primate atlases [19, 20] have been generated, culminating in data from millions of single cells and single nuclei. Both within and across atlas comparisons are of interest. To perform integrative and comparative analyses based on such diverse data sources, there is an urgent need for methods that can flexibly account for heterogeneous dataset-specific effects, while maintaining a high level of integration accuracy.

Many methods have been developed to align single-cell datasets [10], including Harmony [21], Seurat [22], online iNMF [23], VIPCCA [24], scVI [25], fastMNN [26], Scanorama [27] and BBKNN [28]. Several of these methods that were designed for large datasets at the time of publication are now less attractive in terms of scalability in the face of atlas-level dataset sizes. For instance, a representative category of methods leverages the mutual nearest neighbors (MNN) to perform data alignment. These MNN-based methods, such as Seurat, fastMNN and Scanorama, require identification of MNN pairs across datasets, thus the time and memory costs quickly become unbearably high when the dataset exceeds one million cells. Another limitation of existing methods is that they are mainly targeted towards integrating datasets of less complex tissues, utilizing strategies such as MNN, matrix factorization, and soft-clustering to capture major biological variations. With these strategies, inaccurate mixing of different cell types can be avoided when clear clustering patterns are present; but when dealing with more complex tissues, they tend to overcorrect fine-grained cell subpopulations, resulting in the loss of power in revealing interesting biological variations [29, 30]. Lastly, most existing methods are designed to correct batch effects caused by technical artifacts. To this end, a number of methods, like BBKNN and fastMNN, assume that the biological variation is much larger than the variation of batch effects. This assumption may not be true when applied across data types and species.

To simultaneously address the above three challenges, we created Portal, a machine learning-based algorithm for aligning atlas-level single-cell datasets with high efficiency, flexibility, and accuracy. Viewing datasets from different studies as distinct domains with domain-specific effects (including technical variation and other sources of unwanted variation), Portal achieves extraordinary data alignment performance through a unified framework of domain translation networks that incorporates an adversarial learning mechanism [31]. To find the correspondence between two domains, our domain translation network utilizes an encoder to embed cells from one domain into a latent space where domain-specific effects are removed, and then uses a generator to map latent codes to another domain. The generator simulates the generation process of domain-specific effects. In each domain, a discriminator is trained to identify where poor alignment between the distributions of original cells and transferred cells occurs. The feedback signal from the discriminator is used to strengthen the domain translation network for better alignment. The nonlinearity of encoders and generators in the adversarial domain translation framework enables Portal to account for complex domain-specific effects. In contrast to existing domain translation methods [32, 33, 34], Portal has the following unique features. First, Portal has a uniquely designed discriminator which can adaptively distinguish domain-shared cell types and domain-unique cell types. Therefore, Portal will not force the alignment of domain-unique cell types, avoiding the risk of overcorrection. Second, without using any cell type label information, three regularizers of Portal can guide domain translation networks to find correct correspondence between domains, account for domain-specific effects, and retain biological variation in the latent space. Third, through a tailored design of lightweight neural networks and mini-batch optimization accelerated by graphics processing units (GPUs), Portal can scale up to datasets containing millions of cells in minutes with nearly constant memory usage. With the above innovations in model and algorithm designs, Portal enables fast and accurate integration of atlas-level datasets across samples, technological platforms, data modalities, and species.

Through a comprehensive benchmarking study, where integration of heterogeneous collections of atlas-level single-cell RNA sequencing (scRNA-seq) data are included, Portal shows its superiority over state-of-the-art alignment algorithms in terms of both computational efficiency and accuracy. We then show that Portal can accurately align cells from complex tissues profiled by scRNA-seq and single-nucleus RNA sequencing (snRNA-seq) as well as align scRNA-seq data and single-cell assay for transposase-accessible chromatin using sequencing (scATAC-seq) data, even in the presence of highly unbalanced cell type compositions. We also apply Portal to the integration of cells in differentiation processes, especially the alignment of the gradient of cells in the spermatogenesis process across multiple species (mouse, macaque, and human). Using these diverse and challenging experiments, we demonstrate Portal’s versatility and power for a broad range of applications. Comprehensive analyses of real, expert annotated data confirm that integrated cell embeddings provided by Portal can be reliably used for identification of rare cell populations via clustering or label transfer, studies of differentiation trajectories, and transfer learning across data types and across species. Portal is now publicly available as a Python package (https://github.com/YangLabHKUST/Portal), serving as an efficient, reliable and flexible tool for integrative analyses.

## Results

### Method Overview: Portal learns a harmonized representation of different datasets with adversarial domain translation

Expression measurements from different datasets fall into different domains due to the existence of domain-specific effects, including technical variation and other sources of unwanted variation (Fig. 1**a**), causing difficulty when performing joint analyses. Without loss of generality, here we consider two domains, 𝒳 and 𝒴. We assume that domain 𝒳 and domain 𝒴 can be connected through a low-dimensional shared latent space Ƶ, which captures the biological variation and is not affected by the domain-specific effects. By taking the measurements of cells from 𝒳 and 𝒴 as inputs, we aim to learn a harmonized representation of cells in latent space Ƶ to obtain data alignment between 𝒳 and 𝒴.

**Figure 1:**
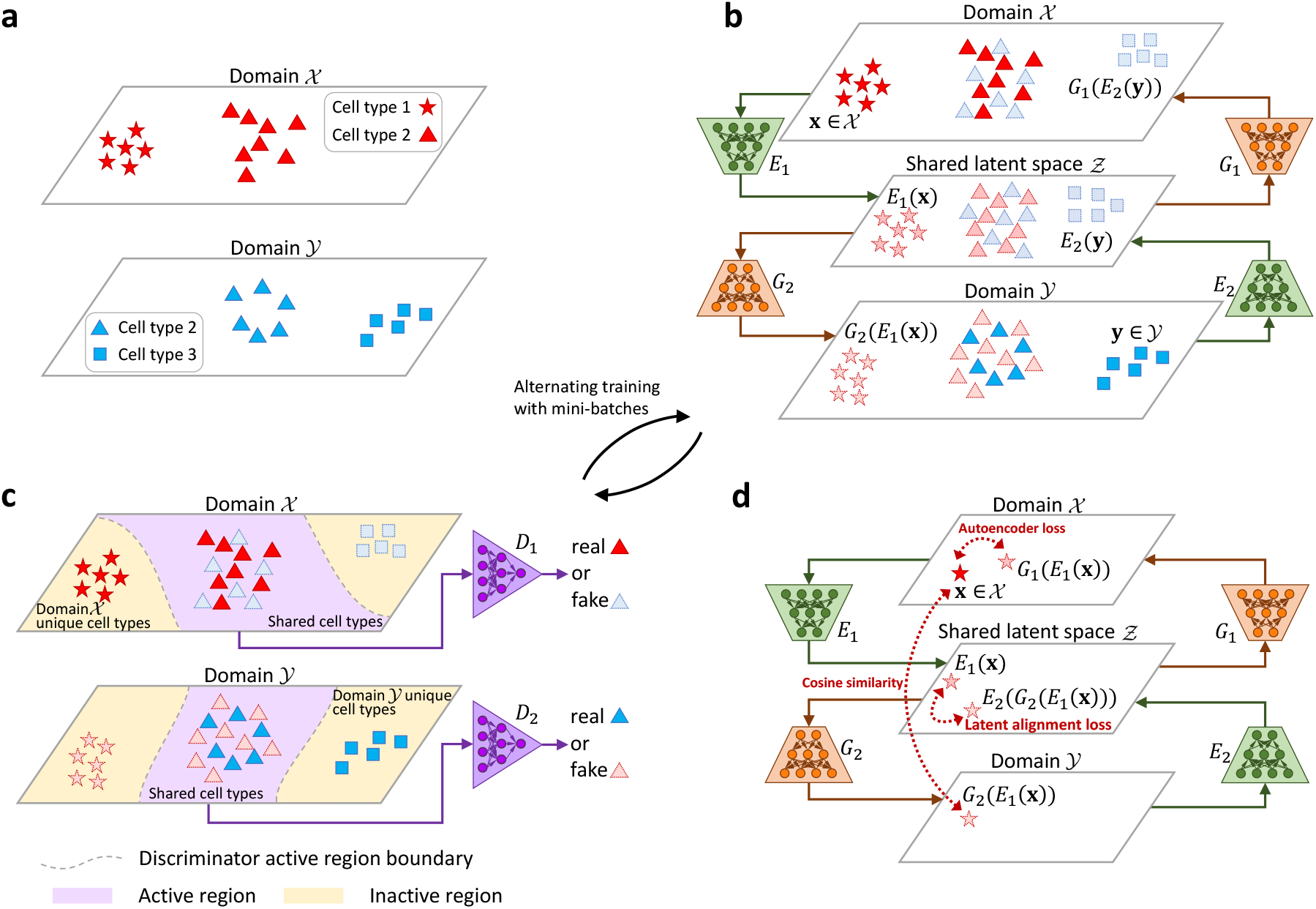
Overview of Portal. **a**. Portal regards different single-cell datasets as different domains. Joint analyses of these datasets are confounded by domain-specific effects, representing the unwanted technical variation. **b**. Portal employs encoders *E*_1_(·), *E*_2_(·) to embed the biological variation of domains 𝒳 and 𝒴 into a shared latent space Ƶ, where domain-specific effects are removed. The generating process of domain-specific effects are captured by two generators *G*_1_(·) and *G*_2_(·). Encoder *E*_1_(·) and generator *G*_2_(·) form a domain translation network *G*_2_(*E*_1_(·)) mapping from 𝒳 to 𝒴; Encoder *E*_2_(·) and generator *G*_1_(·) form another domain translation network mapping from 𝒴 to 𝒳. **c**. Encoders and generators are trained by competing against specially designed discriminators *D*_1_(·) and *D*_2_(·). In each domain, a discriminator is trained to distinguish between original cells in this domain and cells transferred from another domain, providing feedback signals to assist alignment. To prevent overcorrection of domain-unique cell types, the discriminators in Portal with the tailored design are also able to distinguish between domain-unique cell types and domain-shared cell types. With this design, Portal can focus only on merging cells of high probability to be of domain-shared cell types, while it remains inactive on cells of domain-unique cell types. **d**. Portal leverages three regularizers to help it find correct and consistent correspondence across domains, including the autoencoder regularizer, the latent alignment regularizer and the cosine similarity regularizer.

We achieve the above goal through a unified framework of adversarial domain translation, namely “Portal”. Domains and the shared latent space are connected by encoders and generators (Fig. 1**b**). Encoder *E*_1_(·) : 𝒳 → Ƶ is designed to remove the domain-specific effects when mapping cells from 𝒳 into Ƶ, and generator *G*_1_(·) : Ƶ → 𝒳 is designed to simulate the domain-specific effects when mapping cells from Ƶ into 𝒳. By symmetry, encoder *E*_2_(·) : 𝒴 → Ƶ and generator *G*_2_(·) : Ƶ → 𝒴 are designed with the same role in connecting 𝒴 and Ƶ. To transfer cells between 𝒴 and 𝒳 through shared latent space Ƶ (Fig. 1**b**), encoder *E*_2_(·) and generator *G*_1_(·) work together to form one domain translation network *G*_1_(*E*_2_(·)) : 𝒴 → Ƶ → 𝒳. Clearly, encoder *E*_1_(·) and generator *G*_2_(·) form another domain translation network *G*_2_(*E*_1_(·)) : 𝒳 → Ƶ → 𝒴. To achieve the mixing of original cells and transferred cells, discriminators *D*_1_(·) and *D*_2_(·) are deployed in domains 𝒳 and 𝒴 to identify where poor mixing occurs (Fig. 1**c**). The discriminators’ feedback then guides the domain translation networks to improve the mixing.

However, the well mixing of original cells and transferred cells in each domain does not imply extraordinary data alignment across domains. First, a domain-unique cell population should not be mixed with cells from another domain. Second, cell types *A* and *B* in domain 𝒳 could be incorrectly aligned with cell types *B* and *A* in domain 𝒴, respectively, although the distributions of original cells and transferred cells are well mixed. To address these issues, Portal has the following unique features, which distinguishes it from existing adversarial domain translation frameworks [32, 33]. On one hand, we deploy the tailored design of discriminators *D*_1_(·) and *D*_2_(·) such that they can distinguish domain-unique cell types from cell types shared across different domains. The domain-unique cell types will be treated as outliers and left in the discriminator’s inactive region (Fig. 1**c**). In such a way, these cell types will not be enforced for alignment, avoiding the risk of overcorrection. On the other hand, we design three regularizers to find correct correspondence across domains and avoid incorrect alignment when the distributions are well mixed.

Specifically, let **x** and **y** be the samples from domains 𝒳 and 𝒴, respectively. We consider the following framework of adversarial domain translation,

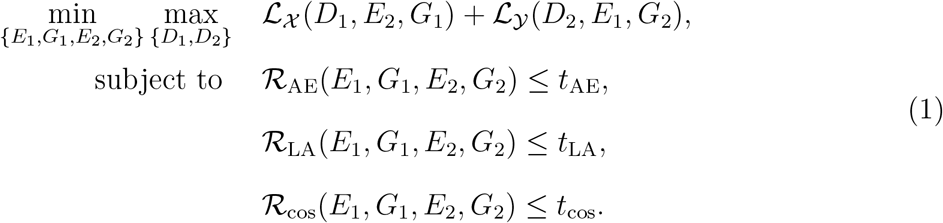

In model (1), ℒ_𝒳_ (*D*_1_, *E*_2_, *G*_1_) := 𝔼[log *D*_1_(**x**)]+𝔼[log(1−*D*_1_(*G*_1_(*E*_2_(**y**))))] and ℒ_𝒴_(*D*_2_, *E*_1_, *G*_2_) := 𝔼[log *D*_2_(**y**)] + 𝔼[log(1 − *D*_2_(*G*_2_(*E*_1_(**x**))))] are the objective functions for adversarial learning of domain translation networks *G*_1_(*E*_2_(·)) and *G*_2_(*E*_1_(·)) in 𝒳 and 𝒴, respectively. Discriminators *D*_1_(·) and *D*_2_(·) are trained to distinguish between “real” cells (i.e. original cells in a domain), and “fake” cells (i.e. transferred cells generated by domain translation networks) by minimizing ℒ_𝒳_ + ℒ_𝒴_, while the domain translation networks are trained against the discriminators by maximizing ℒ_𝒳_ +ℒ_𝒴_. These three regularizers ℛ_AE_, ℛ_LA_ and ℛ_cos_ play a critical role in finding correct correspondence of cells between two domains, accounting for domain-specific effects, and retaining biological variation in the latent space (Fig. 1**d**). More specifically, the first regularizer 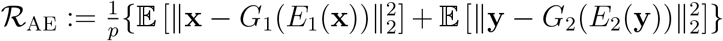, where *p* is the dimensionality of domains 𝒳 and 𝒴, requires the autoencoder consistency in domains 𝒳 and 𝒴; the second regularizer 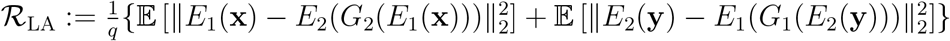, where *q* is the dimensionality of Ƶ, imposes the consistency constraint in the latent space; and the third regularizer 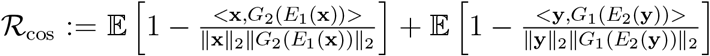 introduces the cross-domain correspondence by preserving the cosine similarity between a sample and its transferred version; *t*_AE_, *t*_LA_ and *t*_cos_ are their corresponding constraint parameters. More detailed explanation can be found in the Methods section.

We solve the above optimization problem via alternating updates by stochastic gradient descent. The algorithm is extremely computationally efficient with the support of stochastic optimization accelerated by GPUs. After the training process, Portal learns a harmonized representation of different domains in shared latent space Ƶ. Samples from 𝒳 and 𝒴 can be transferred into latent space Ƶ to form an integrated dataset {*E*_1_(**x**)}_**x**∈𝒳_ ∪ {*E*_2_(**y**)}_**y**∈𝒴_ using encoders *E*_1_(·) and *E*_2_(·), facilitating the downstream integrative analysis of cross-domain single-cell datasets.

### Accurate integration of atlas-level datasets within minutes and requiring lower memory consumption compared to other methods

The rapid accumulation of large-scale single-cell datasets requires integration algorithms to efficiently handle datasets containing millions of cells without loss of accuracy. For a comprehensive comparison, we first benchmarked Portal and existing representative methods, including Harmony [21], Seurat v3 [22], online iNMF [23], VIPCCA [24], scVI [25], fastMNN [26], Scanorama [27] and BBKNN [28], in terms of integration performance following a recent benchmarking study [30]. Using a number of scRNA-seq datasets from diverse tissue types with curated cell cluster annotations, including mouse spleen, marrow, and bladder [7], we quantitatively evaluated the integration performance of each method. We first evaluated alignment performance, which can sometimes be interpreted as batch correction performance, of all compared methods. The score for batch correction was computed by leveraging a collection of batch correction metrics designed in existing studies, including k-nearest neighbor batch-effect test (kBET) [35], principal component regression of the batch covariate (PCR batch) [35], average silhouette width across batches (batch ASW) [35], graph integration local inverse Simpson’s Index (graph iLISI) [30, 21] and graph connectivity [30]. The higher the batch correction score, the higher the degree of mixing across datasets. We also assessed the score for conservation of biological variation using different metrics, including adjusted rand index (ARI) [36], normalized mutual information (NMI) [37], cell type ASW, graph cell type local inverse Simpson’s Index (graph cLISI) [30, 21], isolated label F1 [30], isolated label silhouette [30] and cell cycle conservation [30]. By jointly accounting for these metrics, the score can be used to evaluate different methods’ ability to preserve information such as cell type identities. Inappropriate merging of cell types during integration will result in a low score of biological variation conservation. Finally, we computed the overall score as a 40:60 weighted average of the batch correction score and the conservation of biological variation score to indicate the overall integration performance. Based on our benchmarking results, we found that in general, BBKNN, Scanorama, fastMNN, scVI and VIPCCA had less satisfactory overall integration performance compared to the other four methods (Fig. 2**a**, the first three columns; Figs. S5, S7, S9 and S11). As indicated by the relatively low batch correction scores of BBKNN, Scanorama, fastMNN and scVI, we found that observable batch effects still exist in the integration results that they produced (Figs. S6, S8 and S10). Although VIPCCA showed reasonable performance in terms of removing batch effects, incorrect mixing of distinct cell types was often observed in VIPCCA’s integration results (Fig. S6). Therefore, its overall scores are relatively low due to the loss of biological variation (Figs. S5, S7).

**Figure 2:**
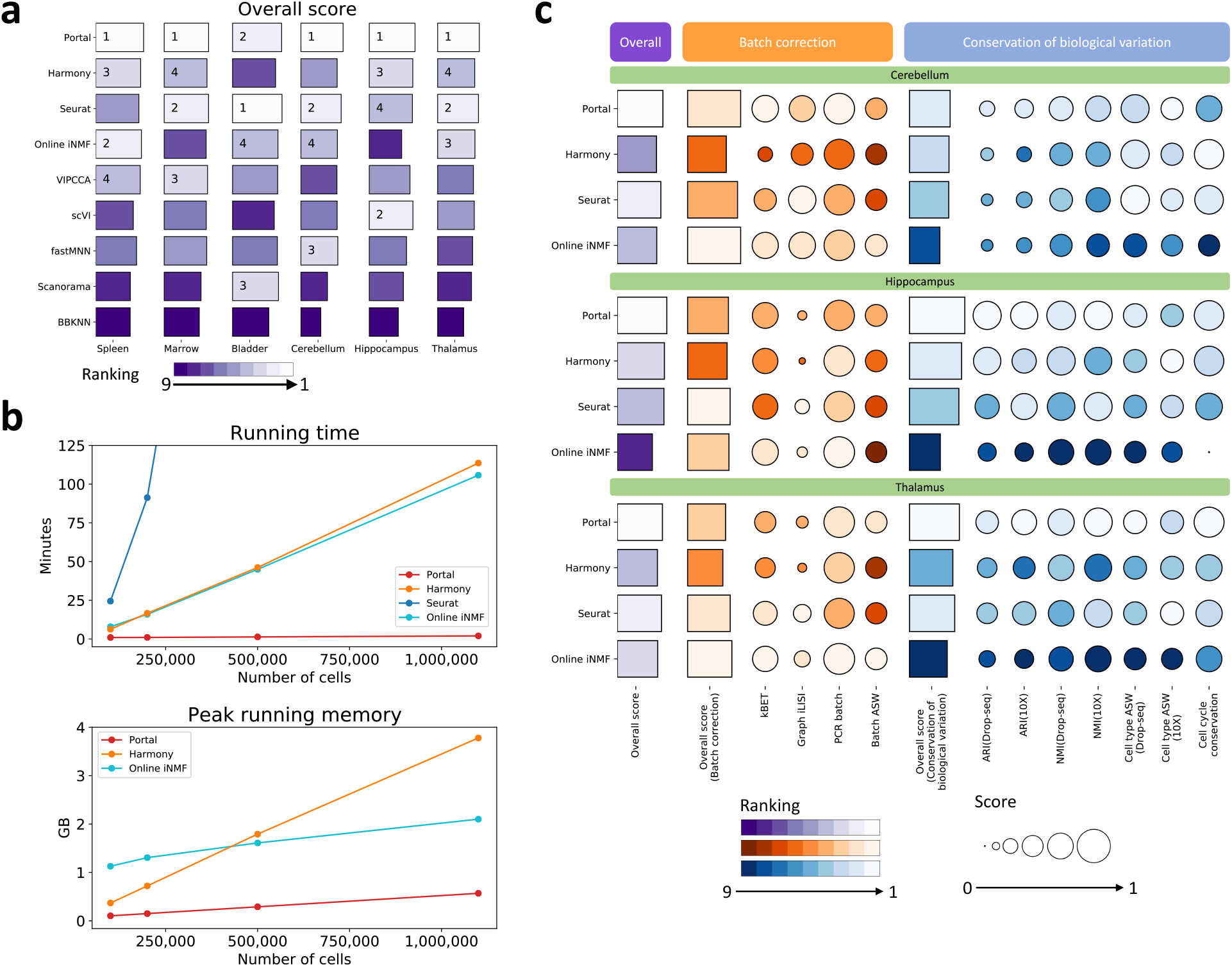
Benchmarking of Portal and other state-of-the-art integration methods. **a**. Overall scores of the compared methods evaluated on mouse spleen, marrow, bladder, cerebellum, hippocampus and thalamus datasets. The ranking was visualized by color gradient, where lighter color indicates better performance. **b**. The running time and the peak running memory required by the benchmarked methods. The datasets were sampled from two mouse brain atlas datasets (*n* = 100, 000, 250, 000, 500, 000, and 1, 100, 167). Seurat required 24.52 GB on the dataset with 100, 000 cells, which was not comparable to the other three benchmarked methods in terms of the peak running memory usage. **c**. Batch correction and biological variation conservation evaluated using three shared tissues from two mouse brain atlases (profiled by Drop-seq and 10X), including cerebellum, hippocampus, and thalamus. Biological variation conservation performance was assessed based on fine-grained annotations provided by the original publications [8, 9].

Among those methods with high user popularity, Harmony, Seurat, and online iNMF also showed the best overall integration performance results (Fig. 2**a**, the first three columns; Figs. S6, S8 and S10). To offer precise and robust integration performance, Seurat [22] utilizes the detection of mutual nearest neighbors (MNN) to build correspondence between datasets in the shared embedding space obtained by applying canonical correlation analysis (CCA). Harmony [21] learns a simple linear correction for dataset-specific effects by running an iterative soft clustering algorithm, enabling fast computation on large datasets. Online iNMF [23] is a recently developed approach based on widely used integration method LIGER [38]. It extends LIGER’s non-negative matrix factorization to an iterative and incremental version to improve its scalability, while it has nearly the same performance as LIGER. For the remainder of this study, we focus our discussion on comparisons between Portal and these three high-performing and popular methods in the main text. The comparisons with other methods are provided in Fig. 2**a** (the last three columns), and Supplementary Information (Figs. S12 - S18).

Next, we evaluated the speed, memory usage, alignment quality, and integration accuracy using a more challenging integration task. We used two mouse brain atlases [8, 9] as bench-marking datasets for a more in-depth comparison of Portal and three other methods. One atlas contains Drop-seq data of 939,489 cells, and another one contains 10X Genomics (10X) data of 160,678 cells. These two mouse brain atlases have data from three shared brain regions: cerebellum, hippocampus, and thalamus. There are many small clusters of neuron subtypes in these datasets, where gene expressions between subclusters could have a relatively small difference. Thus, these datasets are more challenging to integrate compared to data with clear clustering patterns.

First, Portal has superior integration accuracy even when handling datasets which contain many subclusters with small difference. The score of biological variation conservation shows that Portal outperforms other state-of-the-art methods in cluster identity preservation, as the scores were assessed based on fine-grained cell type and subtype annotations. In particular, for all three brain regions tested, Portal has the highest ARI and NMI scores among the compared methods (Fig. 2**c**).

Second, Portal also outperforms the other three methods on scalability, in terms of time and memory consumption. For this benchmarking test, we obtained datasets from the original full-sized datasets by combining the two atlases and subsampling proportionally from each atlas, with each dataset having increasing sample size ranging from 100,000 to 1,100,167 (full dataset). The running time and the peak running memory of all methods were recorded using these datasets on the same GPU server. The results show that Portal’s running time and peak running memory remained almost constant even when the sample size increased dramatically (Fig. 2**b**). Compared to the other three methods, the running time required by Portal was also substantially less. On the dataset containing 500,000 cells, Portal’s running time was 80 seconds; when number of cells grew to 1,100,167, Portal’s running time only increased to 120 seconds. In comparison, Harmony and online iNMF both needed more than 40 minutes to integrate 500,000 cells and more than 100 minutes to complete the integration of the full dataset. The running time of Seurat increased most rapidly among the compared methods. It took as much as 511 minutes (over 8.5 hours) to integrate the 500,000-cell dataset. The computational efficiency of Portal is owing to two important factors in its design: 1) its algorithm takes advantage of GPU-accelerated stochastic optimization, such that Portal reads data in mini-batches from the disk rather than having to load the entire dataset at once, which enables fast integration of large single-cell datasets using small amounts of memory; and 2) lightweight neural networks are adopted in Portal to further improve computational efficiency. As such, Portal is also the most memory-efficient approach among the benchmarked methods (Fig. 2**b**). Peak running memory required by Portal ranged from 0.29 GB on 500,000-cell dataset to 0.57 GB on the full million-cell dataset. Notably, Portal’s lightweight networks and mini-batch stochastic optimization algorithm enable us to control the GPU peak running memory usage at a constant level of 0.06 GB. Among compared methods, online iNMF used less memory than Harmony and Seurat when the sample size became larger than 500,000, because it is also trained in mini-batches. However, its peak running memory was 2.10 GB on the million-cell dataset, which is 2.7 times more than Portal’s. Seurat required remarkably more memory usage than the other three methods. For clarity of visualization, we did not display the peak running memory required by Seurat as it ranged from 24.52 GB on the 100,000-cell dataset to 276.41 GB on the 500,000-cell dataset.

Finally, and importantly, Portal’s high performance in speed and memory consumption does not compromise its ability to align cell type clusters. The batch correction score shows that Portal’s alignment ability is comparable to, if not better than, the other state-of-the-art methods, indicating that Portal is capable to effectively remove domain-specific effects.

### Portal preserves subcluster and small cluster identities in complex tissues thereby facilitating identification of rare subpopulations

When integrating complex tissues, one problem that can arise is the inadvertent loss of small cell populations and subpopulations. Due to more nuanced differences between clusters, or due to the imbalance in cell numbers for very small cell populations, these “fine-grained” groups of cells may become inappropriately combined with other groups after integration. In the brain, for example, there are many subpopulations of neurons which are distinguished from each other using a few key gene markers while still all bearing the neuron signature; furthermore, some of these neuronal subtypes could be rare compared to other subtypes. To demonstrate that Portal can preserve the nuanced information of such small cell populations and subpopulations, we performed further analysis on the mouse hippocampus tissue integration results. Both mouse brain atlas datasets contain extensive data for this brain region (Fig. 3), and both studies identified a wide range of transcriptionally distinct cell subpopulations, including a variety of neuron subtypes, whose nuanced transcriptional differences should ideally be preserved by integration methods.

**Figure 3:**
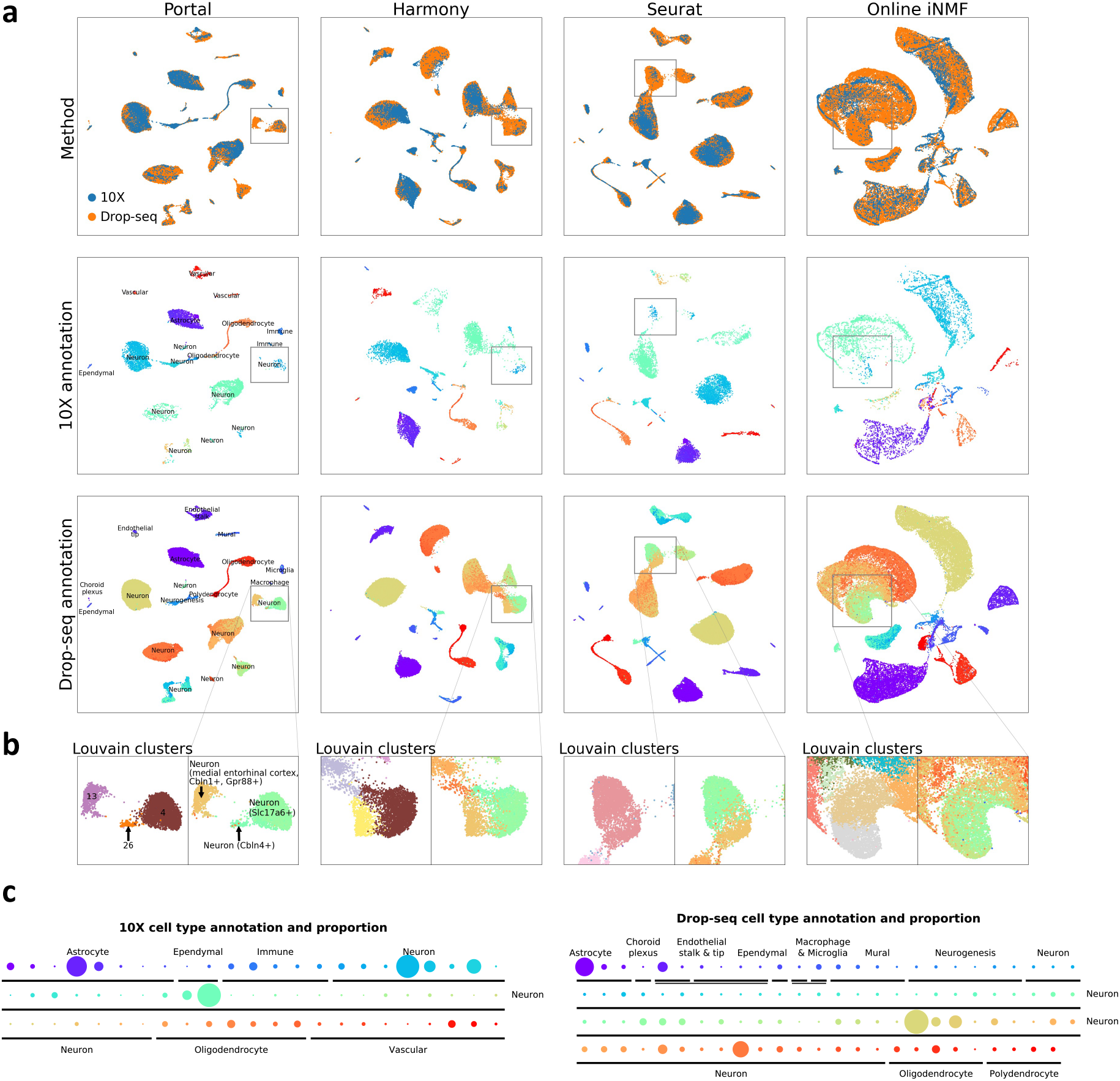
Preservation of fine-grained neuron subpopulations in the integration of hippocampus datasets. **a**. We visualized integration results from Portal, Harmony, Seurat and online iNMF of hippocampus datasets profiled by Drop-seq and 10X with UMAP [39]. Top panels are UMAP plots colored by profiling methods. Middle and bottom panels are UMAP plots of cells from the 10X dataset, the Drop-seq dataset after integration respectively, colored by fine-grained annotations (**c**). **b**. We marked a region containing three distinct neuron subpopulations. Results from Louvain clustering algorithm were presented for a comparison of cluster identity preservation performance. **c**. Cell type annotations and proportions of the two datasets from their original publications [8, 9]. The comparison among proportions of subpopulations was visualized by the sizes of corresponding dots.

After applying Portal and the other three benchmarked methods to integrate the data, we used the integrated cell representations to perform clustering. Using the Louvain method [40] with default resolution, we obtained 29 (Portal), 29 (Harmony), 25 (Seurat) and 30 (online iNMF) clusters, respectively (Fig. S19). Particularly, we focused on one region where the cell proportions between two datasets were highly unbalanced, as marked in Fig. 3**a**. Only a few of cells in this region are from the 10X dataset, making it challenging to build alignment between datasets while preserving subpopulations from the Drop-seq dataset. In the original publication [8], cells from the Drop-seq dataset within the marked region were all annotated as neurons but further classified into three transcriptionally distinct subpopulations, namely: *Cbln1*+/*Grp88*+ medial entorhinal cortex neurons; *Slc17a6*+ neurons; and *Cbln4*+ neurons. Among the benchmarked methods, Portal was the only method that clearly clustered these cells into three coherent groups in the integrated embedding space. Specifically, clusters 4, 13, and 26 identified by the Louvain method recovered the *Slc17a6*+ neuron; *Cbln1*+/*Grp88*+ medial entorhinal cortex neuron; and the *Cbln4*+ neuron subpopulations, respectively (Fig. 3**b**). Each cluster was confirmed by the high expression level of the annotated marker genes (Fig. S20**a**). Notably, these three groups only accounted for 4.79%, 1.76% and 0.32% of the total sample size, respectively, demonstrating Portal’s ability to preserve identities of rare subpopulations. However, the differences among these three subpopulations were not well preserved by the other three methods, making it difficult to detect them each distinctly using the Louvain clustering method (Fig. 3**a, b**). As shown in Fig. S20**c**, we also identified eight protein coding genes that were the most significantly differentially expressed among clusters, indicating the different functions of each of the three neuron subtypes. Cluster 4 showed high expression levels of *Camk2n1, Map1b, Nrgn, Syt1*, and no detectable expression of *Camk2d, Igfbp5, Nr4a2* and *Ntng1*. A different pattern was observed in cluster 13: high expression of *Camk2d, Camk2n1, Map1b* and *Syt1*, and no detectable expression of the other four genes. Cluster 26, meanwhile, showed moderate levels of expression of all eight genes. In the marked region, cells from the 10X dataset were mostly concentrated in clusters 4 and 13. The alignment by Portal was confirmed by the consistent gene expression levels in clusters 4 and 13 between the two datasets (Fig. S20**b**). Besides the eight differentially expressed genes, we also examined a larger set of genes, and computed the cross correlation of these genes pairwise between cells from all three groups. This analysis showed that cells within each cluster had higher similarity in gene expression than cells from other clusters, further showing the biological difference between these three clusters that should not be mixed after integration (Fig. S20**d**). The above results highlight Portal’s power to preserve rare cell types.

The integrative analysis on the hippocampus tissue demonstrates Portal’s ability to maintain nuanced transcriptional differences for small subpopulations. This means that Portal can also be used to “call out” rare subpopulations in one dataset based on integration with another dataset via label transfer. To illustrate this feature, we take 10X and SMART-seq2 (SS2) data generated for a mouse lung scRNA-seq atlas [7] as an example: the typically larger sample size of the 10X dataset facilitates powerful clustering analyses for identification of cell types; while the greater sequencing depth and sensitivity of SS2 enables deeper investigation into cell biology [41]. To leverage the different strengths of the two technologies, we used Portal to perform integrated analysis on 1,676 SS2 cells and 5,404 10X cells (Fig. S21**a**). Specifically, we defined the 10X dataset annotations from the original publication [7] as reference labels (Fig. S21**b**), then made use of the Portal’s integration results to identify cell types for the SS2 dataset based on these reference labels. After integration, for each SS2 cell, label transfer was performed by detecting its nearest neighbors among 10X cells. From this analysis, we identified four subpopulations of myeloid cells for the SS2 dataset, namely alveolar macrophages, dendritic cell and interstitial macrophages, classical monocytes, and non-classical monocytes (Fig. S21**d**). Transferred labels of these four subpopulations were validated by known marker gene expression levels [42]. For example, compared to classical monocytes, non-classical monocytes showed lower expression of *Ccr2* and higher expressions of *Treml4* (Fig. S22). Consistent with the gene expression pattern of alveolar macrophages in the 10X dataset, alveolar macrophages annotated by Portal in the SS2 dataset had high expression levels of marker genes *Mrc1* and *Siglec5*. Notably, in the SS2 dataset, the alveolar macrophage subpopulation only accounted for 0.78% of total sample size, and could not be distinguished from the other SS2-profiled macrophages in the original publication [7]. Based on the original labels, alveolar macrophages were unidentified as they were labeled in a more general group named “dendritic cell, alveolar macrophage, and interstitial macrophage” (Fig. S21**c**). Making good use of the larger 10X dataset, Portal successfully identified extremely rare subpopulations within the SS2 dataset. We then used the mouse lemur bladder scRNA-seq datasets from the Tabula Microcebus Consortium [43] as another example to demonstrate Portal’s ability for discovering rare subpopulations via label transfer. In this example, mouse lemur bladder tissue was also profiled by both SS2 and 10X. When we integrated these datasets and transferred labels from the 10X dataset to the SS2 dataset using Portal, we were able to distinguish a very small myofibroblast subpopulation of just 11 cells in the SS2 dataset from the rest of the fibroblasts (Fig. S23**a**). We verified their myofibroblast identity based on their high expressions of known marker genes *ACTA2, MYH11, TAGLN* [44] (Fig. S23**b**).

### Integration of comprehensive whole-organism cell atlases

So far, Portal has shown impressive performance in aligning tissue-level atlases where nuanced transcriptional differences among subpopulations can be maintained after integration. We next assess Portal’s capabilities under another challenging scenario: integrating two atlases across an entire organism, where one of the atlases includes many more organs and tissue types than the other. This can be very challenging for some integration algorithms due to having “missing cell types” in one of the datasets [10]. In contrast to these approaches, Portal uses discriminators with tailored design in the adversarial domain translation framework to distinguish domain-specific cell types from cell types shared across domains automatically, and is thus robust to non-overlapped tissue samples.

To build a foundation for extensive study of cell populations across the whole organism, the Tabula Muris Consortium [7] profiled cells from 20 tissues using a combination of SS2 (44,779 cells) and 10X (54,865 cells) (Fig. 4). Notably, seven of these 20 tissues were only profiled by SS2 but not 10X: brain (myeloid and non-myeloid), diaphragm, fat, large intestine, pancreas and skin. We used Portal to build a comprehensive integrated mouse atlas that merges all the cells, and we found Portal to show extraordinary accuracy in aligning cells of the same cell type from the two datasets profiled by different platforms, not only in the shared latent space but also in both domains (Figs. 4**a, b** and S24). After Portal integration, tissue-specific cell types of SS2-only tissues, such as microglial cells in brain (myeloid), cell types in large intestine, and pancreatic islets cells, were all successfully and correctly remained separated from other cell types. The other three benchmarked methods, however, failed to retain many tissue-specific cell types unmixed with other cell types. For instance, they mixed microglial cells together with other macrophage cells, even though the data from these two cell types were clearly transcriptionally different (Figs. 4**e** and S24).

**Figure 4:**
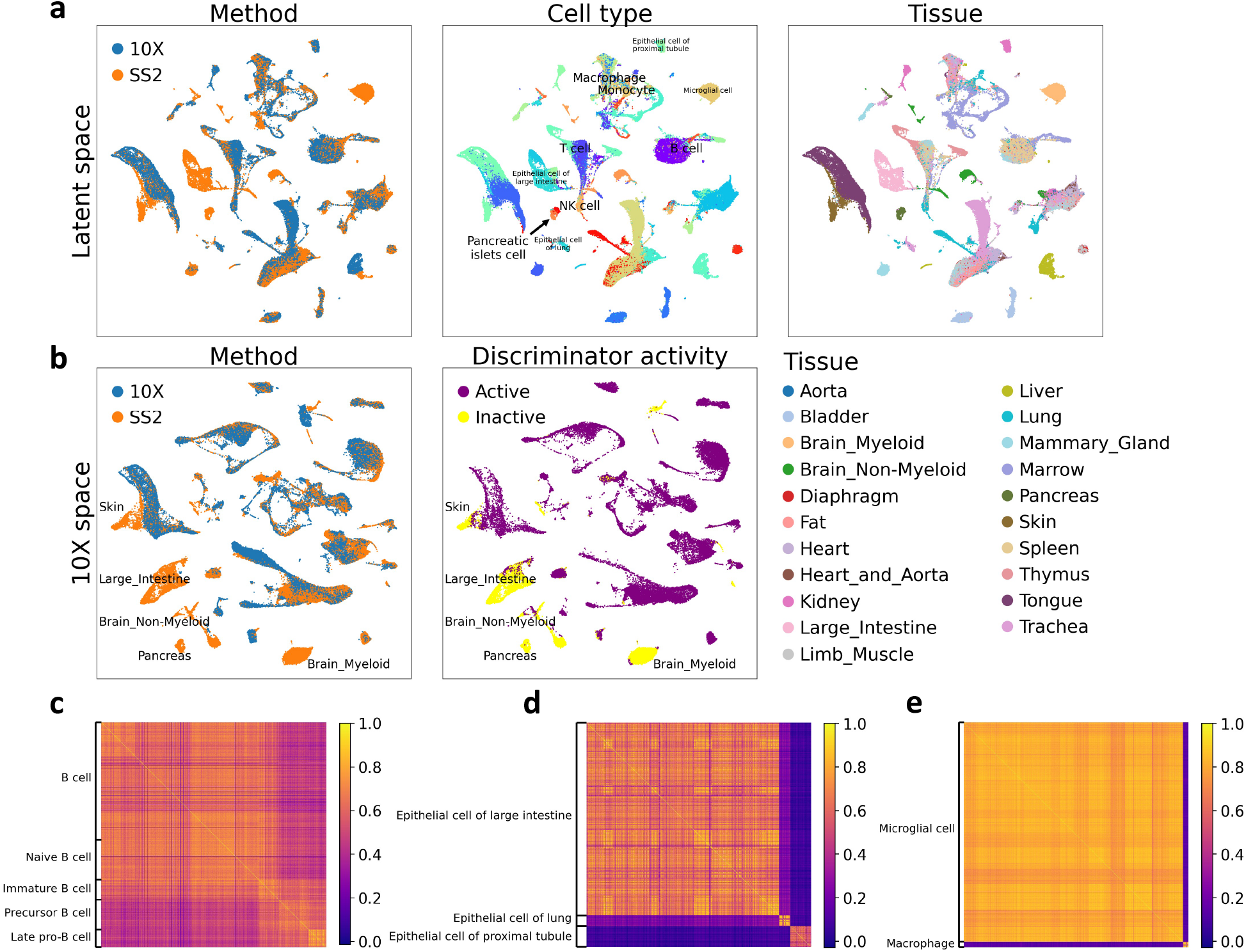
Construction of mouse cell atlas across the entire organism by integrating atlas datasets from the Tabula Muris project. We applied Portal to integrate the datasets obtained by 10X and SS2. There were cells from unique tissues presented in the SS2 dataset. **a**. UMAP plots of Portal’s integration results in the shared latent space, colored by profiling methods, cell types and tissues. **b**. Portal also transferred cells from the space of SS2 dataset to the space of the 10X dataset (10X space). In the 10X space, 10X cells were fixed as reference. Portal only aligned SS2 cells of shared cell types between datasets to 10X cells, while maintaining the identities of SS2 cells belonging to tissue-unique cell types. This was achieved by the special design of discriminator activity in Portal. **c, d**. Correlations among cells from subpopulations of B cells (**c**) and epthelial cells (**d**). **e**. Transcriptional distinction between macrophage and microglial cells.

Using this construction of a mouse cell atlas across organs, we also confirmed that the designed boundaries for discriminator active region in Portal (Fig. 1**c**) indeed helped to maintain the biological variation. By looking into the domain of 10X data (10X space), the discriminator in the 10X domain was found inactive for tissue-specific cell types that were only in the SS2 dataset (Fig. 4**b**). For these cells, Portal ensured that their identities were preserved by making the adversarial learning objective inactive on them automatically. Portal’s ability to conserve information of cell populations indicates its reliability for integrating atlas-level single-cell datasets across entire organisms.

Besides the alignment between datasets, Portal’s integration result could characterize the similarities and differences among cell types. For example, immune cells such as B cells, T cells, natural killer cells (NK cells), monocytes and macrophages were profiled by both platforms and contained in multiple tissues including brain (myeloid), diaphragm, fat, kidney, limb muscle, liver, lung, mammary gland, marrow, spleen, and thymus. Portal correctly kept the subpopulations belonging to the same type of immune cells close to each other, revealing the resemblance of immune cells across different tissues. For instance, the transcriptional correlation of all types of B cells, containing B cells, naive B cells, immature B cells, precursor B cells, and late pro-B cells confirmed such similarity (Fig. 4**c**). In addition, the epithelial cells of different tissues were identified by Portal as disjoint clusters, which was consistent with the biological distinction among these cell types (Fig. 4**d**).

### Portal successfully and efficiently aligns datasets across different data types

As most of existing methods were developed only for integrating scRNA-seq datasets, aligning datasets with different data types could be problematic for these approaches. Here we illustrate that Portal can flexibly account for the distinction between different data types and yield accurate integration results.

We first examined integration of scRNA-seq data and snRNA-seq data. For frozen samples such as biobanked tissues, and for tissue types that have unique morphology or phenotypes, such as brain, fat, or bone, it can be challenging or sometimes even impossible to extract intact cells for scRNA-seq profiling [45, 46]. To bypass this issue, snRNA-seq has been developed. Although nuclear transcriptomes are shown to be representative of the whole cell [47], distinctions between the whole cell and nucleus in terms of the transcript type and composition make scRNA-seq data and snRNA-seq data intrinsically different [45]. Aligning these two types of data is desirable, as the combined dataset enables joint analysis that can take advantages of both techniques, and help to improve statistical power for the analysis. Especially for comparing multiple complex tissues, with some cell types being shared and others being non-overlapping, researchers could benefit from such integrated joint analysis – one example being the integration of brain snRNA-seq data with scRNA-seq data of blood to examine similarities and differences between immune cells in each tissue milieu. However, due to the inherent difference in these two data types, aligning scRNA-seq and snRNA-seq data is not the same as batch effects correction. Compared to batch effects among scRNA-seq datasets, technical noise and unwanted variation arising from different data types are often more complex and have higher strength [45, 48]. Thus, using standard batch effects correction to integrate across data types may result in loss of alignment accuracy or important biological signals.

We evaluated Portal’s ability to integrate snRNA-seq data and scRNA-seq data using three mouse brain atlas datasets, including one snRNA-seq dataset profiled by SPLiT-seq [49], and two scRNA-seq datasets profiled by Drop-seq and 10X [8, 9]. In this task, we applied integration methods to harmonize these three atlases across all brain regions. To test the accuracy of integration results, we only used cells that had annotations provided by the authors in each atlas project. After selecting cells with cell type annotations, 319,359 cells in the Drop-seq dataset, 160,678 cells in the 10X dataset, and 74,159 nuclei in the SPLiT-seq remained for integration.

Prior to any integration, the raw datasets were clustered by the experimental methods rather than the cell types (Fig. S25**a**), and shared cell types between the three datasets did not align well, indicating the initial discrepancy between the three large datasets. After integration, UMAP visualizations showed that the different alignment methods gave varying results. Portal (Fig. S25**b**) and Seurat (Fig. S25**d**) achieved the best alignment of data among different methods, showing good mixing of cells annotated with the same cell type label, while also preserving subcluster data structure in the integrated results. In particular, the alignment of scRNA-seq (10X, Drop-seq) and snRNA-seq (SPLiT-seq) datasets was comparably good as that of the two scRNA-seq datasets, indicating successful alignment between the two data types without loss of biologically important variations between clusters. Online iNMF (Fig. S25**e**), although it successfully clustered and aligned the same cell types together, within each cluster the streaky pattern suggested potential numerical artefacts in the integrated data. Furthermore, online iNMF alignment resulted in loss of biological variation, which was most easily observable in the coalescence of the previously distinct neuron subpopulations (Fig. S25**a**) into one large amorphous cluster (Fig. S25**e**). Harmony, however, showed poor mixing of the snRNA-seq data in some of the cell types, such as the astrocytes, where the scRNA-seq datasets were well-mixed after alignment, but the snRNA-seq data were not mixed well with the rest (Fig. S25**c**). Similar to online iNMF, some of the neurons’ subcluster structure appeared to be lost after the integration by Harmony. Overall, Portal and Seurat presented the best scRNA-seq and snRNA-seq data alignment performance; however, not including data preprocessing time, Seurat took over 17 hours to complete the task, while Portal only took 87 seconds (details of the procedure are included in the Methods section).

We further assessed Portal’s ability to integrate across data types when no cell type, or very few cell types are shared. In this scenario, we applied Portal to integrate one human PBMC scRNA-seq dataset [50] with two human brain snRNA-seq datasets [51, 52], respectively, as two examples. In the first example (Fig. S26), where no cell type was shared between datasets, Portal did not mixed any two populations of cells together, showing its robustness. More importantly, it embedded monocytes and dendritic cells from the PBMC dataset close to macrophages from the brain dataset, indicating the similarities among these immune cells across tissue types. In comparison, overcorrection was observed in results from other state-of-the-art methods. For example, Seurat mixed T cells from blood with excitatory neurons and inhibitory neurons from brain inappropriately (Fig. S26). The reliability of Portal was also demonstrated in the second example (Fig. S27). It correctly aligned cells of the only shared cell type (T cell) between datasets, while it did not mix other distinct cell types (Fig. S27).

Besides integration of scRNA-seq data and snRNA-seq data, we then applied Portal to align scRNA-seq data and scATAC-seq data. As an epigenomic profiling method, scATAC-seq measures chromatin accessibility, providing a complementary view to scRNA-seq. Integrative analyses of scRNA-seq and scATAC-seq data are very helpful to leverage and unify information from the both aspects [53, 22]. For this task, we used one scRNA-seq PBMC dataset profiled by CITE-seq and one scATAC-seq PBMC dataset profiled by ASAP-seq [54]. For a better evaluation, we compared Portal with Seurat, online iNMF and VIPCCA, which had shown their ability of cross-omics integration in the original publications. A recent state-of-the-art method, scJoint [55], was also included in the comparison, as it was designed specifically for scRNA-seq and scATAC-seq data alignment.

As shown in the UMAP visualizations (Fig. 5), Portal, scJoint and VIPCCA were able to align the two datasets correctly, while online iNMF and Seurat did not align some cell clusters: for example, monocytes in online iNMF’s integration, and a cluster of mixed cell types from ASAP-seq in Seurat’s result. Among the benchmarked methods, Portal showed superior performance on the preservation of biological signals. After Portal’s integration, B cells and T cells were kept as disjoint clusters, while subpopulations of T cells were remained to be close to each other. In comparison, the coalescence of the previously distinct cell type clusters in VIPCCA’s result indicates the loss of information. Unlike other methods, scJoint requires cell type label information of scRNA-seq datasets as its input. It utilizes the cell type annotations to construct embedding of cells. As a result, cells from the scRNA-seq dataset with different cell type labels are forced to form disjoint clusters. Biological information was largely lost in scJoint’s integration of PBMC data: the subpopulations of T cells (naive CD4+ T cells, naive CD8+ T cells, effector CD4+ T cells, effector CD8+ T cells) lost their similarity and became far apart from each other (Fig. 5). Portal and scJoint were also benchmarked with a more challenging task: we manually removed B cells from the CITE-seq dataset such that B cells became a dataset-specific population. The results further demonstrated Portal’s robustness to unbalanced cell type compositions even in cross-omics integration, while scJoint showed comparatively inferior performance (Fig. S28).

**Figure 5:**
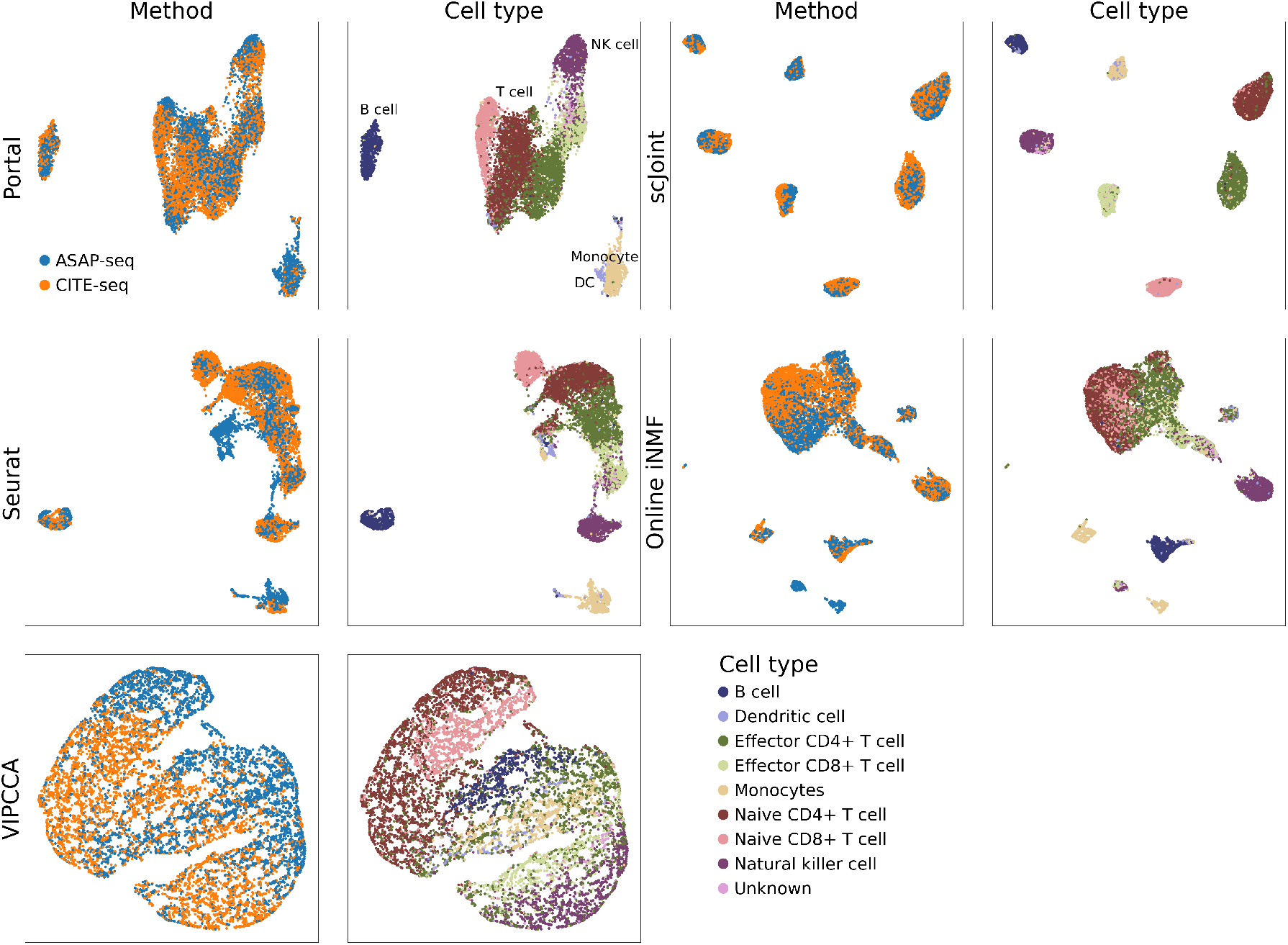
Comparison of Portal and other cross-omics integration methods on the alignment of scRNA-seq and scATAC-seq data. We applied Portal, scJoint, Seurat, online iNMF, and VIPCCA to align the scRNA-seq dataset (profiled by CITE-seq) and the scATACseq dataset (profiled by ASAP-seq) of peripheral blood mononuclear cells (PBMCs) [54]. UMAP plots were colored by profiling methods and cell types, respectively.

### Portal aligns spermatogenesis differentiation process across multiple species

Portal does not need to specify the structure and the strength of unwanted variation when integrating datasets. Instead, it can flexibly account for general difference between datasets, including batch effects, technical noises, and other sources of unwanted variation, by nonlinear encoders and generators in the adversarial domain translation framework. Therefore, Portal is also applicable for merging datasets with intrinsic biological divergence, revealing biologically meaningful connections among these datasets. In this section, we demonstrate that Portal can successfully align scRNA-seq datasets of the testes from different species including mouse, macaque and human (Fig. 6).

**Figure 6:**
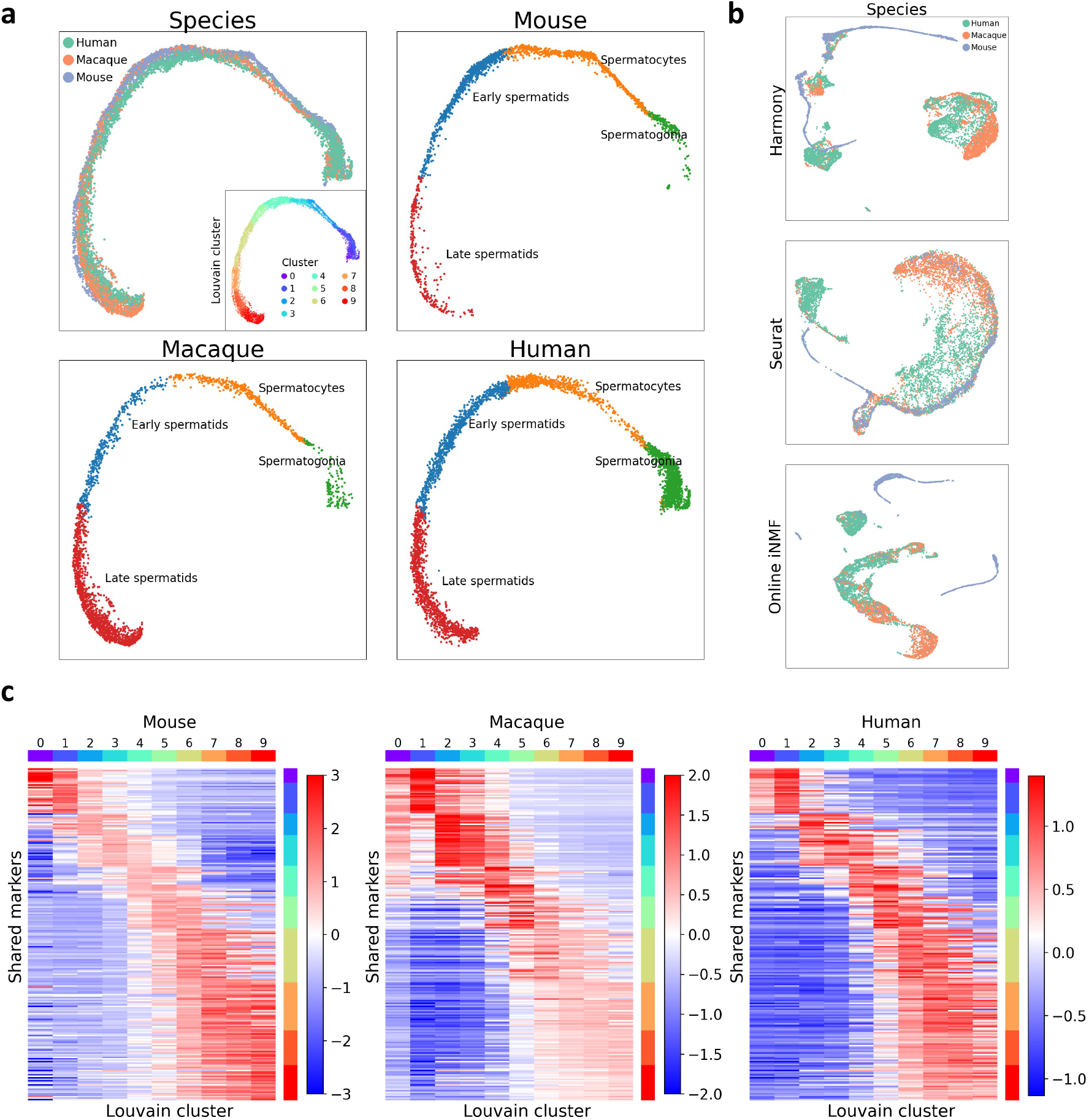
Integration of spermatogenesis datasets across different species, including mouse, macaque and human. **a**. The UMAP plot of Portal’s result colored by species, as well as UMAP plots of integrated mouse, macaque, human datasets visualized separately. Ten clusters were obtained by applying the Louvain clustering algorithm, facilitating detailed comparative analysis across species. **b**. Integration results of Harmony, Seurat and online iNMF. **c**. Portal identified 228 highly variable genes that are shared in the spermatogenesis process across all three mammalian species.

Compared to merging datasets from the same species, cross-species integration poses additional unique challenges. Although the transcriptomes of different species may share expression of homologous or orthologous genes, the number of shared genes varies between different species and is limited. Furthermore, two species may have genes with very similar sequence and be annotated in the transcriptome by the same name, but have altered function, which means that expression of the same gene in different species can denote different cell function [56]. In other words, the amount of information one can utilize for integration becomes limited and fuzzier while the variation across datasets becomes far larger, with limited number of shared genes and even fewer shared highly variable genes across different species. Nonetheless, cross-species integration can be very meaningful despite its challenges, as it can generate quick draft annotations of new or less-studied species’ atlases and cell types via label transfer from well-studied species. This saves time in the manual annotation process of single-cell tissue atlas generation for new species. Such integration can also enable detailed comparisons between species, such as comparisons of cell type composition, discovery of cell types unique to a particular species, or cross-species comparisons of the same cell types.

Mammalian spermatogenesis is a continuous and irreversible differentiation process from spermatogonial stem cells (SSCs) to sperm cells [57, 58, 59, 60, 16]. Due to the unique degenerate nature of the Y chromosome (Y-chr), Y-chr gene expression is intricately and tightly regulated in the spermatogenesis process through meiotic sex chromosome inactivation (MSCI) [61, 62, 63, 64, 65]. Interestingly, Y-linked genes are highly divergent between different species, including between closely related primates such as the chimpanzee, macaque, and human [61, 66, 67]; yet MSCI as a process is conserved across many species and is required for male fertility [64, 68]. This evidence suggests that while the evolution of genes on the Y-chr generated diverse species-specific genetic combinations, the tight control of gene expression through MSCI is required to ensure genetic stability [61]. Recently, cross-species comparisons of “escape genes” that are able to maintain or re-activate their expression despite MSCI repression during spermatogenesis have generated fascinating insights on evolutionary biology, and on sex chromosome evolution [63, 65, 69, 16]. In this biological context, integrating datasets with continuous and gradient developmental trajectories, such as for spermatogenesis data, requires integration methods to preserve the continuous structure of each dataset, while still providing high accuracy of cell type alignment between datasets. This is more difficult when, like in spermatogenesis data, there are no distinct clusters, making integration of such data a particularly difficult task. After confirming Portal’s capability of preserving the gradual change of cells based on two examples (Figs. S29 and S30), we perform cross-species integration of testes datasets from three species, including one mouse [59], one macaque and one human [16], aligning the different stages of spermatogenesis across species thereby highlighting unique features of each. The successful integration of these spermatogenesis trajectories serves as a demonstration of the power of Portal in complex and low-information data alignment, and how it can facilitate the annotation and discovery process for new single-cell tissue atlases.

We first annotated the mouse sample according to the pattern of marker genes (Spermatogonia: *Sycp1, Uchl1, Crabp1, Stra8*; Spermatocytes: *Piwil1, Pttg1, Insl6, Spag6*; Early spermatids: *Tssk1, Acrv1, Spaca1, Tsga8*; Late spermatids: *Prm1, Prm2, Tnp1, Tnp2*) [57, 58]. Then we used Portal to harmonize the three samples, where the integration was accomplished in the mouse sample domain: The cells from the mouse sample were used as reference, and cells from the other two species were mapped to the mouse sample domain by Portal. Based on our annotation of the mouse sample, we transferred the broad cell type labels to cells from the macaque and human samples according to the nearest neighbors, using the alignment given by Portal (Fig. 6**a**). To check whether the alignments were correct for broad cell type identities, we visualized the UMAPs for cells from each species labeled by their original published annotations [16], and we confirmed concordant cell type integration across species (Fig. S31). Then, we used Louvain clustering algorithm to cluster the cells from all three species based on integrated cell representations. Ten clusters were found, and the cluster names were relabeled by their order of progression from the spermatogonia along the developmental trajectory (Fig. 6**a**). We then visualized the expression of known spermatogenesis markers [57, 58, 16] in each Louvain cluster and found that the Louvain clusters generated by Portal’s alignment clearly captured the key transcriptomic features for each stage of spermatogenesis, and correctly identified cells from each stage for all three species (Fig. S32, S33). Furthermore, each Louvain cluster represented a more fine-grained classification of cells within the labeled broad spermatogenesis cell types. Using these clusters we assessed the transcriptomic changes throughout the differentiation trajectory with higher resolution (Fig. S32, S33). Notably, many of the marker genes known to define stages of spermatogenesis in human were not shared or sometimes not expressed in macaque and/or mouse scRNA-seq data. For example, human genes *SYCP3, YBX2, SPACA4, H1FNT, PRM1*, and *TNP1* were known to mark human spermatogenesis progression, but they were absent in the macaque dataset. As only highly variable genes that were expressed in all three species were considered in the integration process, these genes were not used by Portal. However, they showed clear expression in the cell clusters where they were expected to be expressed after integration (Fig. S33), confirming the correctness of Portal’s integration result. The above results show that Portal can provide an accurate integration even for genes not measured by all three samples. As a comparison, Harmony, Seurat and online iNMF were also applied. However, Harmony and online iNMF were unable to maintain the gradient developmental trajectories of spermatogenesis process for at least one species. All of the three methods showed less satisfactory ability to align cells across the three species (Fig. 6**b**).

Cross-species data integration can be a quick and easy way to generate draft cell atlas annotations for new species via label transfer from well-annotated species, but moreover, such integrated data can be used to highlight interesting biological features of shared cell types. In our Louvain clusters for spermatogenesis, for each species, we selected top 200 highly expressed genes of every cluster. By taking the intersection of those genes across three species, we then identified 228 highly variable genes that are shared in the spermatogenesis process across all three mammalian species (Fig. 6**c**). For the highly expressed genes that were unique to only one species, we compared their expressions across all three species (Fig. S34). Such comparisons could give insight into shared and divergent features of spermatogenesis across different species.

## Discussion

Taking advantage of machine learning methodologies, Portal is an efficient and powerful tool for single-cell data integration that easily scales to handle large datasets with sample sizes in the millions. As a machine learning-based model, Portal is easy to train, and its training process is greatly accelerated by using GPUs. Meanwhile, mini-batch optimization allows Portal to be trained with a low memory usage. Besides, it also makes Portal applicable in the situation where the dataset is not fully observed, but arrives incrementally.

The nonlinearity of neural networks makes Portal a flexible approach that can adjust for complex dataset-specific effects. Nonetheless, according to benchmarking studies, strong ability for removing dataset-specific effects often comes with the weakness in conserving biological variation [48, 30], e.g., being prone to overcorrection. Portal overcomes this challenge by its model and algorithm designs. First, the boundaries of discriminator scores help Portal to protect dataset-unique cell types from overcorrection. Second, the use of three specifically designed regularizers not only assists Portal to find correct correspondence across domains, but also enables Portal to have high-level preservation of subcluster and small cluster identities in both datasets.

Two existing popular methods are Seurat and BBKNN. Seurat often provides integration results with high accuracy, but also requires high computational cost, preventing its usage on large-scale datasets; while BBKNN is well-known for its extremely fast speed, its comparatively less precise results are sometimes a concern for users (Figs. S5 - S17). A major advance of Portal over these existing state-of-the-art integration approaches is its ability to achieve high efficiency and accuracy simultaneously. With speed comparable or faster than BBKNN, and significantly lower memory requirement than BBKNN (Fig. S18), Portal presents similar batch correction performance as well as superior information preservation performance compared to that of Seurat (Figs. S5 - S17).

Portal also has advantages over several existing deep learning-based methods for single-cell data integration. Currently, the majority of deep learning-based methods leverages the variational autoencoders (VAEs) framework [70]. scVI [25], as a prominent representative of VAE-based methods, is scalable to atlas-level datasets. It utilizes the zero-inflated negative binomial (ZINB) distribution in its modeling, which may be less efficient in capturing complex data structures [24]. scANVI [71] is another VAE-based method with similar pros and cons of scVI, as it is an extension of scVI that incorporates cell type information into its model. Recently, VIPCCA [24] was proposed to leverage VAE-based networks to perform nonlinear canonical correlation analysis (CCA) efficiently. However, we empirically found that it favors the removal of batch effects over the preservation of biological information (Figs. S6, S12 and S16). scGen [72] utilizes a VAE to find a difference vector in the latent space of each cell type across batches. Similar to scANVI, scGen requires cell type information as its input. There are also some methods using strategies other than VAEs. One category is deep learning-based methods utilizing searched MNN pairs as the reference, and then using neural networks to correct batch effects, such as iMAP [73] and deepMNN [74]. Consequently, the second stage of correcting batch effects heavily relies on the first stage of constructing MNN pairs. Moreover, searching MNN pairs is usually performed on CPUs and could be less computationally efficient for larger datasets. Some deep learning-based methods focus on integrating cross-omics datasets, including cross-modal autoencoder [75] and scJoint [55]. However, they require additional information like cell type information or paired data points for data alignment. They may not be applicable when such information is unavailable. Compared to existing deep learning-based methods, Portal neither relies on a parametric distribution for single-cell data, nor requires MNN pairs to serve as anchors for integration. Owing to its unified framework with unique designs for single-cell datasets, Portal enjoys high flexibility to handle complex datasets and dataset-specific effects with varying strength, and high scalability to deal with millions of cells efficiently.

By leveraging the adversarial domain translation framework, Portal can build meaningful alignment between datasets with efficient utilization of information. From single tissue types to complex cell atlases, Portal showed extraordinary information preservation performance throughout all integration tasks. This feature of Portal is exemplified by integration of the spermatogenesis trajectory across three species, where only a limited number of highly variable genes were shared and utilized by Portal. Improvements can further be made if an effective way of leveraging the whole transcriptome of all species is developed, which is left for future work to address. Nonetheless, such cross-species integration allows biologists to easily identify shared and divergent cellular programs across different species, which is particularly useful for addressing questions of evolutionary biology. In our example of mouse, macaque, and human testes tissue integration, identifying genes that are primate-specific can help to generate hypotheses about the evolution of primates and shed light on the applicability of various animal models for biological research.

It is now clear that using single-cell technologies to assemble comprehensive whole organism atlases encompassing diverse cell types is accelerating biological discovery, and this demand will only grow as more datasets are generated. The demand for integration of such datasets, along with the size of these datasets, will expand correspondingly. We expect that Portal, with its fast, versatile, and robust integration performance, will play a valuable and essential role in the modern life scientist’s single-cell analysis.

## Methods

### The model of Portal

Expression measurements of cells from two different studies are viewed as datasets originated from two different domains 𝒳 and 𝒴. After standard data preprocessing of the expression data, Portal performs joint principle component analysis (PCA) across datasets and adopts the first *p* principal components of cells as the low-dimensional representation of cells, namely, cell embeddings. Portal takes the cell embeddings as the input to achieve data alignment between 𝒳 and 𝒴. To learn a harmonized representation of cells, Portal introduces a *q*-dimensional latent space Ƶ to connect 𝒳 and 𝒴, where the latent codes of cells in Ƶ are not affected by domain-specific effects but capture biological variation.

Portal achieves the integration of datasets through training a unified framework of adversarial domain translation. Let **x** and **y** be the cell embeddings in 𝒳 and 𝒴, respectively. For domain 𝒳, Portal first employs encoder *E*_1_(·) : 𝒳 → Ƶ to get a latent code *E*_1_(**x**) ∈ Ƶ for all **x** ∈ 𝒳. Encoder *E*_1_(·) is designed to remove domain-specific effects in 𝒳. To transfer cells from 𝒳 to 𝒴, Portal then uses generator *G*_2_(·) : Ƶ → 𝒴 to model the data generating process in domain 𝒴, where domain-specific effects in 𝒴 are induced. *E*_1_(·) and *G*_2_(·) together form a domain translation network *G*_2_(*E*_1_(·)) that maps cells from 𝒳 to 𝒴 along 𝒳 → Ƶ → 𝒴. By symmetry, encoder *E*_2_(·) : 𝒴 → Ƶ and generator *G*_1_(·) : Ƶ → 𝒳 are utilized to transfer cells from 𝒴 to 𝒳 along the path 𝒴 → Ƶ → 𝒳.

Portal trains domain translation network *G*_2_(*E*_1_(·)) : 𝒳 → 𝒴, such that the distribution of transferred cells *G*_2_(*E*_1_(**x**)) can be mixed with the distribution of cell embeddings **y** in domain 𝒴. Discriminator *D*_2_(·) is employed in domain 𝒴 to identify where the poor mixing of the two distributions occurs. The competition between domain translation network *G*_2_(*E*_1_(·)) and discriminator *D*_2_(·) is known as adversarial learning [31]. Discriminator *D*_2_(·) will send a feedback signal to improve the domain translation network *G*_2_(*E*_1_(·)) until the two distributions are well mixed. By symmetry, domain translation network *G*_1_(*E*_2_(·)) : 𝒴 → 𝒳 and discriminator *D*_1_(·) deployed in domain 𝒳 form another adversarial learning pair. The feedback signal from *D*_1_(·) improves *G*_1_(*E*_2_(·)) until the well mixing of the transferred cell distribution *G*_1_(*E*_2_(**y**)) and the original cell distribution **x** in domain 𝒳.

Notice that the well mixing of the transferred distribution and the original distribution does not necessarily imply the correct correspondence established between 𝒳 and 𝒴. First, cells from a unique cell population in domain 𝒳 should not be forced to mix with cells in domain 𝒴. Second, cell types *A* and *B* in domain 𝒳 could be incorrectly aligned with cell types *B* and *A* in domain 𝒴, respectively, even if the two distributions are well mixed. These problems can occur because we don’t have any cell type label information as an anchor for data alignment across domains. To address these, Portal has the following unique features, distinguishing it from existing domain translation methods [32, 33]. First, Portal has a tailored discriminator for the integrative analysis of single-cell data, which can prevent mixing of unique cell types in one domain with a different type of cell in another domain. Second, Portal deploys three regularizers to find correct correspondence during adversarial learning; these regularizers also play a critical role in accounting for domain-specific effects and retaining biological variation in the shared latent space Ƶ.

We propose to train domain translation networks under the following framework:

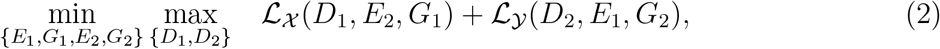

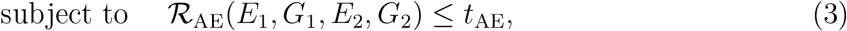

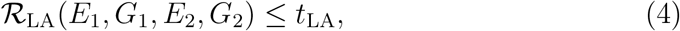

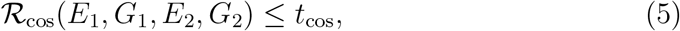

where component (2) is the objective function of adversarial learning for single-cell data integration; components (3), (4) and (5) are regularizers for imposing the autoencoder consistency, the latent alignment consistency and cosine similarity to preserve cross-domain correspondence, respectively. We have investigated the roles of each component in Portal and provided more results (Figs. S1 and S2) in the Supplementary Information. We explain each component in more detail in the next section.

#### Adversarial learning with discriminator score thresholding

The adversarial training between discriminators and domain translation networks is formulated as a min-max optimization problem (2), where ℒ_𝒳_ (*D*_1_, *E*_2_, *G*_1_) = 𝔼[log *D*_1_(**x**)] + 𝔼[log(1 − *D*_1_(*G*_1_(*E*_2_(**y**))))] and ℒ_𝒴_(*D*_2_, *E*_1_, *G*_2_) = 𝔼[log *D*_2_(**y**)] + 𝔼[log(1 − *D*_2_(*G*_2_(*E*_1_(**x**))))] are the objective functions for adversarial learning in domain 𝒳 and domain 𝒴, respectively. Given domain translation network *G*_1_(*E*_2_(·)), discriminator *D*_1_(·) : 𝒳 → (0, 1) is trained to distinguish the transferred cells *G*_1_(*E*_2_(**y**)) from the original cells **x**, where a high score (close to 1) indicates a “real cell” in domain 𝒳, and a low score (close to 0) indicates a “transferred cell” from domain 𝒴. This is achieved by maximizing ℒ_𝒳_ with respect to *D*_1_(·). Similarly, discriminator *D*_2_(·) in domain 𝒴 is updated by maximizing ℒ_𝒴_. Given discriminators *D*_1_(·) and *D*_2_(·), the domain translation networks are trained by minimizing ℒ_𝒳_ + ℒ_𝒴_ with respect to *E*_1_(·), *G*_2_(·) and *E*_2_(·), *G*_1_(·), such that the discriminators cannot distinguish transferred cells from real cells. This is equivalent to 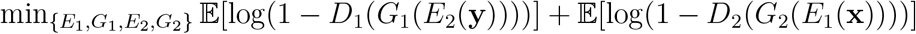. However, direct optimization of this objective function is known to suffer from severe gradient vanishing [31, 76]. Therefore, we adopt the “logD-trick” [31] to stabilize the training process. Denote 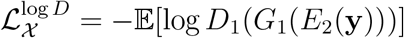 and 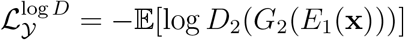. In practice, we minimize 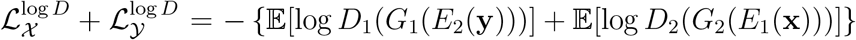 with respect to *E*_1_(·), *G*_2_(·) and *E*_2_(·), *G*_1_(·), instead of minimizing ℒ_𝒳_ + ℒ_𝒴_ = 𝔼[log(1 − *D*_1_(*G*_1_(*E*_2_(**y**))))] + 𝔼[log(1 − *D*_2_(*G*_2_(*E*_1_(**x**))))].

Although the above adversarial learning can make the transferred cells and real cells well mixed, it can falsely force cells of a unique cell population in one domain to mix with cells in another domain, leading to overcorrection. Consider a cell population that is present in 𝒳 but absent in 𝒴 as an example. On one hand, discriminator *D*_1_(·) can easily identify cells from the unique cell population as real cells in 𝒳. Cells in the nearby region of this cell population have extremely high discriminator scores. Some cells in 𝒴 will be mapped into this region by the domain translation network *G*_1_(*E*_2_(·)), leading to incorrect mixing of cell types in 𝒳. On the other hand, cells transferred from 𝒳-unique population will have low *D*_2_ scores in 𝒴. Discriminator *D*_2_(·) will incorrectly force the domain translation network *G*_2_(*E*_1_(·)) to mix these cells with real cells in domain 𝒴. The cell identity as a domain-unique population in 𝒳 is lost.

From the above reasoning, domain-unique cell populations are prone to be assigned with extreme discriminator scores, either too high in the original domain or too low in the transferred domain. Such extreme scores can lead to overcorrection. To address this issue in single-cell data integration tasks, we set boundaries for discriminator scores to make discriminators inactive on such cells. Specifically, the outputs of standard discriminators are transformed into (0, 1) with the sigmoid function, i.e., *D*_*i*_(**x**) = sigmoid(*d*_*i*_(**x**)) = 1*/*(1 + exp(−*d*_*i*_(**x**))), *i* = 1, 2, where *d*_*i*_(**x**) ∈ (−∞, ∞) is the logit of the output. We bound the discriminator score by thresholding its logit to a reasonable range [−*t, t*]:

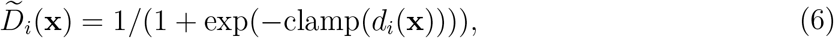

where clamp(·) = max(min(·, *t*), −*t*). By clamping the logit 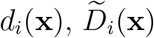 becomes a constant when *d*_*i*_(**x**) < −*t* or *d*_*i*_(**x**) > *t*, providing zero gradients for updating the parameters of encoders and generators. Meanwhile, 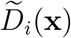 remains the same as *D*_*i*_(**x**) when *d*_*i*_(**x**) ∈ [−*t, t*]. By such design, the adversarial learning mechanism in Portal is only applied to cell populations that are likely to be common across domains. In Portal, we then use this modified version of discriminators 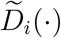 to avoid incorrect alignment of domain-unique cell populations. For clarity, we still use the notation *D*_*i*_(·) to represent 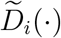 hereinafter.

#### Regularization for autoencoder consistency

Encoder *E*_1_(·) : 𝒳 → Ƶ and generator *G*_1_(·) : Ƶ → 𝒳 form an autoencoder structure, where *E*_1_(·) removes domain-specific effects in 𝒳, and *G*_1_(·) recovers them. Similarly, *E*_2_(·) : 𝒴 → Ƶ and *G*_2_(·) : Ƶ → 𝒴 form another autoencoder structure. Therefore, we use the regularizer in (3) for the autoencoder consistency, where 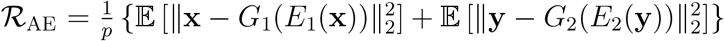, *p* is the dimensionality of 𝒳 and 𝒴.

#### Regularization for cosine similarity correspondence

Besides the autoencoder consistency, the cosine similarity regularizer in (5) plays a critical role in data alignment between domains, where 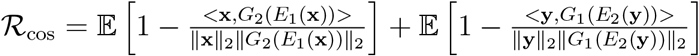 is the regularizer that imposes the cross-domain correspondence on domain translation. The key idea is that a cell and its transferred version should not be largely different from each other in terms of cosine similarity. This is because cosine similarity is scale invariant and insensitive to domain-specific effects, including differences in sequencing depth and capture efficiency of protocols used across datasets [77, 26, 21]. Thus, the cosine similarity regularizer is helpful to uncover robust correspondence between cells of the same cell type across domains.

#### Domain-specific effects removal in the shared latent space by latent alignment regularization

Portal decouples domain translation into the encoding process 𝒳 → Ƶ (or 𝒴 → Ƶ) and the generating process Ƶ → 𝒴 (or Ƶ → 𝒳). Although adversarial learning enables the domain translation networks to effectively transfer cells across domains, it can not remove domain-specific effects in shared latent space Ƶ. To enable encoders *E*_1_(·), *E*_2_(·) to eliminate domain-specific effects in 𝒳 and 𝒴, we propose the latent alignment regularizer in (4) for the consistency in latent space Ƶ, where 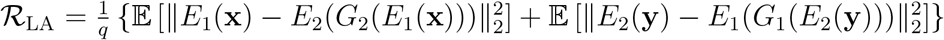, *q* is the dimensionality of Ƶ, *E*_1_(**x**) is the latent code of a real cell **x** ∈ 𝒳 and *E*_2_(*G*_2_(*E*_1_(**x**))) is the latent code of its transferred version, *E*_2_(**y**) is the latent code of a real cell **y** ∈ 𝒴 and *E*_1_(*G*_1_(*E*_2_(**y**))) is the latent code of its transferred version. The regularizer (4) encourages the latent codes of the same cell to be close to each other. This regularizer helps encoders *E*_1_(·) and *E*_2_(·) to remove domain-specific effects, such that the latent codes in Ƶ preserve biological variation of cells from different domains.

### Algorithm

Now we develop an alternative updating algorithm to solving the optimization problem of adversarial domain translation with the three regularizers. To efficiently solve the optimization problem, we replace the constraints (3), (4) and (5) by its Lagrange form. We introduce three regularization parameters *λ*_AE_, *λ*_LA_ and *λ*_cos_ as coefficients for the regularizers. The optimization problem of Portal is rewritten as

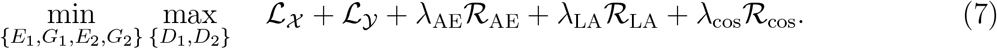

As we adopt the “logD-trick” for updating domain translation networks formed by *E*_1_(·), *G*_2_(·) and *E*_2_(·), *G*_1_(·), the optimization problem (7) is modified accordingly as

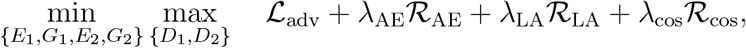

where ℒ_adv_ stands for the adversarial learning objective, whose value is ℒ_𝒳_ +ℒ_𝒴_ when maximizing with respect to *D*_1_(·), *D*_2_(·), and it is replaced with 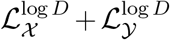 when minimizing with respect to *E*_1_(·), *G*_1_(·), *E*_2_(·), *G*_2_(·).

Let the parameters of the networks *E*_1_(·), *E*_2_(·), *G*_1_(·), *G*_2_(·), *D*_1_(·) and *D*_2_(·) be denoted as 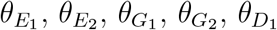 and 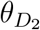 Then we collect the parameter sets as 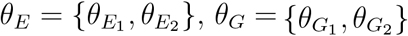 and 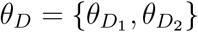 We use the Monte Carlo estimators to approximate expectations in Portal’s objective. With a mini-batch of 2*m* samples including {**x**^(1)^, **x**^(2)^, · · ·, **x**^(*m*)^} from 𝒳 and {**y**^(1)^, **y**^(2)^, · · ·, **y**^(*m*)^} from 𝒴, the Monte Carlo estimators are given by

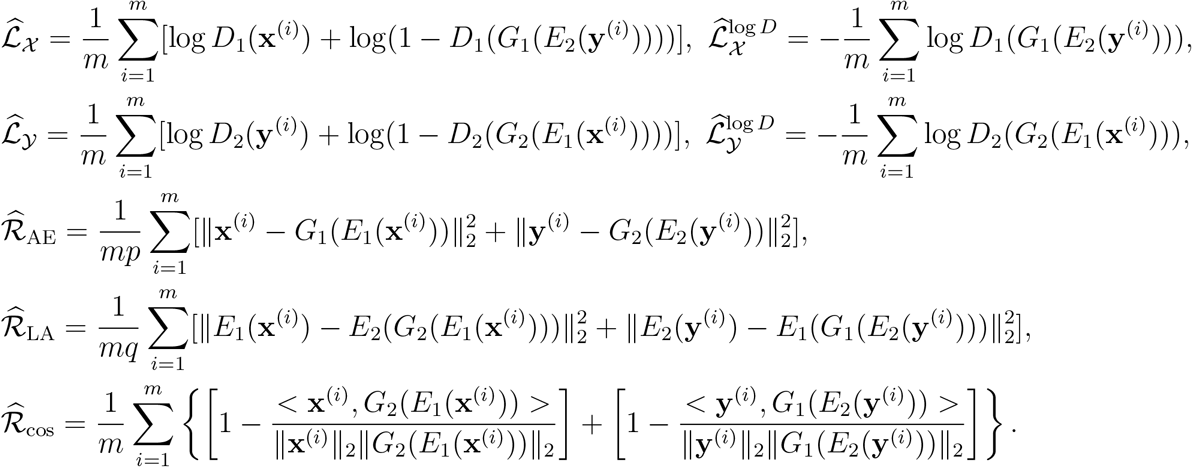

The implementation of Portal is summarized in Algorithm 1.

#### Algorithm 1

Stochastic gradient descent training of Portal.

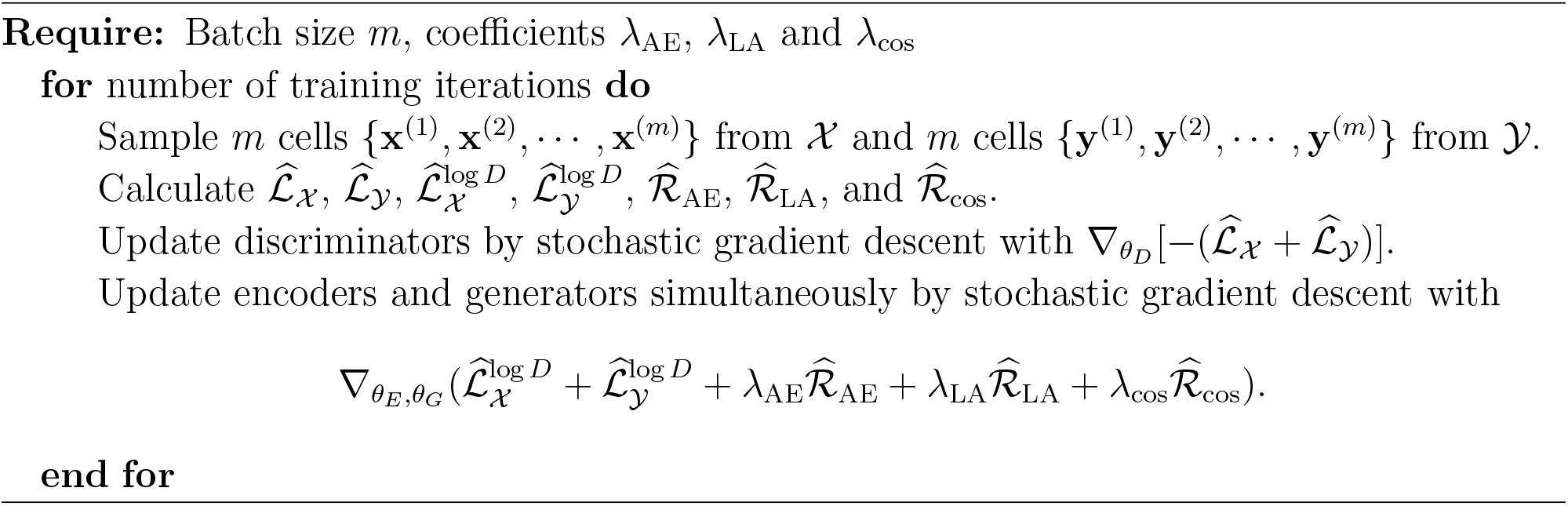

After training, cells from domains 𝒳 and 𝒴 are encoded into Ƶ to construct an integrated dataset, which can be applied to downstream analysis. In each domain, the original cells and transferred cells are also well integrated. For integration of multiple datasets, Portal can handle them incrementally, by transferring all other datasets into the domain formed by one dataset. **Network structure**. Portal uses lightweight networks which enable computationally efficient training when dealing with large-scale datasets. The details of Portal’s networks, including network structures, the number of layers and parameters are shown in the Supplementary Information (Tables S1, S2 and S3).

## Analysis details

### Data preprocessing

We used raw read or unique molecular identifier (UMI) matrices depending on the data source for all scRNA-seq and snRNA-seq datasets, and gene activity matrices for scATAC-seq datasets. We then performed standard data preprocessing for each count matrix, including log-normalization, feature selection, scaling and dimensionality reduction. For each dataset represented by a cell-by-gene count matrix, we first adopted the log-normalization, following the Seurat and Scanpy pipelines [22, 78]. For each cell, its library size was normalized to 10, 000 reads. Specifically, the counts abundance of each gene was divided by the total counts for each cell, then multiplied by a scaling factor of 10, 000. The normalized dataset was then transformed to log scale by the function log(1 + *x*). In order to identify a subset of features that highlight variability across individual cells, we adopted the feature selection procedure from the Seurat pipeline. For each dataset, we selected *K* top highly variable genes ranked by dispersion with the control of means. In this paper, we used *K* = 4, 000 throughout all analyses except for the cross-species analysis. In the cross-species analysis, we used *K* = 3, 000 since the usage of a larger number of features would result in the situation that correspondence across species is dominated by the distinction (e.g., altered functions of genes annotated by the same name). For each selected variable gene, we centered and standardized its expressions across individual cells to have mean at zero and variance at one. After the above procedures, which were applied to individual datasets, we continued to preprocess data across datasets. For those datasets to be integrated, we collected genes that were identified as top highly variable genes in all of them as features for integration. We extracted the scaled data with these features from each dataset, and then concatenated them based on features to perform joint PCA. Top *p* = 30 principle components were kept for all dataset as inputs to Portal. For the shared latent space, we set its dimensionality to be *q* = 20 throughout all analyses.

### Unifying gene names for cross-species integration

We retrieved pairwise orthologues (human vs mouse, human vs macaque) respectively from Ensembl Biomart, and merged them to obtain one-to-one-to-one orthologues by using human Ensembl gene names as reference. One-to-one-to-one orthologues across the three species were used to unify gene names. Genes included in the list were used by Portal. To facilitate the usage of Portal, we have included the used gene lists (orthologues human mouse.txt, orthologues human macaque.txt) as well as the reproducible code for the cross-species integration, among all details for reproducing the experiments throughout our paper at https://github.com/YangLabHKUST/Portal.

### Hyperparameter setting

Hyperparameters used in Portal are *m, t, λ*_AE_, *λ*_LA_, *λ*_cos_, where *m* is the batch size used by Portal for mini-batch training; *t* is the absolute value of boundaries for the logit of discriminator scores (−*t* < *d*_*i*_(**x**) < *t, i* = 1, 2); *λ*_AE_, *λ*_LA_, *λ*_cos_ are coefficients for autoencoder consistency regularizer ℛ_AE_, latent alignment regularizer ℛ_LA_ and cosine similarity regularizer ℛ_cos_ respectively. Throughout all analyses, we set *m* = 500, *t* = 5.0, *λ*_AE_ = 10.0, *λ*_LA_ = 10.0. Hyperparameter *λ*_cos_ was tuned within the range [10.0, 50.0] with interval 5.0 according to the mixing metric, where the mixing metric was designed in Seurat to evaluate how well the datasets mixed after integration. The insight into tuning *λ*_cos_ is as follows: During domain translations, there is a trade-off between preservation of similarity across domains and flexibility of modeling domain differences. Since ℛ_cos_ is designed to preserve the cosine similarity during translations, a higher value of *λ*_cos_ can enhance the cosine similarity as the cross-domain correspondence, and a lower *λ*_cos_ allows domain translation networks to deal with remarkable differences between domains. Following this intuition, we empirically find out that *λ*_cos_ = 10.0 has a good performance when harmonizing datasets with intrinsic differences, for example, datasets used in cross-species analysis or cross-modal integration (scRNA-seq and scATAC-seq). For other integration tasks, *λ*_cos_ = 20.0 often yields reasonable results, which is adopted as the default setting in our package. Slightly better alignment results could be achieved by tuning *λ*_cos_. Through a parameter sensitivity analysis, we have shown that Portal’s performance is insensitive to the choice of hyperparameters (Figs. S3, S4 in the Supplementary Information).

### Label transfer

Suppose we wish to transfer labels from domain 𝒳 to domain 𝒴. As Portal produces integrated cell representations in each domain and the shared latent space, we can use any of these representations to perform label transfer. For each cell in domain 𝒴, we find its *k* = 20-nearest neighbors among the cells in domain 𝒳 based on the integrated result. The metric for finding nearest neighbors can be Euclidean distance in shared latent space, or cosine similarity in domains. The labels in domain 𝒴 are finally determined by majority voting.

### Evaluation metrics

We assessed all metrics based on Portal’s integration results in shared latent space Ƶ. We used kBET [35], PCR batch [35], batch ASW [35], graph iLISI [30, 21] and graph connectivity [30] to assess the ability of batch correction. We used ARI [36], NMI [37], cell type ASW, graph cLISI [30, 21], isolated label F1 [30], isolated label silhouette [30] and cell cycle conservation [30] to evaluate the conservation of biological variation. The metrics, if necessary, were rescaled to [0, 1] such that a higher value represents a better performance.

#### kBET

For each selected cell, kBET adopts a Pearson’s *χ*^2^-based test to check whether the batch label distribution in its neighbourhood is similar to the global batch label distribution or not. In our experiments, we ran 100 replicates of kBET with 1,000 random samples, and used the median of average acceptance rates as the output. The neighbourhood size was chosen following the default setting in kBET’s official code.

#### PCR batch

PCR batch quantifies the removal of batch effects by comparing the variance contributions of the batch effects to datasets before integration (*V C*_before_) and after integration (*V C*_after_), respectively. In our experiments, we concatenated datasets by batches to obtain the dataset before integration. PCR batch score was calculated as 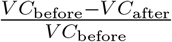. We clamped PCR batch score to [0, 1], where a higher score means that the impact of batch effects is eliminated after integration.

#### Batch ASW

Batch ASW calculates the silhouette width of cells with respect to batch labels. If batch effects are corrected in cell embeddings, the evaluated ASW (with respect to batches) should be close to -1, indicating the good mixing of cells across batches. We rescaled the score by 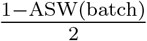.

#### Graph iLISI

The original iLISI is defined as the effective number of datasets in a neighborhood, where 1 means poor mixing, and 2 indicates good mixing of two datasets. Graph iLISI extends iLISI by enabling the calculation on graphs. The values were rescaled to [0, 1] by subtracting 1.

#### Graph connectivity

Graph connectivity assesses whether the graph correctly connects cells of same cell type labels among batches. We used the Scanpy pipeline to derive graph representation of integrated cell embeddings. The neighborhood size was set to be 15 (default setting in Scanpy).

#### ARI

ARI measures the degree to which the two clustering results match. It ranges from 0 to 1, where 0 indicates that the two clustering labels are independent to each other, and 1 means that the two clustering labels are the same up to a permutation. We obtained clustering results following the Seurat clustering pipeline with its default setting, and assessed ARI by comparing identified clusters and cell type annotations.

#### NMI

NMI computes normalized mutual information between two clustering results, ranging from 0 to 1. An NMI value close to 0 means that there is nearly no mutual information, while a value close to 1 indicates high correlation between the two clustering results. Similar to ARI, we calculated NMI with clusters identified by the Seurat pipeline and cell type annotations.

#### Cell type ASW

Cell type ASW evaluates ASW with respect to cell type labels, where a higher score means that cells are closer to cells of the same cell type. As ASW lies between -1 and 1, we rescaled the score by cell type 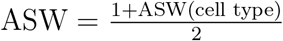.

#### Graph cLISI

The original cLISI measures the effective number of cell types in a neighborhood, where 1 means that the cell population is well preserved, and larger values indicate the mixing of different cell populations. Graph cLISI extends cLISI by enabling the calculation on graphs. The values were rescaled to [0, 1], where higher values indicated good performance of preserving biological variation.

#### Isolated label F1

Isolated label F1 is developed to measure the ability of integration methods to preserve dataset-specific cell types. We adopted the Seurat pipeline to cluster cells in the integrated dataset, and evaluated the cluster assignment of dataset-specific cell types based on the F1 score [79]. Isolated label F1 ranges between 0 and 1, where 1 shows that all cells of dataset-specific cell types are captured in seperate clusters.

#### Isolated label silhouette

Isolated label silhouette, which is similar to Isolated label F1, also measures the conservation of dataset-specific cell types. Instead of using the F1 score, it evaluates ASW of dataset-specific cell types. In our experiments, we rescaled the score to [0,1].

#### Cell cycle conservation

Cell cycle conservation measures how well the cell cycle effect is preserved by integration approaches. It compares cell cycle scores before integration (*CC*_before_) and after integration (*CC*_before_) by calculating 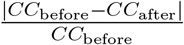, where score 0 indicates perfect conservation of cell cycle effects. We used the gene list from the study [80] as reference, and calculated the cell cycle score based on the Scanpy pipeline. We rescaled the score to [0,1] such that a higher score indicates a better result.

### Benchmarking of the running time and the memory usage

Standard data preprocessing such as normalization, feature selection and dimension reduction could be performed incrementally using mini-batches to control memory usage. In Portal’s preprocessing, we adopted the incremental strategy and used a chunk size of 20,000. For example, the preprocessing of Portal took 63.4 minutes, requiring 22.0 GB peak running memory on the two mouse brain atlases datasets with 1,100,167 cells. The preprocessing time could be reduced to 37.7 minutes when the chunk size was increased to 200,000, with 36.4 GB peak running memory. Some other methods may not be able to adopt a mini-batch implementation. For the two mouse brain atlases datasets, Harmony took 17.6 minutes to finish preprocessing, but required 127.1 GB memory usage. Online iNMF performed preprocessing with mini-batches. Its default preprocessing procedure on the two mouse brain atlases datasets took 15.9 hours, with 0.6 GB memory usage. For a fair comparison, the time and memory usages of data preprocessing procedures were not included in our benchmarking.

### Integration of multiple datasets

For multiple datasets, Portal integrates them in an incremental manner, by transferring all other datasets into the domain constructed by the first dataset. Here we used the integration of two scRNA-seq datasets (profiled by Drop-seq and 10X) [8, 9] and one snRNA-seq dataset (profiled by SPLiT-seq) [49] to illustrate this procedure. In this example, Portal ran in two steps:

#### Step 1

Portal trained domain translation networks between the 10X dataset (160,678 cells) and the Drop-seq dataset (319,359 cells), which took 45.48s. Then Portal used the trained networks to map 10X cells to the Drop-seq dataset domain, which took 0.08s.

#### Step 2

Portal trained domain translation networks between the SPLiT-seq dataset (74,159 cells) and the integrated 10X and Drop-seq dataset, which took 41.36s. Then Portal mapped SPLiT-seq cells to the integrated 10X and Drop-seq dataset domain, which took 0.06s.

In total, Portal took 86.98s to integrate all three datasets.

The integration of multiple datasets is implemented in one function in Portal package. The code for reproducing the experiment is available as a Jupyter Notebook at https://github.com/YangLabHKUST/Portal, serving as an example for the integration of multiple datasets.

### Visualization

We used the UMAP algorithm [39] for visualization of cell representations in a two-dimensional space. In all analyses, the UMAP algorithm was run with 30-nearest neighbors, minimum distance 0.3, and correlation metric.

## Acknowledgements

The authors would like to thank Camille Sophie Ezran (Stanford University), Dr. Angela Oliveira Pisco (CZ Biohub), and Dr. Hosu Sin (Stanford University) for valuable discussions. This work is supported in part by Hong Kong Research Grant Council [16101118, 24301419, 14301120, 16307818, 16301419, 16308120], the Hong Kong University of Science and Technology’s startup grant [R9405,R9364], the Hong Kong University of Science and Technology Big Data for Bio Intelligence Laboratory (BDBI), the Lo Ka Chung Foundation through the Hong Kong Epigenomics Project, the Chau Hoi Shuen Foundation, the Chinese University of Hong Kong direct grants [4053360, 4053423, 4053476], the Chinese University of Hong Kong startup grant [4930181], the Chinese University of Hong Kong’s Project Impact Enhancement Fund (PIEF) and Science Faculty’s Collaborative Research Impact Matching Scheme (CRIMS), the East China Normal University startup grant, the Shanghai Sailing Program. The computational task for this work was partially performed using the X-GPU cluster supported by the RGC Collaborative Research Fund: C6021-19EF.

## Author contributions

J.Z. and G.W. conceived and developed the method. A.R.W. and C.Y. supervised the project. J.Z., G.W., Z.L., A.R.W. and C.Y. designed the experiments, performed the analyses and wrote the paper. J.M., Y.W. and T.M.C. provided critical feedback during the study and helped revise the manuscript.

## Data availability

All data used in this work are publicly available through online sources.

- Mouse brain cells from Saunders et al. [8] (http://dropviz.org).
- Mouse brain cells from Zeisel et al. [9] (http://mousebrain.org/downloads.html).
- Mouse brain cells from Rosenberg et al. [49] (GSE110823).
- Mouse cell atlas from the Tabula Muris Consortium [7] (https://figshare.com/projects/Tabula_Muris_Transcriptomic_characterization_of_20_organs_and_tissues_from_Mus_musculus_at_single_cell_resolution/27733).
- Mouse lemur cell atlas from Tabula Microcebus Consortium (https://figshare.com/projects/Tabula_Microcebus/112227).
- Human peripheral blood mononuclear cells from Mimitou et al. [54] (GSE156478).
- Human peripheral blood mononuclear cells from Ding et al. [81] (GSE132044).
- Human peripheral blood mononuclear cells from 10X Genomics. [81] (https://support.10xgenomics.com/single-cell-gene-expression/datasets/1.1.0/pbmc3k).
- Mouse spermatogenesis cells from Ernst et al. [59] (https://www.ebi.ac.uk/arrayexpress/experiments/E-MTAB-6946/).
- Human spermatogenesis cells from Shami et al. [16] (GSE142585).
- Macaque spermatogenesis cells from Shami et al. [16] (GSE142585).
- Hematopoietic stem cells from Paul et al. [82] (GSE72857).
- Hematopoietic stem cells from Nestorowa et al. [83] (GSE81682).
- Reprogramming of induced pluripotent stem cells from Schiebinger et al. [84] (GSE122662).
- Human brain cells from Fullard et al. [51] (GSE164485).
- Human brain cells from Tran et al. [52] (https://github.com/LieberInstitute/10xPilot_snRNAseq-human#work-with-the-data).

## Code availability

Portal software is available at https://github.com/YangabHKUST/Portal.

