## Supplementary Information for "Adversarial domain translation networks for fast and accurate integration of large-scale atlas-level single-cell datasets"

#### Supplementary Information: Note

##### Investigation of the role of each component in Portal

In this section, we investigate the role of each component in Portal. The optimization problem solved by Portal is

$$\min_{\{E_1,G_1,E_2,G_2\}} \max_{\{D_1,D_2\}} \quad \mathcal{L}_{\text{adv}} + \lambda_{\text{cos}}\mathcal{R}_{\text{cos}} + \lambda_{\text{LA}}\mathcal{R}_{\text{LA}} + \lambda_{\text{AE}}\mathcal{R}_{\text{AE}}, \tag{S1}$$

where $E_1(\cdot)$ and $E_2(\cdot)$ are encoder networks; $G_1(\cdot)$ and $G_2(\cdot)$ are generator networks; $D_1(\cdot)$ and $D_2(\cdot)$ are discriminator networks; $\mathcal{L}_{\text{adv}}$ stands for the adversarial learning objective, whose value is $\mathcal{L}_{\mathcal{X}} + \mathcal{L}_{\mathcal{Y}}$ when maximizing with respect to $D_1(\cdot), D_2(\cdot)$, and it is replaced with

*These authors contributed to this work equally.

$\mathcal{L}_{\mathcal{X}}^{\log D} + \mathcal{L}_{\mathcal{Y}}^{\log D}$ when minimizing with respect to $E_1(\cdot), G_1(\cdot), E_2(\cdot), G_2(\cdot)$ according to the "logD-trick"; $\mathcal{R}_{\text{cos}} = \mathcal{R}_{\text{cos}}(E_1, G_1, E_2, G_2)$ is a regularizer for the cosine similarity correspondence cross domains; $\mathcal{R}_{\text{LA}} = \mathcal{R}_{\text{LA}}(E_1, G_1, E_2, G_2)$ is a regularizer for the alignment consistency in the latent space; $\mathcal{R}_{\text{AE}} = \mathcal{R}_{\text{AE}}(E_1, G_1, E_2, G_2)$ is a regularizer for the autoencoder consistency; $\lambda_{\text{cos}}$, $\lambda_{\text{LA}}$, $\lambda_{\text{AE}}$ are coefficients for the three regularizers respectively. To demonstrate the roles of the objective function $\mathcal{L}_{\text{adv}}$ and three regularizers ($\mathcal{R}_{\text{cos}}$, $\mathcal{R}_{\text{LA}}$, and $\mathcal{R}_{\text{AE}}$), we rewrite the optimization problem (S1) as

$$\min_{\{E_1,G_1,E_2,G_2\}} \max_{\{D_1,D_2\}} \quad \lambda_{\text{adv}}\mathcal{L}_{\text{adv}} + \lambda_{\text{cos}}\mathcal{R}_{\text{cos}} + \lambda_{\text{LA}}\mathcal{R}_{\text{LA}} + \lambda_{\text{AE}}\mathcal{R}_{\text{AE}}, \tag{S2}$$

with $\lambda_{\text{adv}}$ set to 1.0 in Portal's algorithm. Based on (S2), we are able to study on the impact of each component of Portal by manually setting the corresponding coefficient to zero, and then compare its performance with that of the standard algorithm empirically. Recall that the discriminators are designed to deal with domain-unique cell types by discriminator score thresholding. In this section, we also experimentally verify the effectiveness of such design. Here we took mouse mammary gland scRNA-seq atlas from the Tabula Muris consortium as an example. In the mouse mammary gland data, 4,481 cells were profiled by 10X Genomics (10X), and 2,405 cells were profiled by SMART-seq2 (SS2). With these two datasets, we investigate the role of each component in Portal.

**Role of objective function $\mathcal{L}_{\text{adv}}$.** The objective function $\mathcal{L}_{\text{adv}}$ plays an essential role in learning effective domain translation across different datasets. To demonstrate the importance of adversarial training using $\mathcal{L}_{\text{adv}}$, we removed it from Portal by setting $\lambda_{\text{adv}}$ to zero, then applied this version of Portal (Portal ($\lambda_{\text{adv}} = 0$)) to integrate mouse mammary gland datasets. Comparison between integration results obtained by Portal (Fig. S1**a**) and Portal ($\lambda_{\text{adv}} = 0$)) (Fig. S1**b**) confirmed that cells from different datasets could not be well mixed without the objective function.

**Role of regularizer $\mathcal{R}_{\text{cos}}$.** Regularizer $\mathcal{R}_{\text{cos}}$ helps to establish reliable alignment between different domains. It guides domain translation networks to find correspondence of the same cell type across domains. To confirm this, we fixed $\lambda_{\text{cos}}$ in (S2) as zero to remove $\mathcal{R}_{\text{cos}}$ from Portal, and we denoted this version of Portal as Portal ($\lambda_{\text{cos}} = 0$). After applying Portal ($\lambda_{\text{cos}} = 0$), cells from the two datasets were well mixed, however, the obtained alignment between these datasets was problematic. For example, basal cells from SS2 dataset were incorrectly aligned with B cells and T cells from 10X dataset (Fig. S1**c**). In contrast, the standard version of

Portal built the alignment correctly (Fig. S1**a**). The difference between results obtained by Portal and Portal ($\lambda_{\text{cos}} = 0$) verified the usefulness of $\mathcal{R}_{\text{cos}}$ to establish robust correspondence between datasets.

**Role of regularizer $\mathcal{R}_{\text{LA}}$.** Regularizer $\mathcal{R}_{\text{LA}}$ is introduced to impose the consistency constraint for latent representations of cells. It is helpful to remove domain-specific effects in the latent space. To demonstrate the effectiveness of $\mathcal{R}_{\text{LA}}$, we set $\lambda_{\text{LA}} = 0$. For Portal ($\lambda_{\text{LA}} = 0$), the learned representation in the latent space showed a poor alignment of two datasets (Fig. S1**d**). This result indicated that the learned representation in the latent space would not be a valid integration result without adopting regularizer $\mathcal{R}_{\text{LA}}$.

**Role of regularizer $\mathcal{R}_{\text{AE}}$.** $\{E_1(\cdot), G_1(\cdot)\}$ and $\{E_2(\cdot), G_2(\cdot)\}$ form two autoencoder structures in Portal's framework, $\mathcal{R}_{\text{AE}}$ is hence introduced for regularizing autoencoder consistency. Here we set $\lambda_{\text{AE}} = 0$ to evaluate the role of $\mathcal{R}_{\text{AE}}$ with Portal ($\lambda_{\text{AE}} = 0$). Comparison between results obtained by Portal (Fig. S1**a**) and Portal ($\lambda_{\text{AE}} = 0$) (Fig. S1**e**) indicated that $\mathcal{R}_{\text{AE}}$ was useful to improve the accuracy of Portal's results by imposing the consistency between encoders and generators.

**Role of discriminator score thresholding.** The discriminator score thresholding in Portal is a tailored design for single-cell integration tasks. With such design, Portal does not force the alignment of domain-unique cell types, preventing overcorrection of domain-specific effects. To illustrate the role of the discriminators, we used the same mouse mammary gland data. We manually removed all basal cells from the 10X dataset and thereby basal cell type became a domain-unique cell type in the SS2 dataset. We applied standard Portal and Portal without discriminator score thresholding (denoted as "Portal w/o $D$ score thresholding") for integration. The results in Fig. S2 indicated that, without discriminator score thresholding, Portal could not retain the identity of basal cells in the SS2 dataset, and incorrectly aligned them with T cells and B cells in the 10X dataset.

#### Parameter sensitivity analysis of Portal

Hyperparameters used in Portal are $\lambda_{\text{cos}}$, $\lambda_{\text{LA}}$, $\lambda_{\text{AE}}$, $t$ and $m$:

- $\lambda_{\text{cos}}$ is coefficient for cosine similarity regularizer $\mathcal{R}_{\text{cos}}$ that guides domain translation networks in Portal to establish reliable alignment between cells from different datasets;
- $\lambda_{\text{LA}}$ is coefficient for latent alignment regularizer $\mathcal{R}_{\text{LA}}$ which is helpful to remove domain-

specific effects in the latent space by imposing the consistency constraint for latent representations of cells;

- $\lambda_{\text{AE}}$ is coefficient for autoencoder consistency regularizer $\mathcal{R}_{\text{AE}}$ that regularizes two autoencoder structures {encoder $E_1(\cdot)$, generator $G_1(\cdot)$} and {encoder $E_2(\cdot)$, generator $G_2(\cdot)$};

- $t$ is the absolute value of boundaries for the logit of discriminator scores ($-t < d_i(\mathbf{x}) < t$, $i = 1, 2$). It is included as a tailored design for preventing overcorrection in integration tasks;

- $m$ is the batch size utilized by Portal for mini-batch training.

We have observed that the parameter setting $\lambda_{\text{cos}} = 20.0$, $\lambda_{\text{LA}} = 10.0$, $\lambda_{\text{AE}} = 10.0$, $t = 5.0$ and $m = 500$ (default setting in Portal package) yields reasonable results in integrative analyses of a collection of diverse single-cell datasets. Throughout all analyses, we achieved slightly better alignment results by tuning $\lambda_{\text{cos}}$ according to the mixing metric in Seurat [1], while we fixed all other parameters at their default values.

To examine whether Portal is sensitive to its parameters, we applied Portal to two human peripheral blood mononuclear cells (PBMCs) datasets [2]. We varied one hyperparameter at a time, and fixed other hyperparameters at their default setting: (**a**) $\lambda_{\text{cos}}$ in {10, 15, 20, 25, 30, 35, 40, 45, 50}; (**b**) $\lambda_{\text{LA}}$ in {2, 4, 6, 8, 10, 12, 14, 16, 18}; (**c**) $\lambda_{\text{AE}}$ in {2, 4, 6, 8, 10, 12, 14, 16, 18}; (**d**) $t$ in {3, 4, 5, 6, 7, 8}; and (**e**) $m$ in {200, 500, $1,000$}. We visualized the integration results from Portal with UMAP plots in Fig. S3**a** - **e** respectively. As shown in Fig. S3, Portal mixed the two datasets profiled by 10X and inDrops well and aligned cell types correctly in different hyperparameter settings, indicating that Portal is insensitive to choice of hyperparameters. Besides the UMAP visualizations, we also quantitatively evaluated the integration results in this sensitivity analysis. We predicted each cell's most likely cell type by label transfer based on integrated results, and compared it to the given label. We used this to measure prediction accuracy, which served as an overall evaluation. To measure the results in terms of correction of batch effects and conservation of biological variation, respectively, we evaluated graph integration local inverse Simpson's Index (graph iLISI) [3, 4] and graph cell type local inverse Simpson's Index (graph cLISI) [3, 4] for all results. The scores were rescaled such that higher graph iLISI and graph cLISI scores indicated better batch effects correction performance and better conservation of biological variation, respectively. For comparison, we also evaluated the three metrics before integrating the two datasets as a baseline. As indicated by Fig. S4**a** - **e**, Portal performed effective integration in all three aspects in different hyperparameter settings. In particular, the prediction accuracy was never below 98%, supporting the robustness of Portal in the parameter sensitivity analysis.

### Supplementary Information: Figures

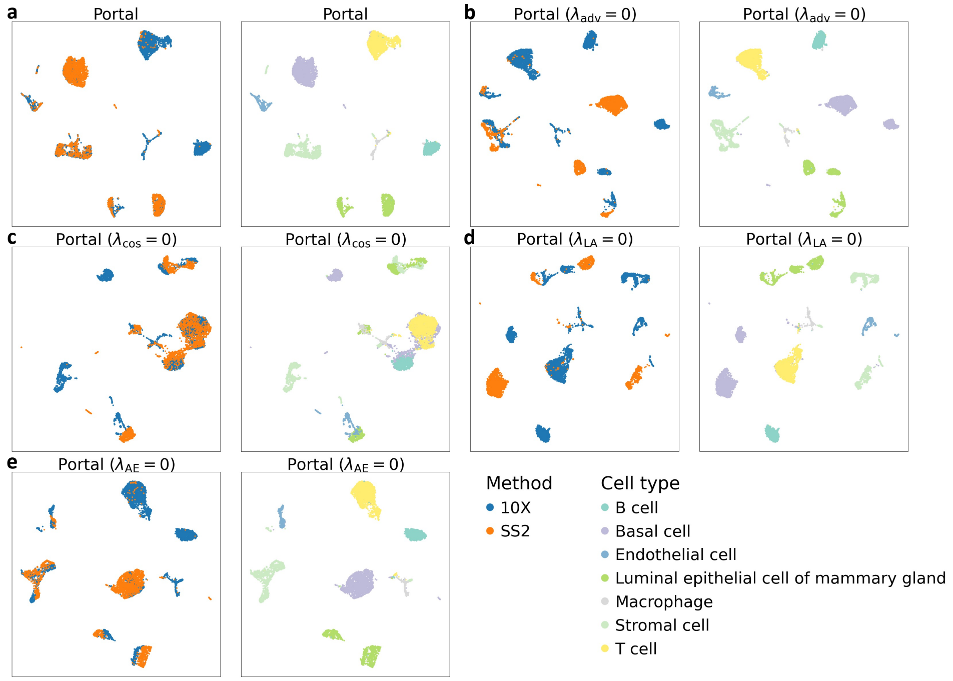


Figure S1: **Investigation of the roles of objective function $\mathcal{L}_{\text{adv}}$ and three regularizers $\mathcal{R}_{\text{cos}}$, $\mathcal{R}_{\text{LA}}$ and $\mathcal{R}_{\text{AE}}$ in Portal.** We used the mouse mammary gland scRNA-seq datasets from Tabluma Muris Consortium in this study. **a**. We applied Portal to integrate the two datasets as a baseline. **b-e**. Then we fixed $\lambda_{\text{adv}}$, $\lambda_{\text{cos}}$, $\lambda_{\text{LA}}$, $\lambda_{\text{AE}}$ in (S2) at zero to evaluate the effectiveness of $\mathcal{L}_{\text{adv}}$, $\mathcal{R}_{\text{cos}}$, $\mathcal{R}_{\text{LA}}$ and $\mathcal{R}_{\text{AE}}$, respectively. Clearly, each component of Poral plays its important role in data integration.

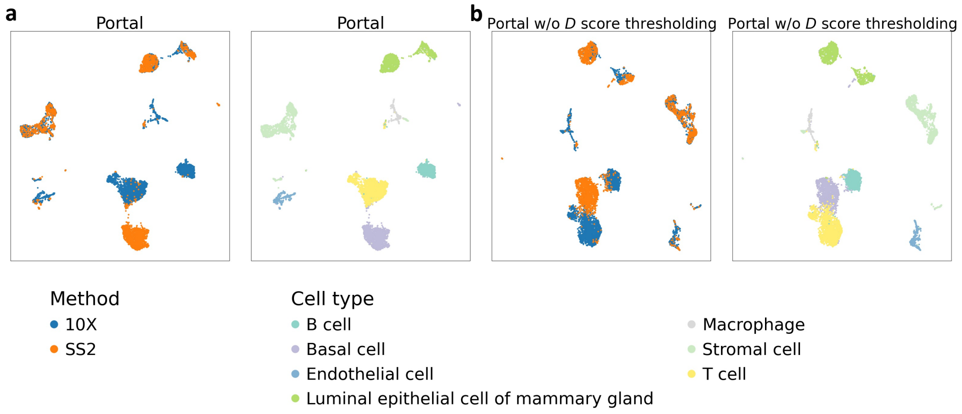


Figure S2: **Investigation of the role of discriminator score thresholding in Portal.** We used the same mouse mammary gland data from Tabluma Muris Consortium, and removed all basal cells from the 10X dataset to make basal cell a domain-unique cell type in the SS2 dataset. **a**. We applied Portal to integrating the two datasets as a baseline. **b**. We removed discriminator score thresholding in Portal to integrate the two datasets.

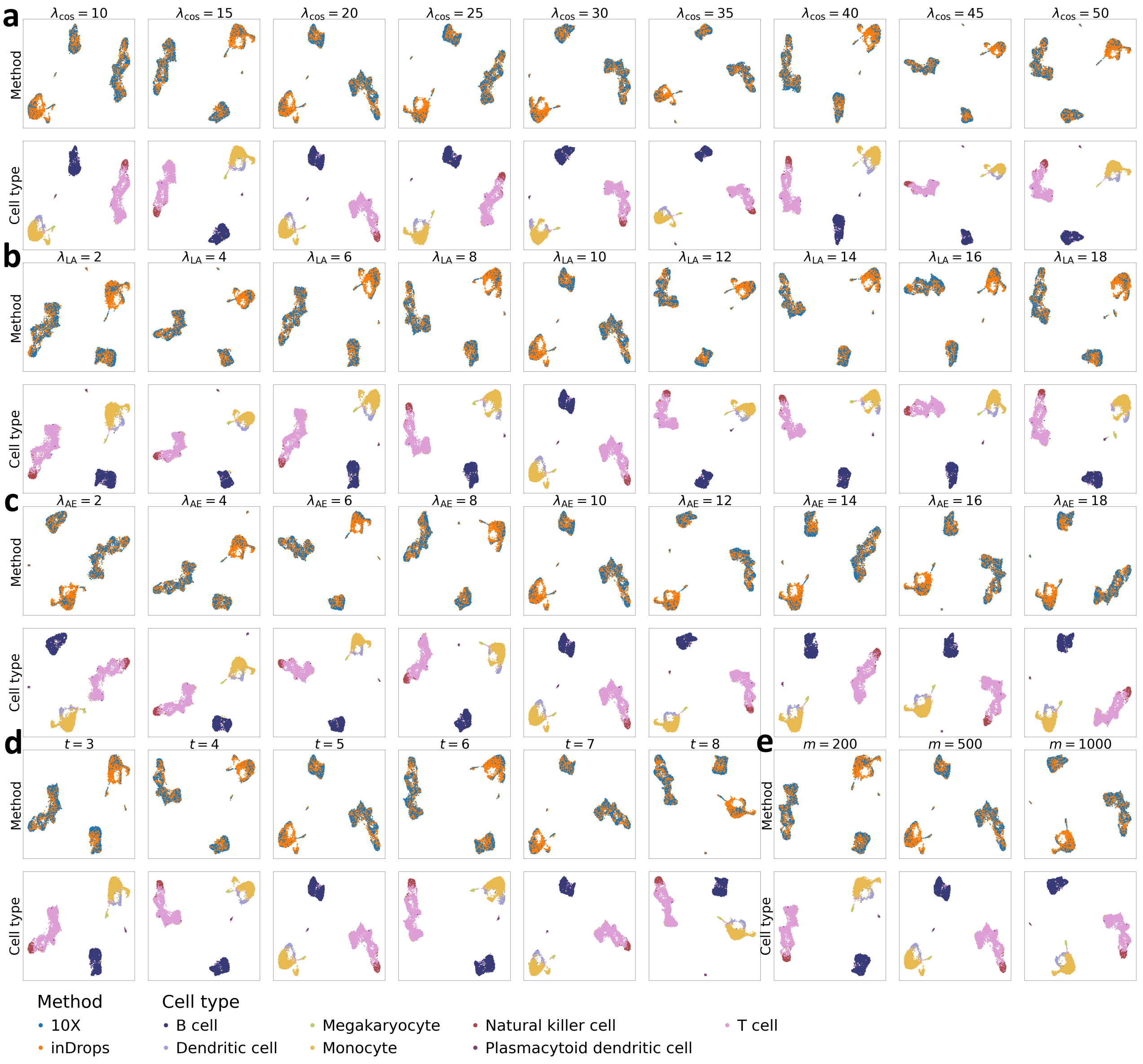


Figure S3: **UMAP visualizations of Portal's integration results in the parameter sensitivity analysis.** We used two human peripheral blood mononuclear cells (PBMCs) datasets from [2] for our parameter sensitivity analysis. UMAP plots were colored by profiling methods and cell types, respectively. We varied hyperparameters in Portal as follows: **a**. $\lambda_{\mathrm{cos}}$ was varied in {10, 15, 20, 25, 30, 35, 40, 45, 50}. **b**. $\lambda_{\mathrm{LA}}$ was varied in {2, 4, 6, 8, 10, 12, 14, 16, 18}. **c**. $\lambda_{\mathrm{AE}}$ was varied in {2, 4, 6, 8, 10, 12, 14, 16, 18}. **d**. $t$ was varied in {3, 4, 5, 6, 7, 8}. **e**. $m$ was varied in {200, 500, 1,000}.

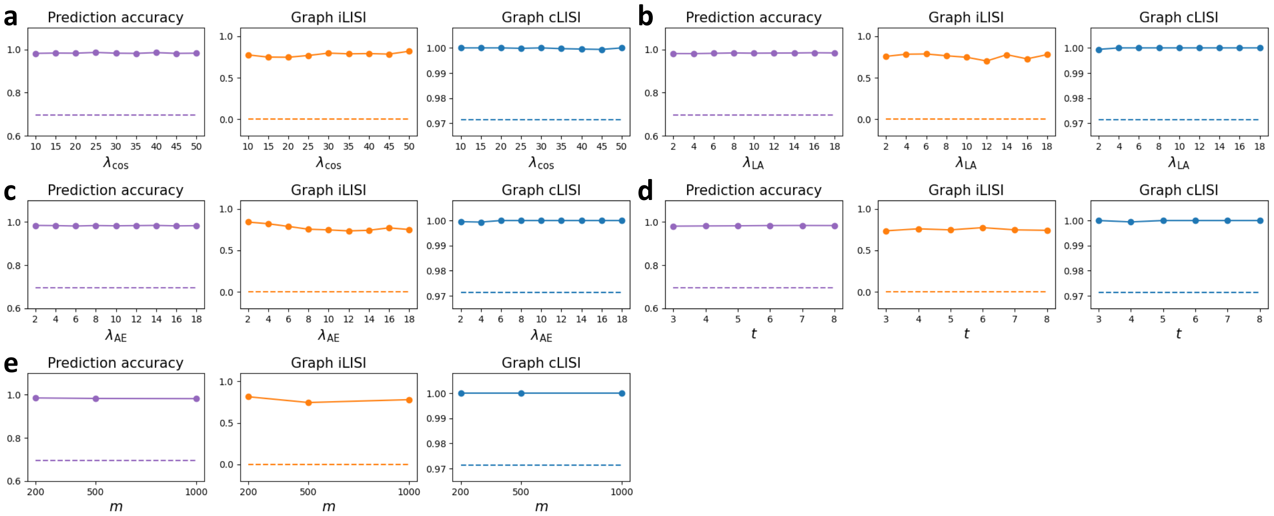


Figure S4: **Quantitative assessment of Portal's integration results in the parameter sensitivity analysis.** We used prediction accuracy as an overall evaluation metric. Graph iLISI and graph cLISI were used to measure the performance in the aspects of batch effects correction and biological variation conservation respectively. We assessed the three metrics with respect to the variation of: **a**. $\lambda_{\text{cos}}$; **b**. $\lambda_{\text{LA}}$; **c**. $\lambda_{\text{AE}}$; **d**. $t$; **e**. $m$ respectively.

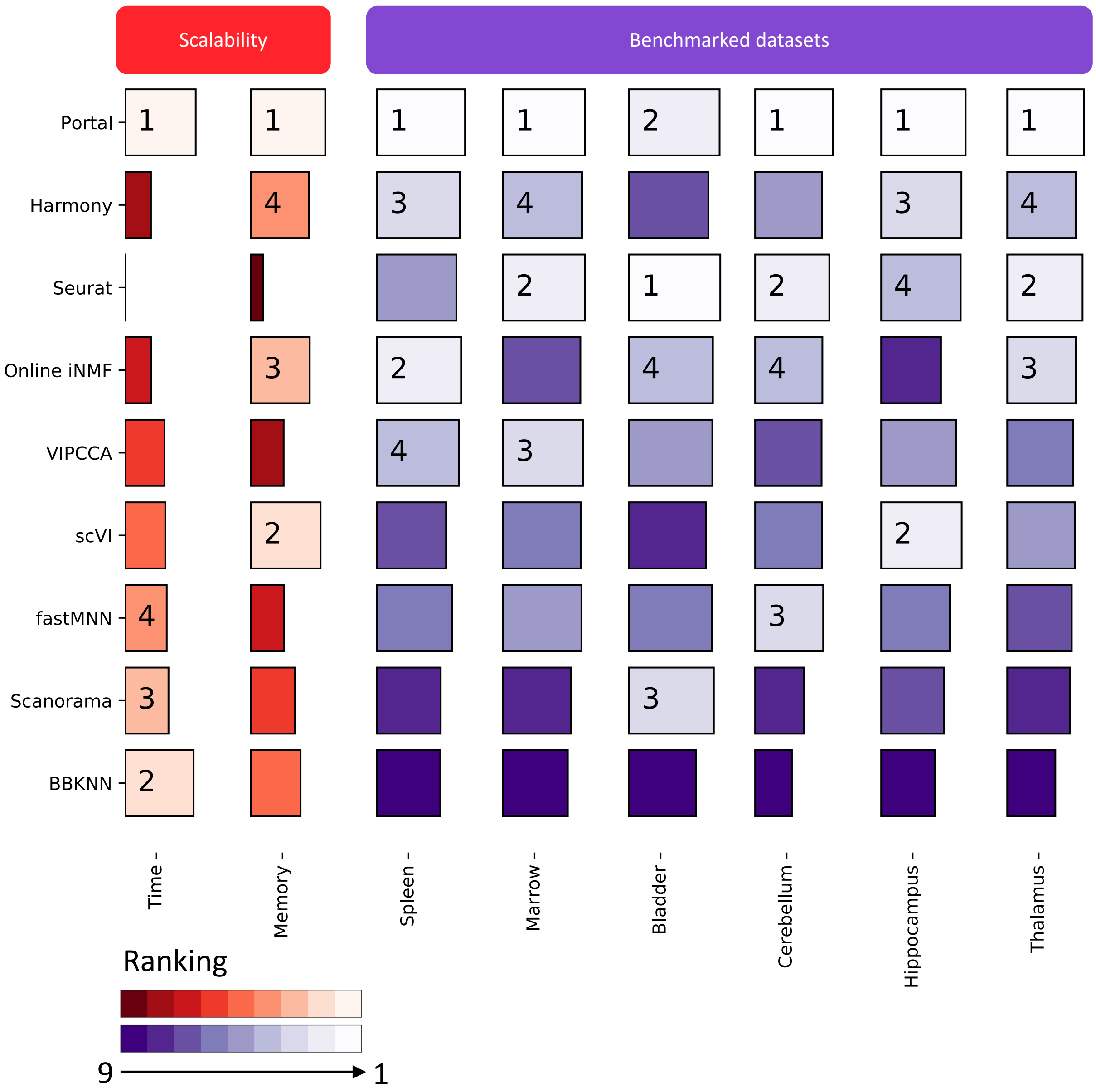


Figure S5: **Benchmarking of Portal, Harmony, Seurat, online iNMF, VIPCCA, scVI, fastMNN, Scanorama and BBKNN using the benchmarking strategy [3].** Among all benchmarked methods, Portal showed best scalability in terms of the running time and the peak running memory usage. It also showed state-of-the-art overall performance scores accounting for both batch correction and conservation of biological variation. Scalability scores were obtained using the running time and the peak running memory required by all methods when integrating 500,000 cells. We started from logarithms of measured time (in seconds) and memory (in MB), then rescaled by a constant baseline and used one minus the values as final scores: $\text{Score}_{time} = 1 - \log(time)/\log(C_{time})$, where $C_{time} = 21,600$ (seconds) (6 hours) was used as a baseline. Seurat took more time than this baseline (8.5 hours), so we set its score to zero. Scores for memory were calculated similarly: $\text{Score}_{memory} = 1 - \log(memory)/\log(C_{memory})$, where $C_{memory} = 1,024^2$ (MB) (1 TB). Calculation of the overall performance scores on each task is shown in Figs. S7, S9, S11, S13, S15 and S17. For each column, top-4 methods were annotated with their corresponding rankings.

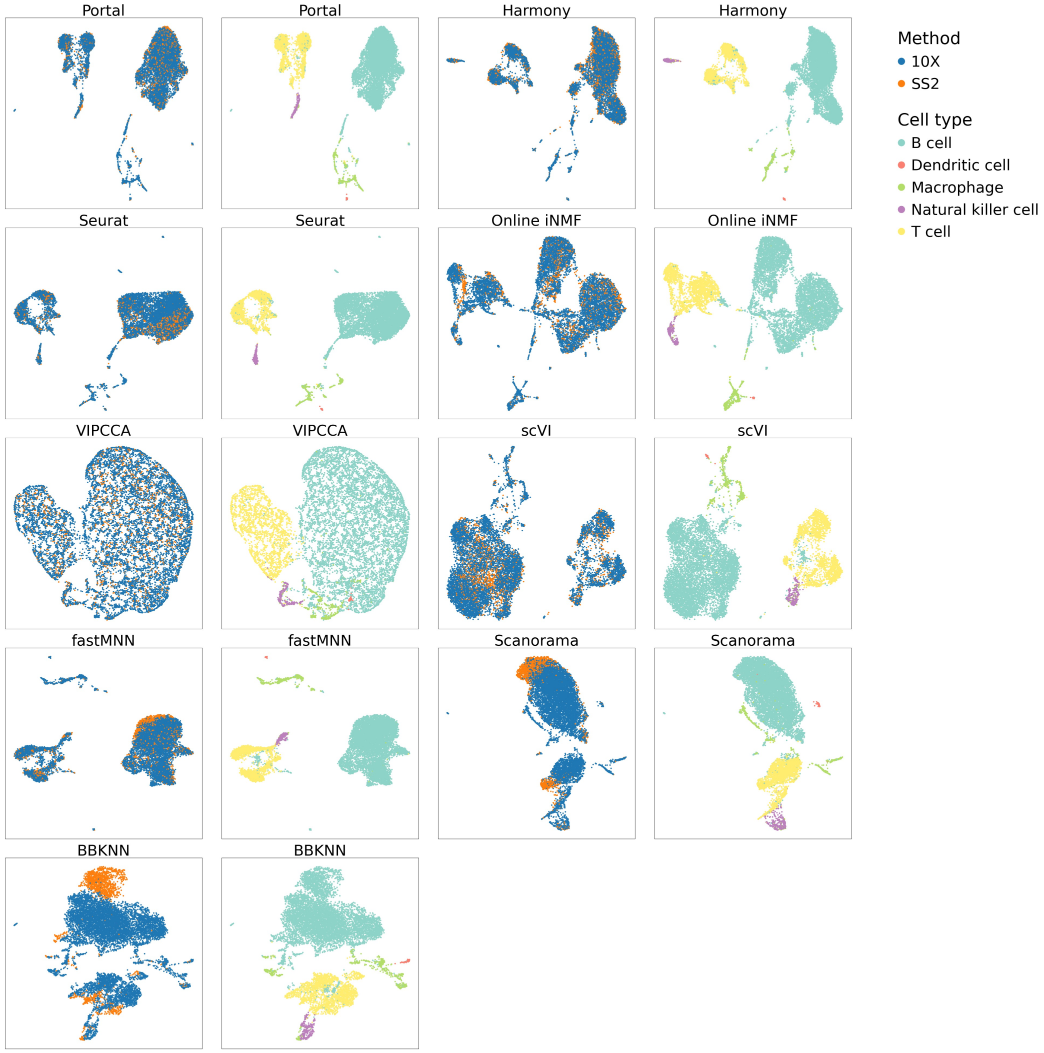


Figure S6: **Comparison of integration methods based on mouse spleen data.** We integrated mouse spleen scRNA-seq datasets profiled by 10X Genomics (10X) and SMART-seq2 (SS2). UMAP plots were colored by profiling methods and cell types, respectively.

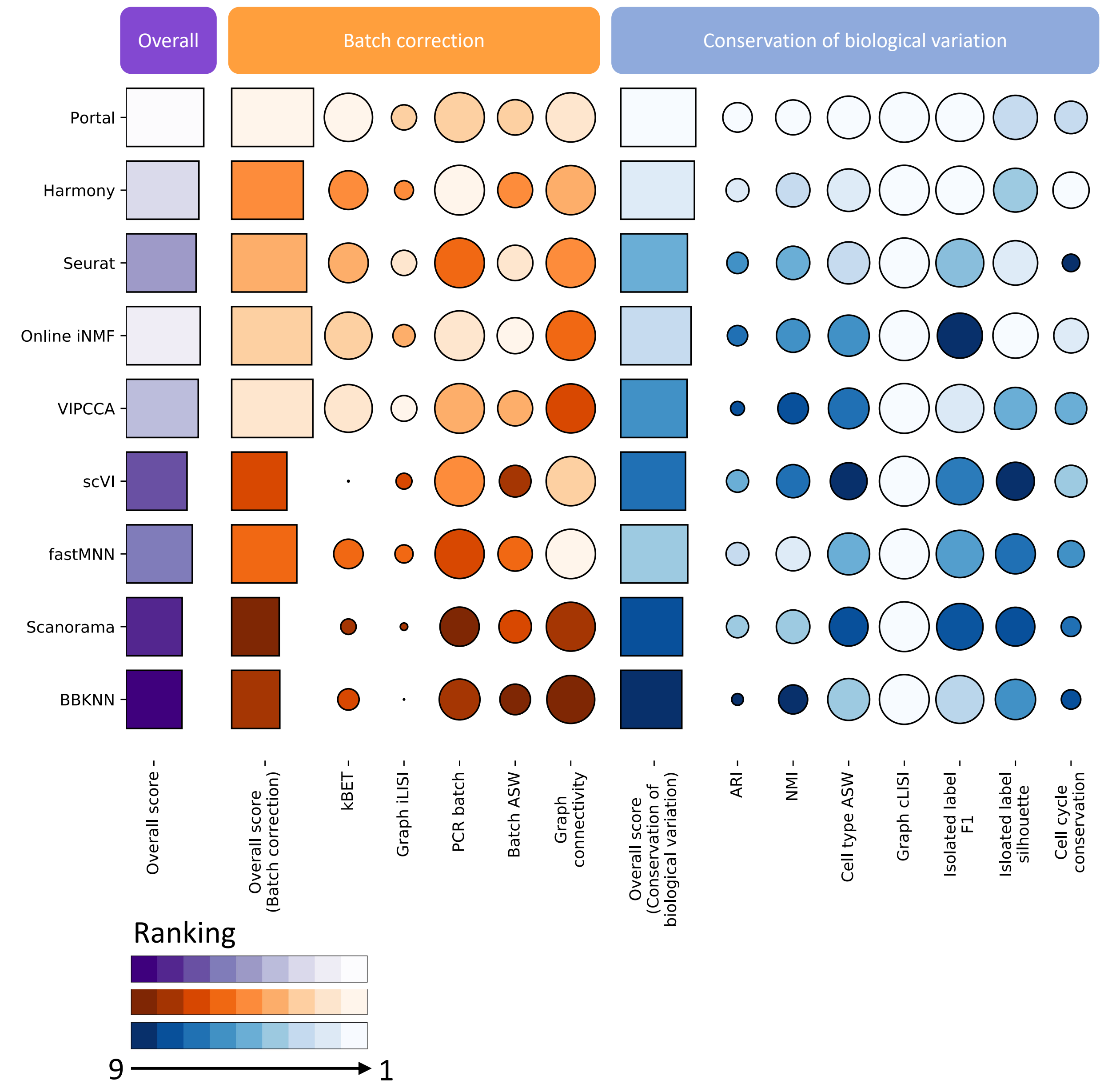


Figure S7: **Benchmarking of integration methods based on mouse spleen data.** Scores for batch correction and scores for conservation of biological variation are computed as the average of metrics which are in their categories. Overall scores are computed by a 40:60 weighted average of scores for batch correction and scores for conservation of biological variation.

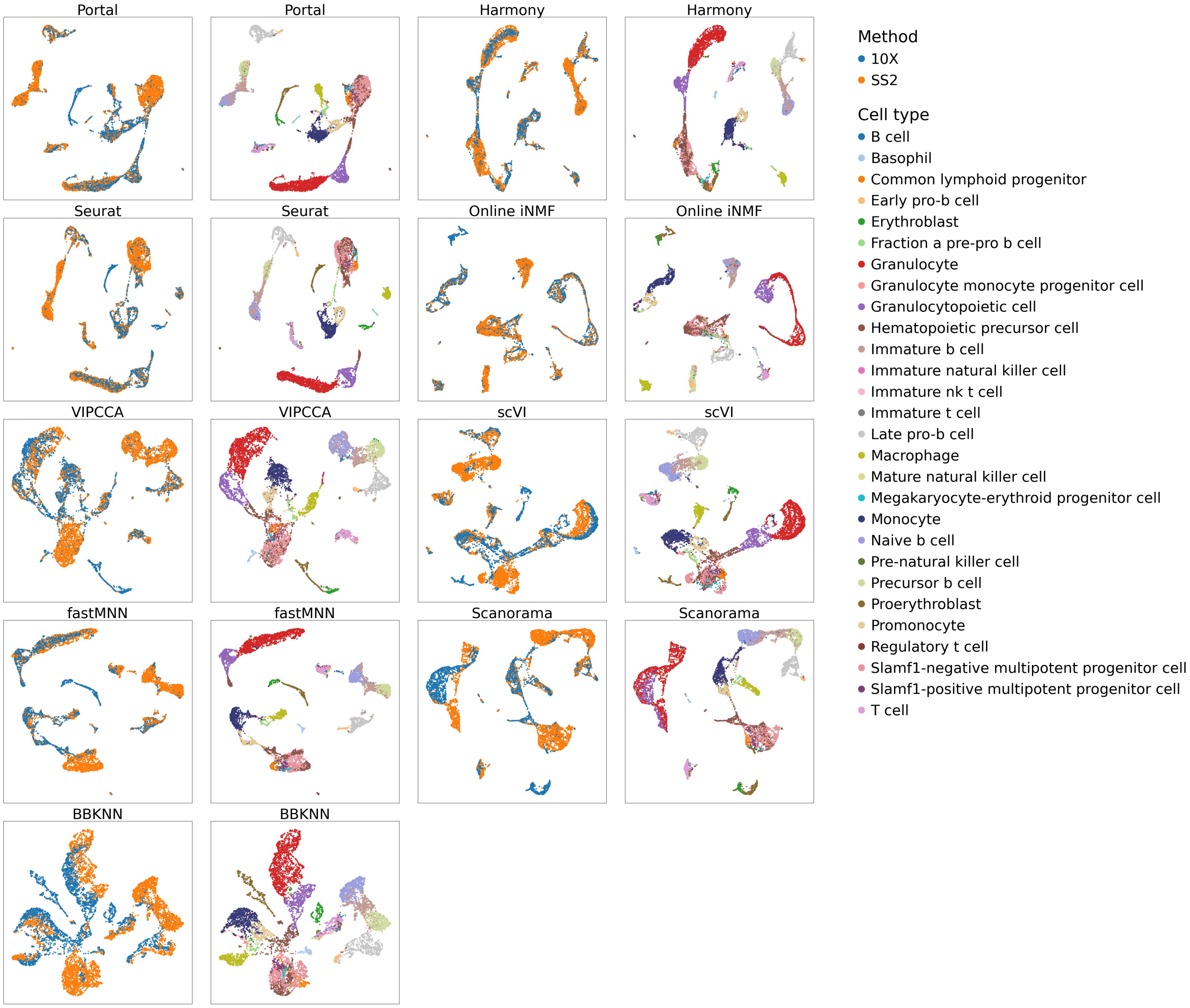


Figure S8: **Comparison of integration methods based on mouse marrow data.** We integrated mouse marrow scRNA-seq datasets profiled by 10X Genomics (10X) and SMART-seq2 (SS2). UMAP plots were colored by profiling methods and cell types, respectively.

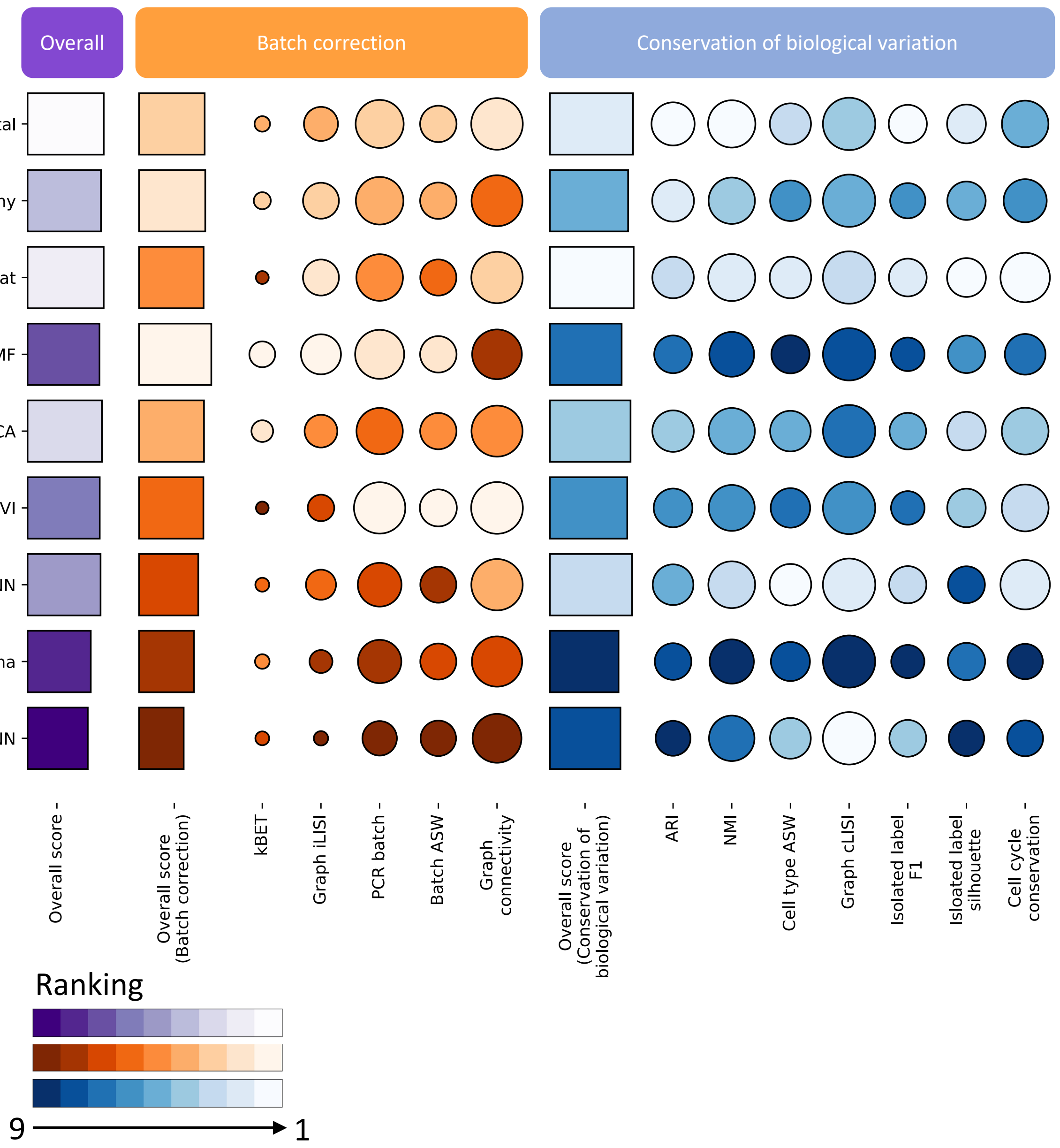


Figure S9: **Benchmarking of integration methods based on mouse marrow data.** Scores for batch correction and scores for conservation of biological variation are computed as the average of metrics which are in their categories. Overall scores are computed by a 40:60 weighted average of scores for batch correction and scores for conservation of biological variation.

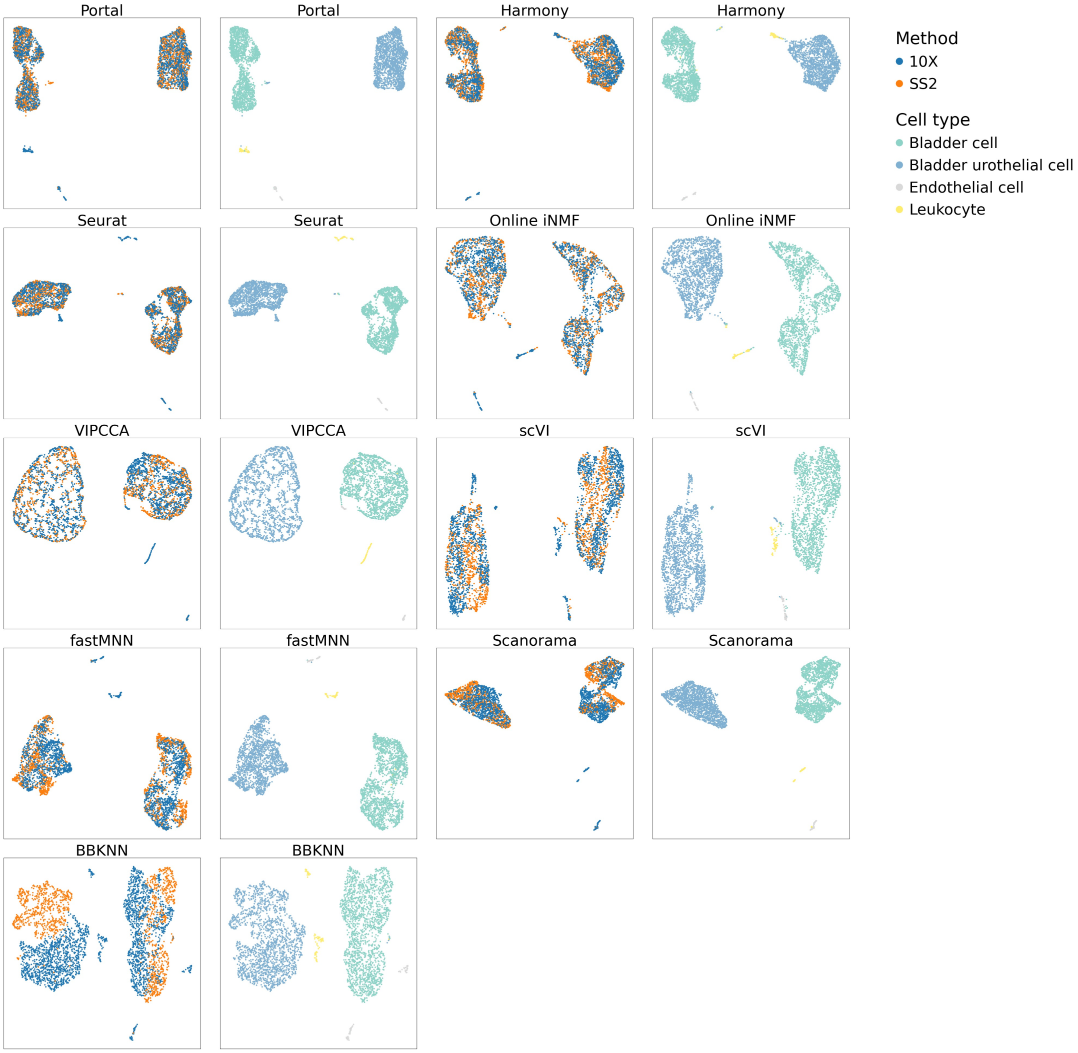


Figure S10: **Comparison of integration methods based on mouse bladder data.** We integrated mouse bladder scRNA-seq datasets profiled by 10X Genomics (10X) and SMART-seq2 (SS2). UMAP plots were colored by profiling methods and cell types, respectively.

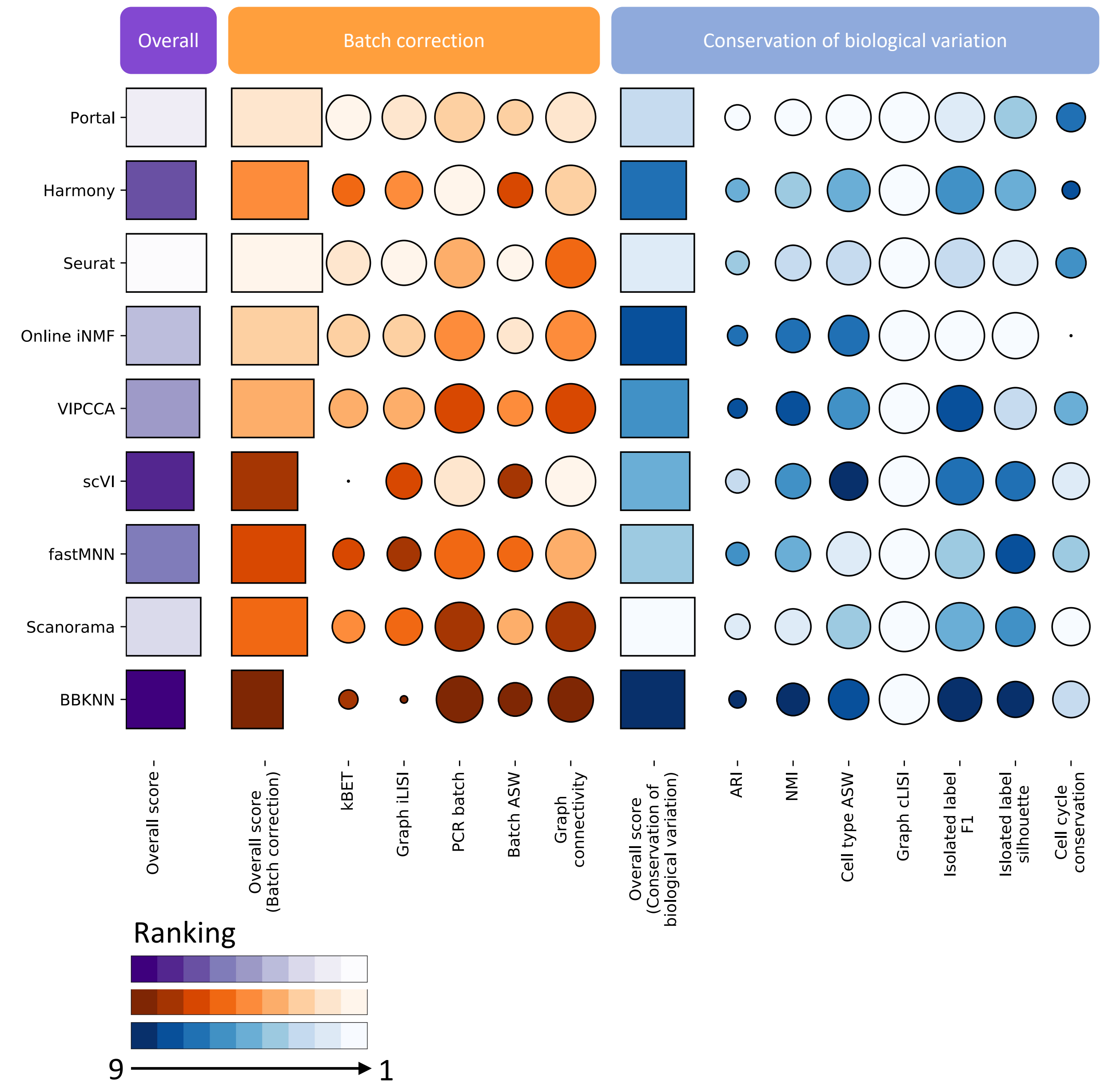


Figure S11: **Benchmarking of integration methods based on mouse bladder data.** Scores for batch correction and scores for conservation of biological variation are computed as the average of metrics which are in their categories. Overall scores are computed by a 40:60 weighted average of scores for batch correction and scores for conservation of biological variation.

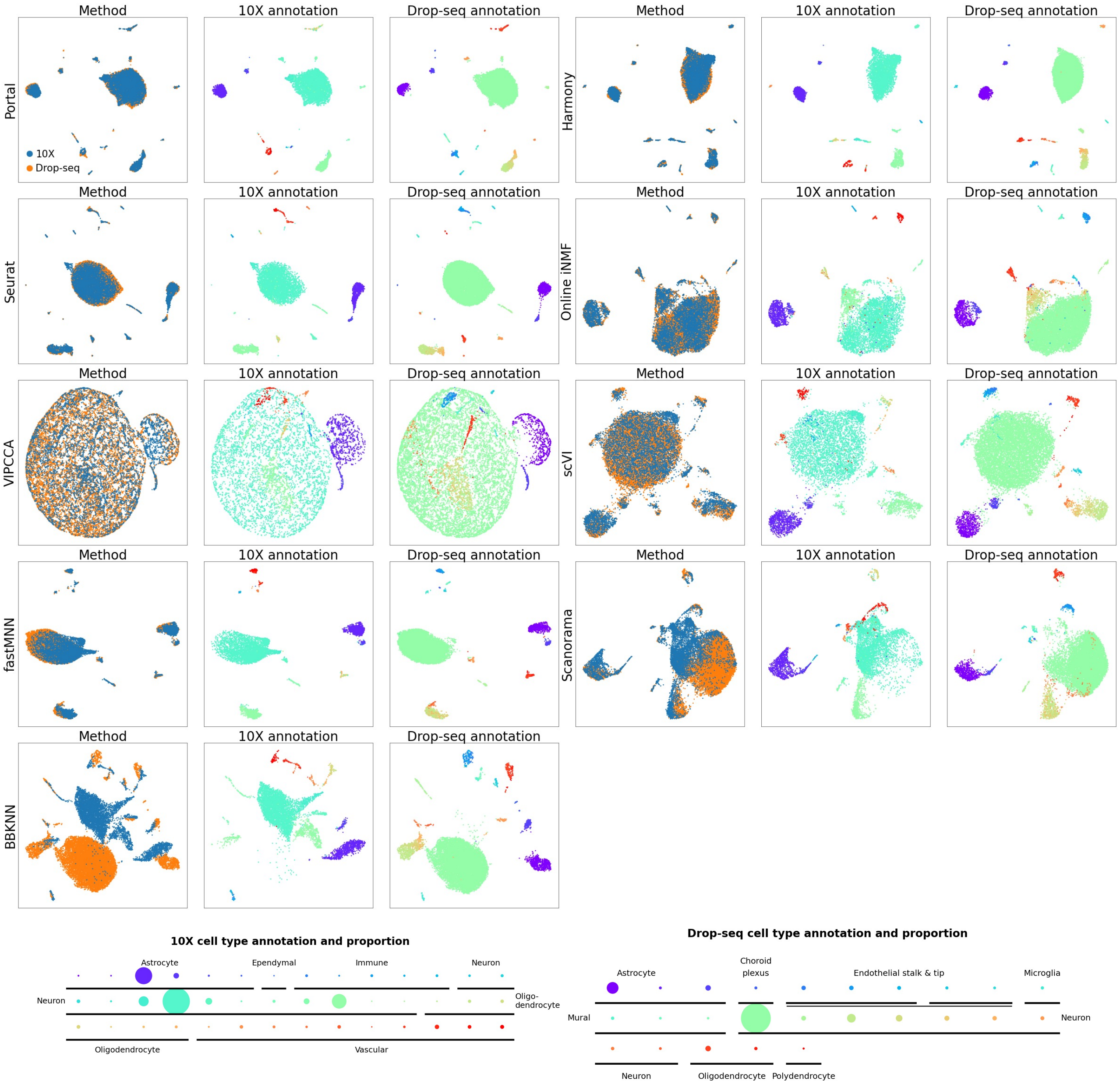


Figure S12: **Comparison of integration methods based on mouse cerebellum data.** We integrated mouse cerebellum scRNA-seq datasets profiled by 10X and Drop-seq. UMAP plots were colored by profiling methods and cell types, respectively. We used dot plots to visualize cell type labels and their proportions. Different colors represent different subpopulations of cells. The comparison among proportions of subpopulations was visualized by the sizes of corresponding dots.

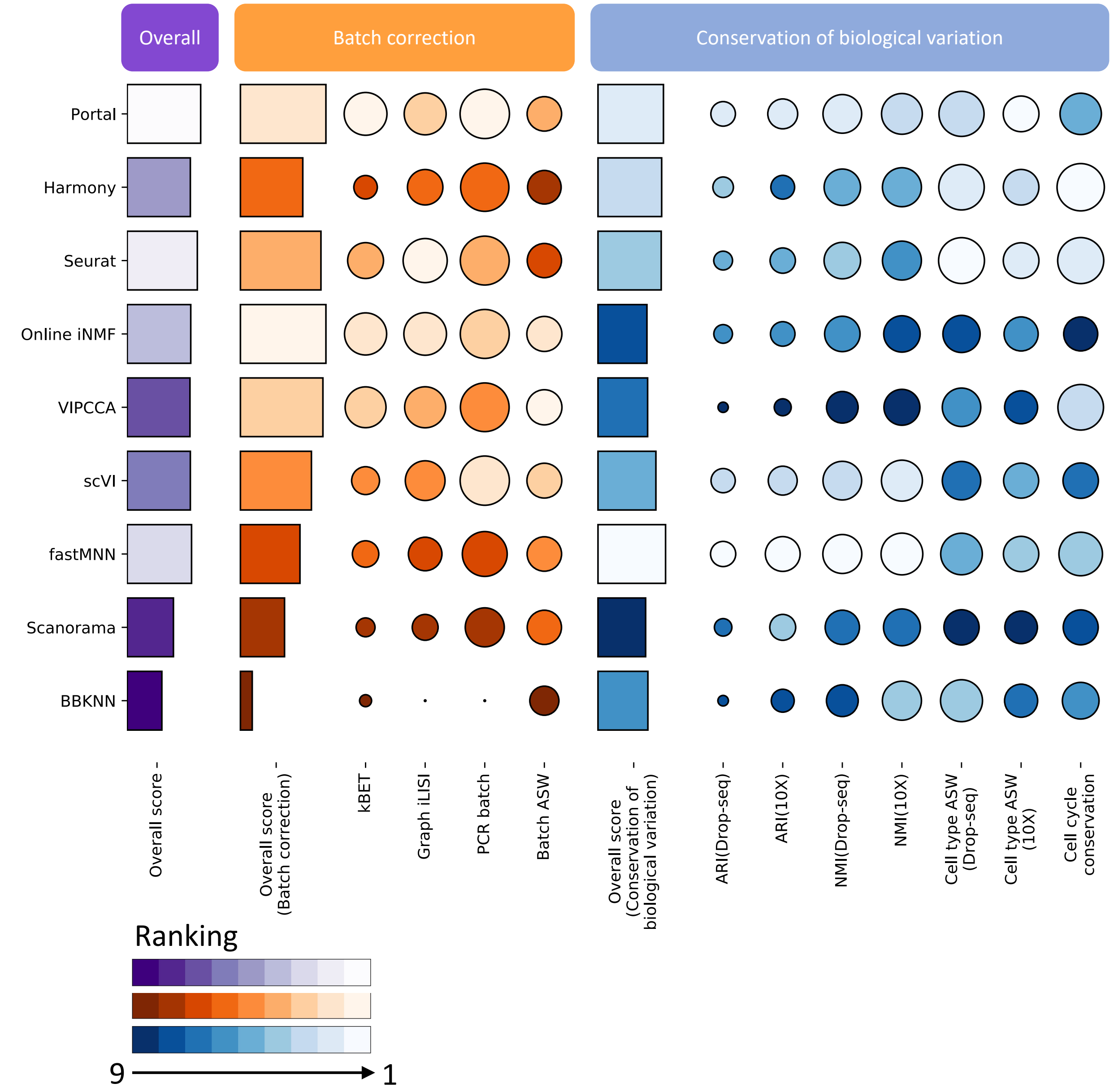


Figure S13: **Benchmarking of integration methods based on mouse cerebellum data.** Scores for batch correction and scores for conservation of biological variation are computed as the average of metrics which are in their categories. As the Drop-seq dataset and the 10X dataset provided detailed annotations in their original publications, we slightly modified the assessment for better evaluation of biological variation conservation. For example, we calculated ARI, NMI and cell type ASW based on both annotations, and used them to replace the metrics that require unified cell type annotations across datasets. Overall scores are computed by a 40:60 weighted average of scores for batch correction and scores for conservation of biological variation.

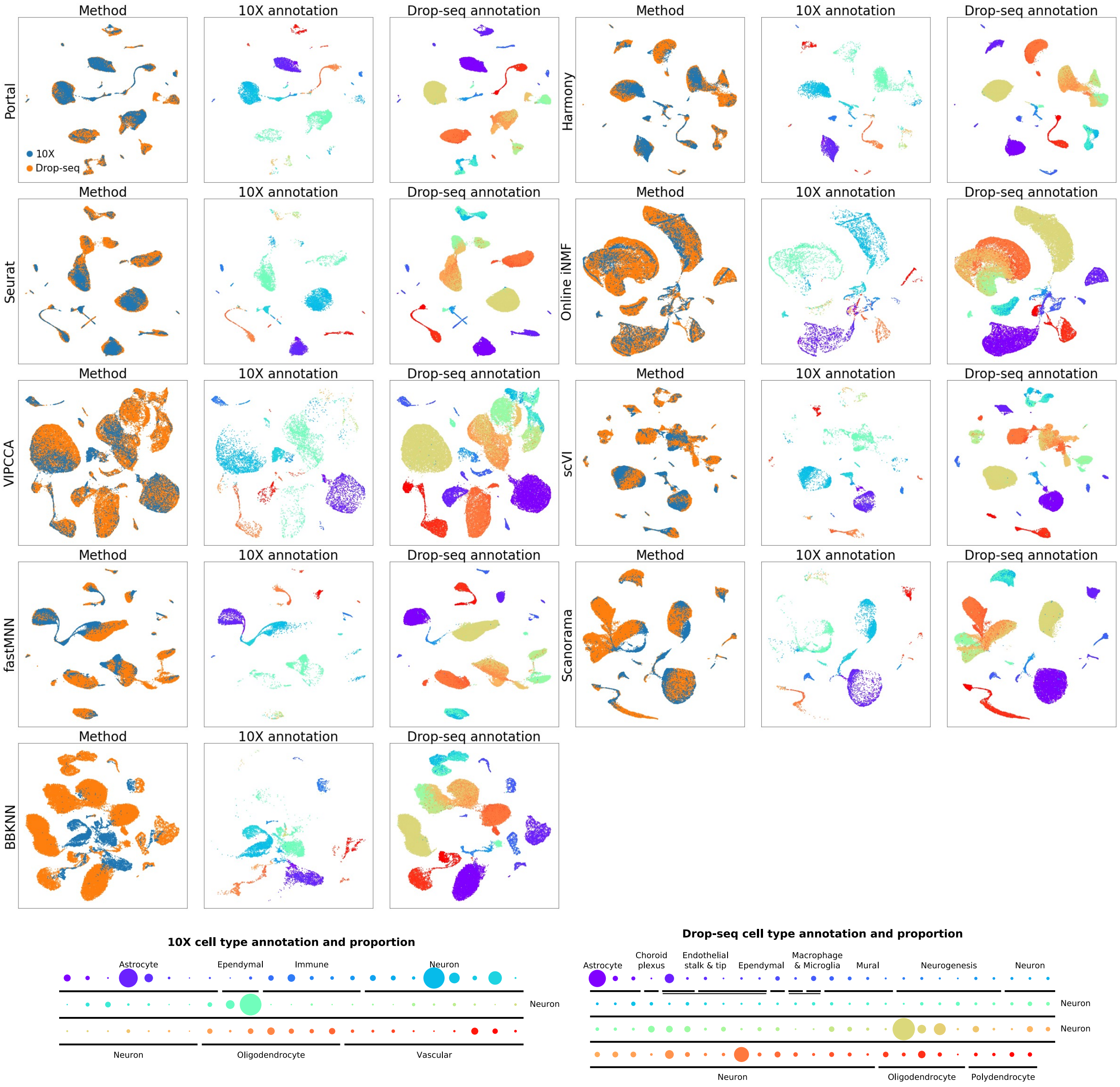


Figure S14: **Comparison of integration methods based on mouse hippocampus data.** We integrated mouse hippocampus scRNA-seq datasets profiled by 10X and Drop-seq. UMAP plots were colored by profiling methods and cell types, respectively. We used dot plots to visualize cell type labels and their proportions. Different colors represent different subpopulations of cells. The comparison among proportions of subpopulations was visualized by the sizes of corresponding dots.

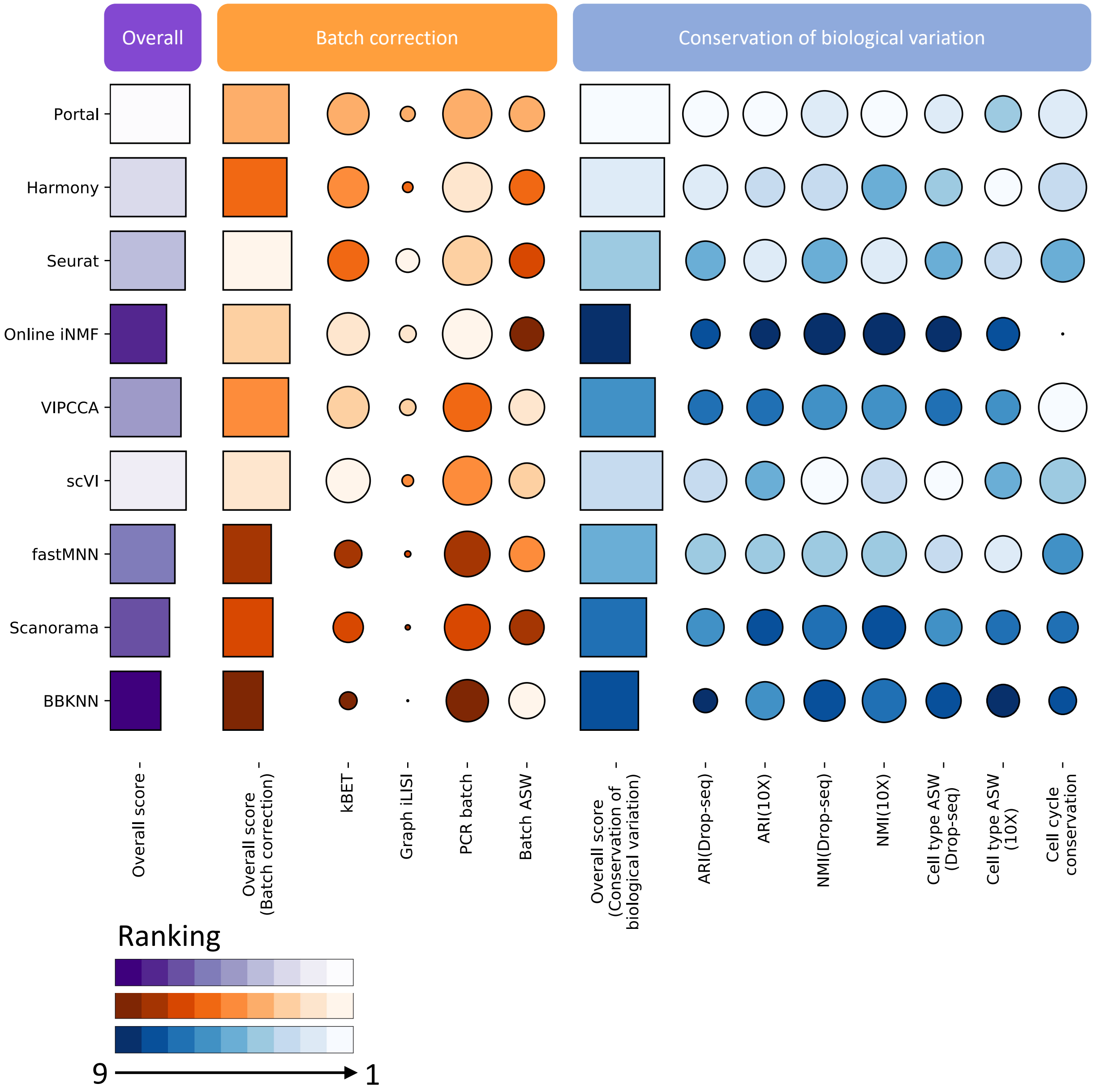


Figure S15: **Benchmarking of integration methods based on mouse hippocampus data.** Scores for batch correction and scores for conservation of biological variation are computed as the average of metrics which are in their categories. As the Drop-seq dataset and the 10X dataset provided detailed annotations in their original publications, we slightly modified the assessment for better evaluation of biological variation conservation. Overall scores are computed by a 40:60 weighted average of scores for batch correction and scores for conservation of biological variation.

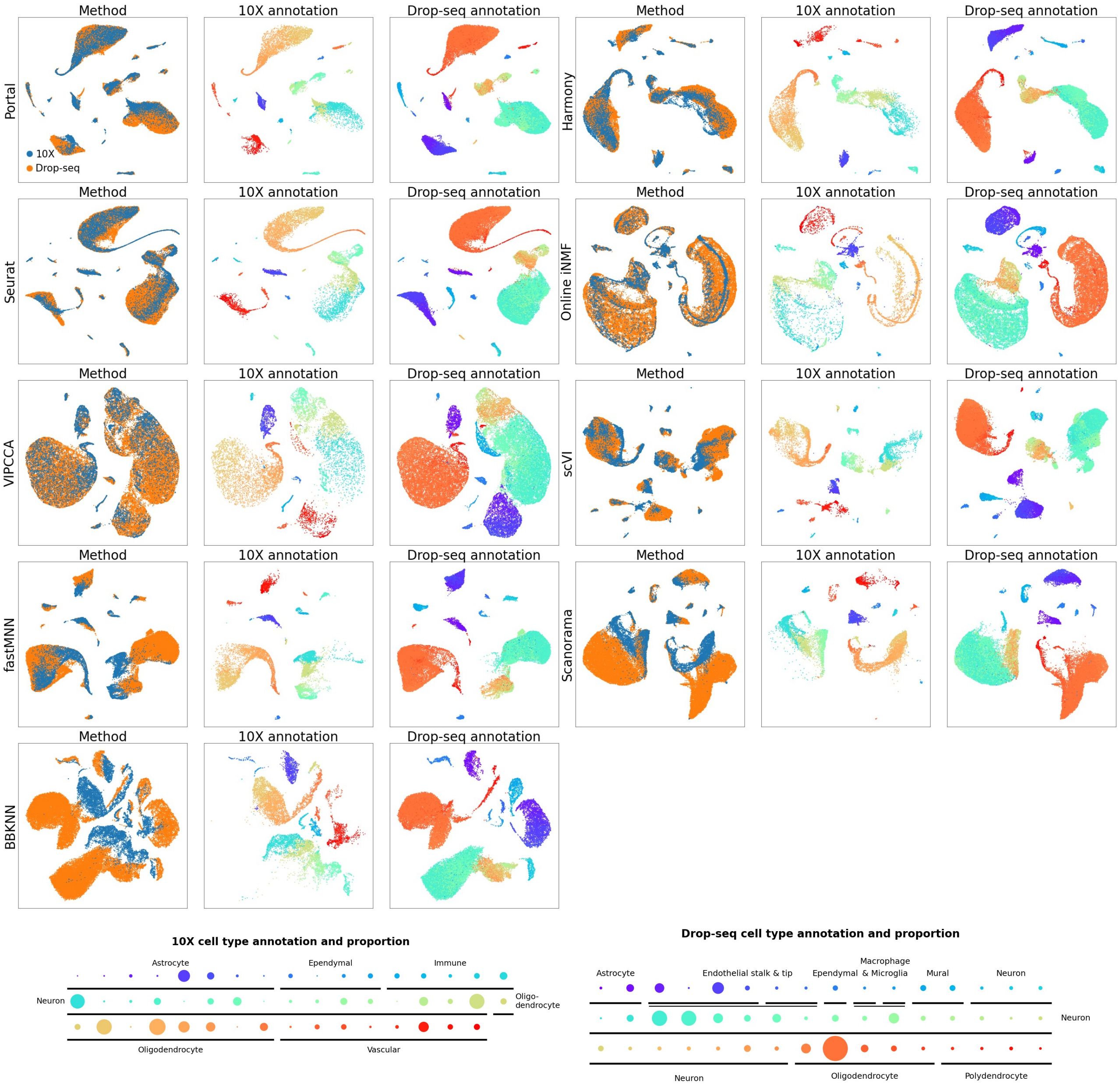


Figure S16: **Comparison of integration methods based on mouse thalamus data.** We integrated mouse thalamus scRNA-seq datasets profiled by 10X and Drop-seq. UMAP plots were colored by profiling methods and cell types, respectively. We used dot plots to visualize cell type labels and their proportions. Different colors represent different subpopulations of cells. The comparison among proportions of subpopulations was visualized by the sizes of corresponding dots.

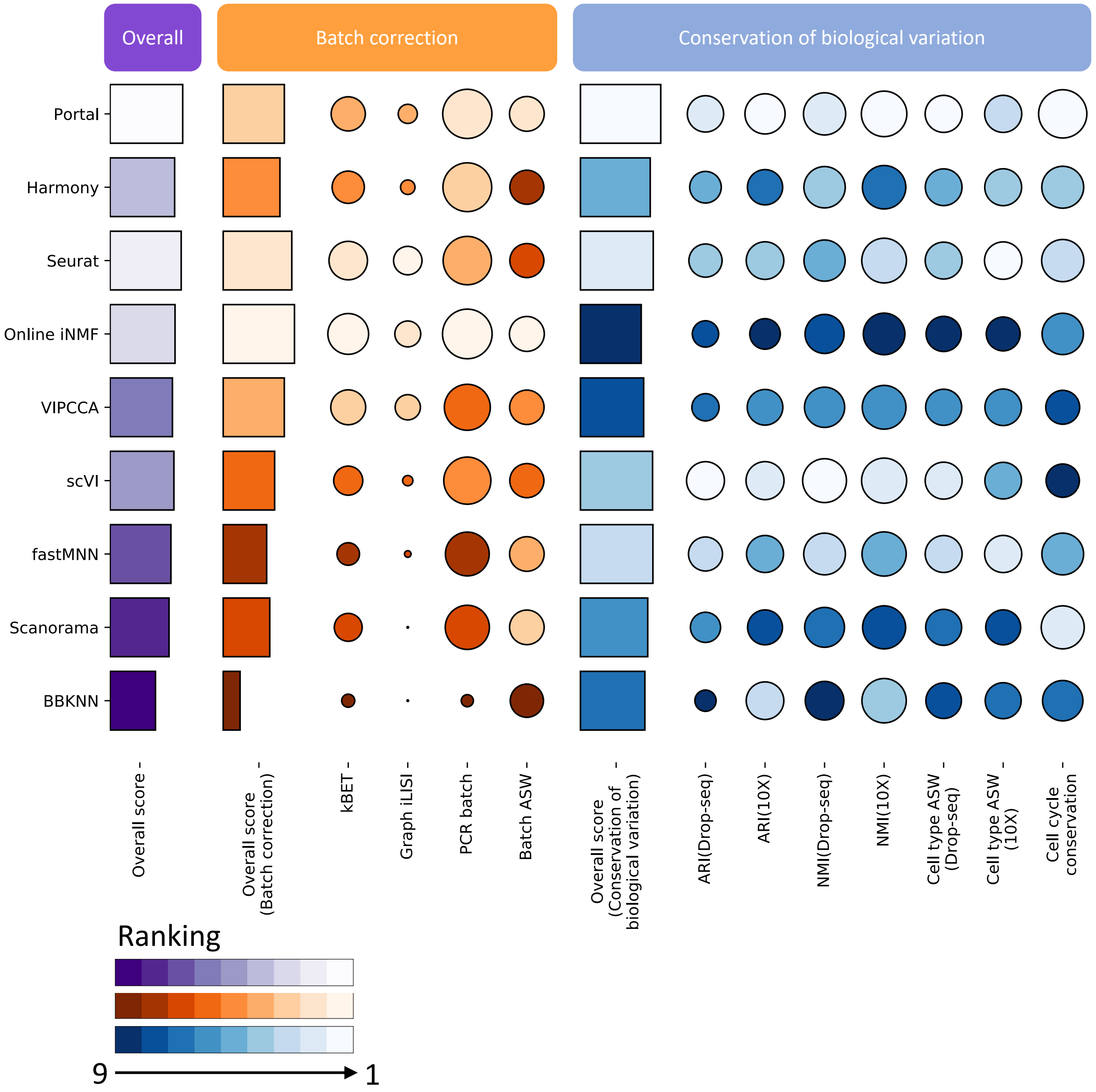


Figure S17: **Benchmarking of integration methods based on mouse thalamus data.** Scores for batch correction and scores for conservation of biological variation are computed as the average of metrics which are in their categories. As the Drop-seq dataset and the 10X dataset provided detailed annotations in their original publications, we slightly modified the assessment for better evaluation of biological variation conservation. Overall scores are computed by a 40:60 weighted average of scores for batch correction and scores for conservation of biological variation.

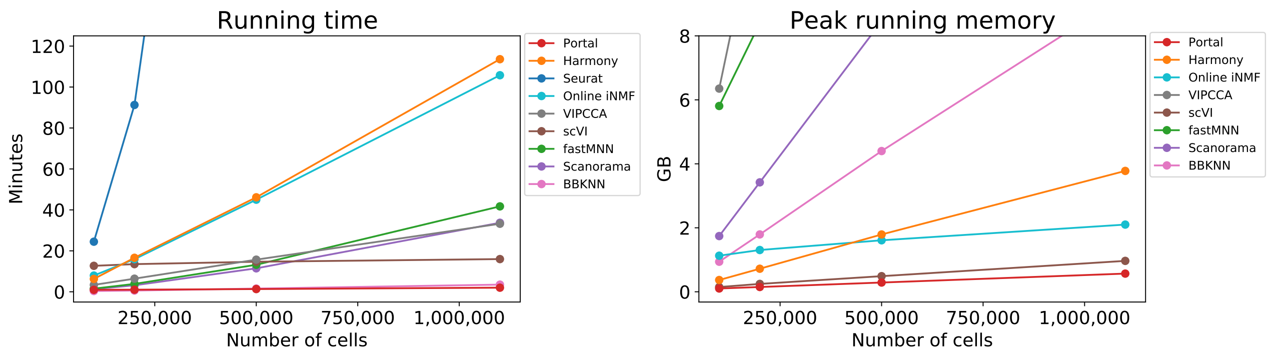


Figure S18: **Benchmarking of Portal, Harmony, Seurat, online iNMF, VIPCCA, scVI, fastMNN, Scanorama and BBKNN.** The running time and the peak running memory of all compared methods were evaluated based on datasets ($n = 100,000$, $250,000$, $50,000$, and $1,100,167$) sampled from two mouse brain atlas datasets. Among all compared methods, Portal and BBKNN were remarkably faster than other methods. However, BBKNN required much more memory usage than Portal as sample size increased. More importantly, BBKNN often provided less satisfactory integration performance as indicated by UMAP plots and quantitative evaluations in Figs. S5, S6, S7, S8, S9, S10, S11, S13, S15 and S17. Similar to BBKNN, the four methods VIPCCA, scVI, Scanorama and fastMNN also showed their comparatively less satisfactory performance compared to that of Portal, Harmony, Seurat and online iNMF (Fig. S5).

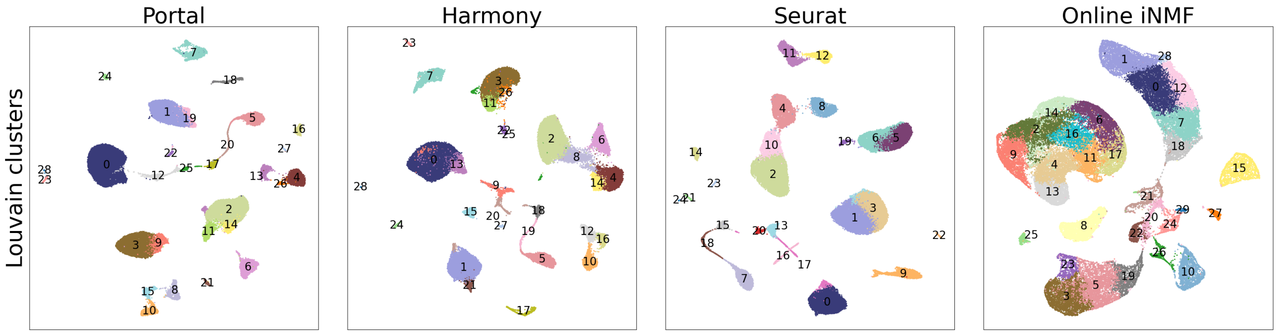


Figure S19: **Clusters identified by applying the Louvain algorithm to cell embeddings obtained by Portal, Harmony, Seurat and online iNMF after integration.** With default resolution setting, Louvain algorithm detected 29 (Portal), 29 (Harmony), 25 (Seurat), 30 (online iNMF) clusters as shown in UMAP plots. UMAP plots were drawn separately and colored by clusters identified in the cell embedding space of each method, respectively.

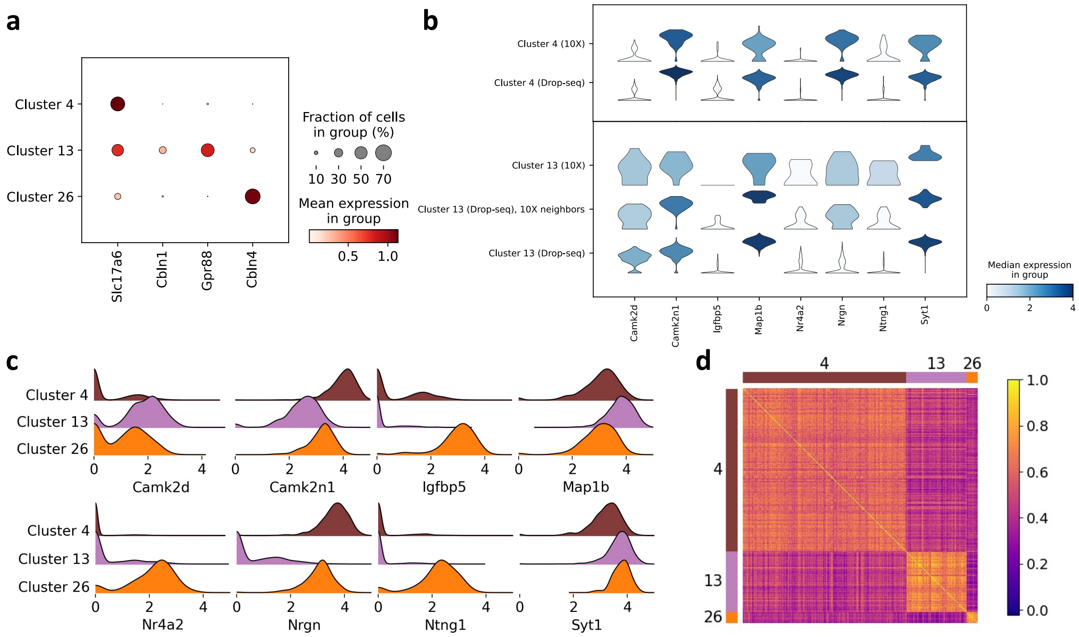


Figure S20: **Detailed verification of Portal's integration result on hippocampus datasets at transcriptome level. a**. We confirmed Portal's alignments of cluster 4, 13, 26 and the three neuron subpopulations by investigating the pattern of marker genes. **b**. The integration result from Portal was validated by the consistent pattern of differentially expressed genes across distinct clusters. Here we compared 10X cells with Drop-seq cells in clusters 4 and 13. 10X cells and Drop-seq cells in cluster 4 were compared directly. In cluster 13, as 10X cells only corresponded to a subset of Drop-seq cells, we compared 10X cells with their neighboring Drop-seq cells. In cluster 13, for each 10X cell, we searched its 5 nearest neighbors among Drop-seq cells. The collected Drop-seq cells were referred to as "10X neighbors" in the violin plot. **c**. We identified eight genes that showed distinct expression patterns across the three clusters. **d**. Transcriptional difference among the three clusters was further demonstrated by examining the correlation between cells using more genes.

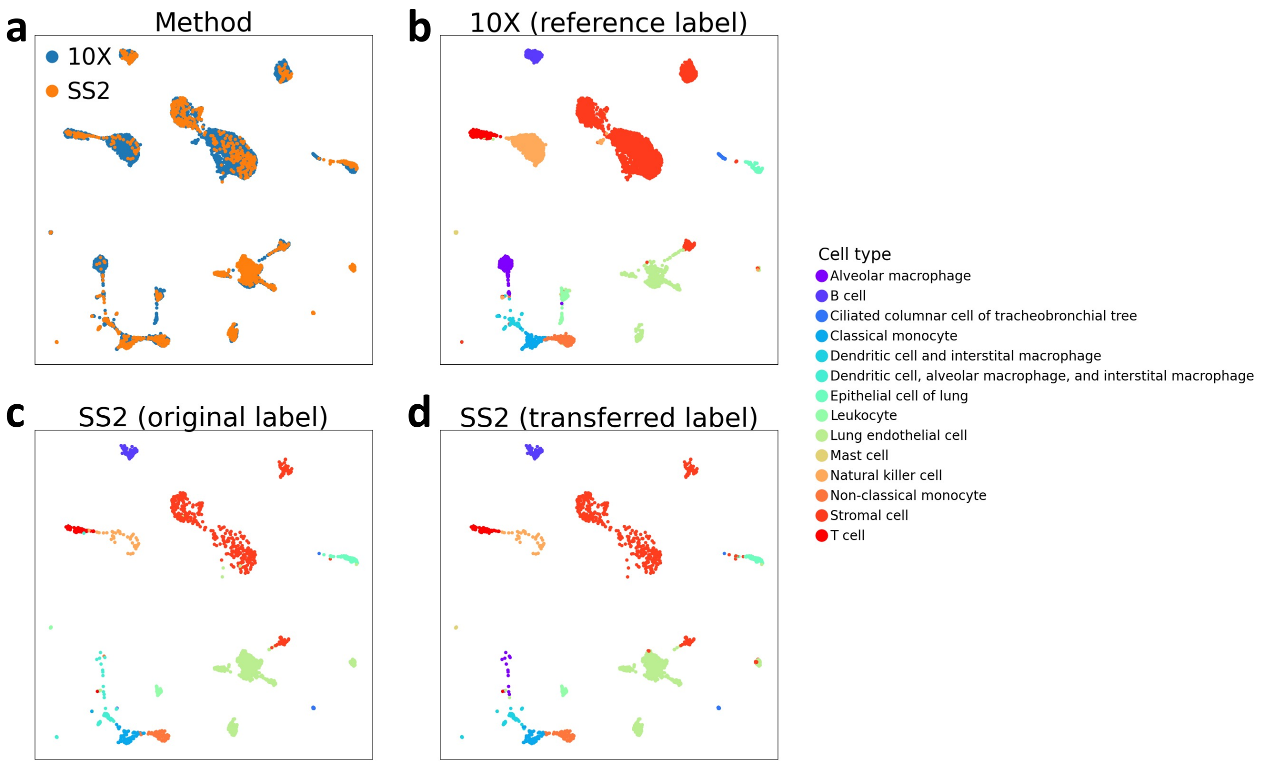


Figure S21: **Identification of rare subpopulations in mouse lung scRNA-seq data via label transfer.** Utilizing Portal's integration result (**a**), we transferred annotations from the 10X dataset (**b**) to the SS2 dataset (**d**). Portal's integration helped to identify fine-grained subpopulation alveolar macrophage (**d**), which was not identified in its original labels (**c**). **a**. UMAP plot of Portal's integration result colored by profiling methods. **b**, **c**. UMAP plots of integrated 10X, SS2 data colored by cell types obtained from their original publication [5]. **d**. UMAP plot of integrated SS2 data colored by transferred labels provided by Portal.

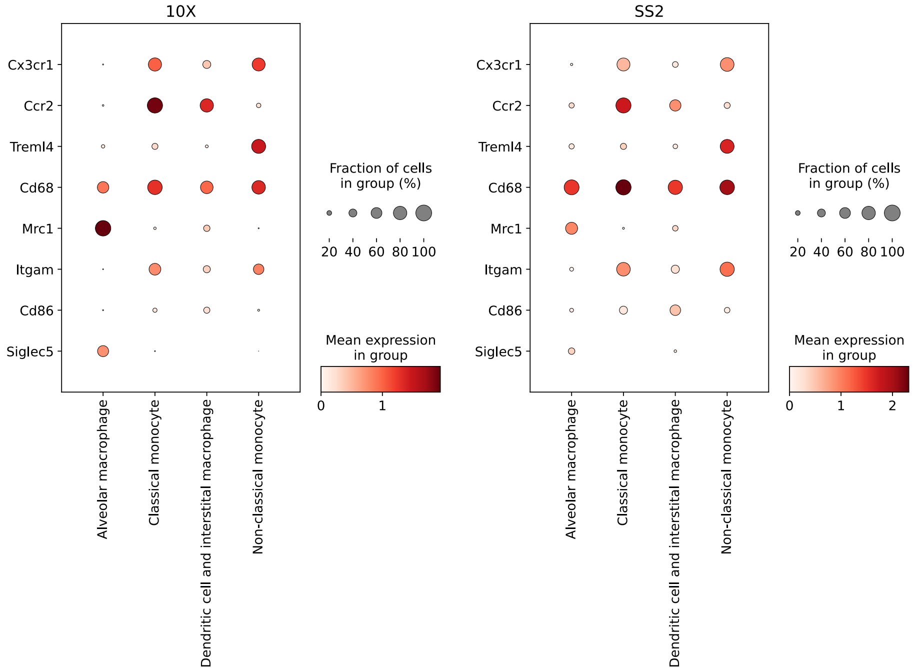


Figure S22: **Marker gene pattern of identified rare subpopulations in mouse lung scRNA-seq data.** By transferring labels from the 10X dataset to SS2 dataset, Portal identified four subpopulations of myeloid cells in SS2 dataset, including alveolar macrophage, dendritic cell and interstitial macrophage, classical monocyte, and non-classical monocyte. To validate the result, we examined four subpopulations' expression levels of marker genes: *Cd68* is a marker of macrophages and monocytes. Between classical monocytes and non-classical monocytes, *Ccr2* is a marker of classical monocytes, *Cx3cr1*, *Treml4* are markers of non-classical monocytes. Between alveolar macrophages and interstitial macrophages, *Mrc1*, *Siglec5* are markers of alveolar macrophages, *Itgam*, *Cd86* are markers of interstitial macrophages.

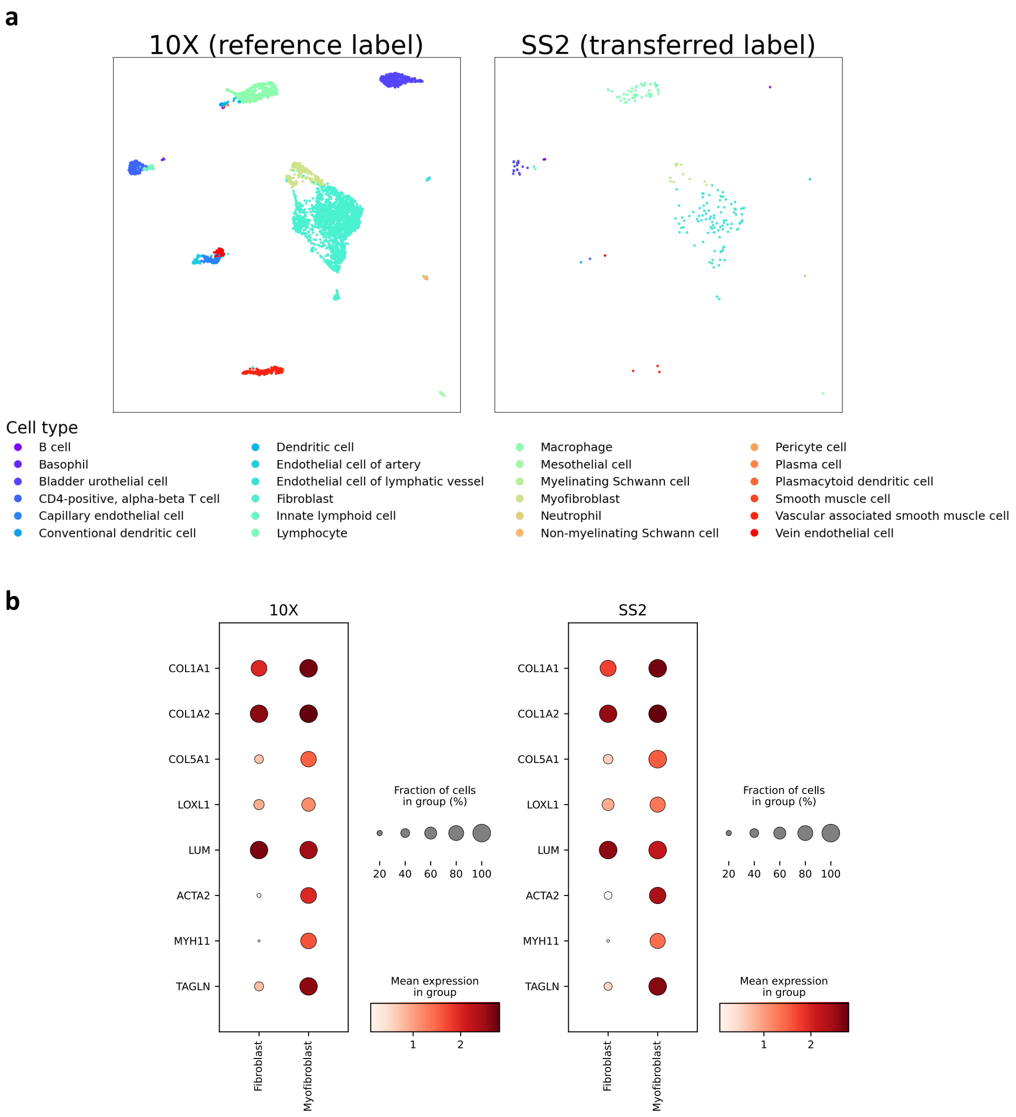


Figure S23: **Identification of myofibroblast subpopulation from fibroblast population in mouse lemur bladder scRNA-seq data via label transfer.** **a**. Portal utilized its integration result to transfer labels from the 10X dataset to the SS2 dataset. Portal successfully identified myofibroblast cells in SS2 dataset, although there were only 11 of them. **b**. We confirmed Portal's identification of myofibroblast cells by validating marker gene pattern. We collected eight marker genes: *COL1A1*, *COL1A2*, *COL5A1*, *LOXL1*, *LUM* are markers of fibroblast and myofibroblast cells. Compared to fibroblast cells, myofibroblast cells should have higher expression levels of markers *ACTA2*, *MYH11* and *TAGLN*.

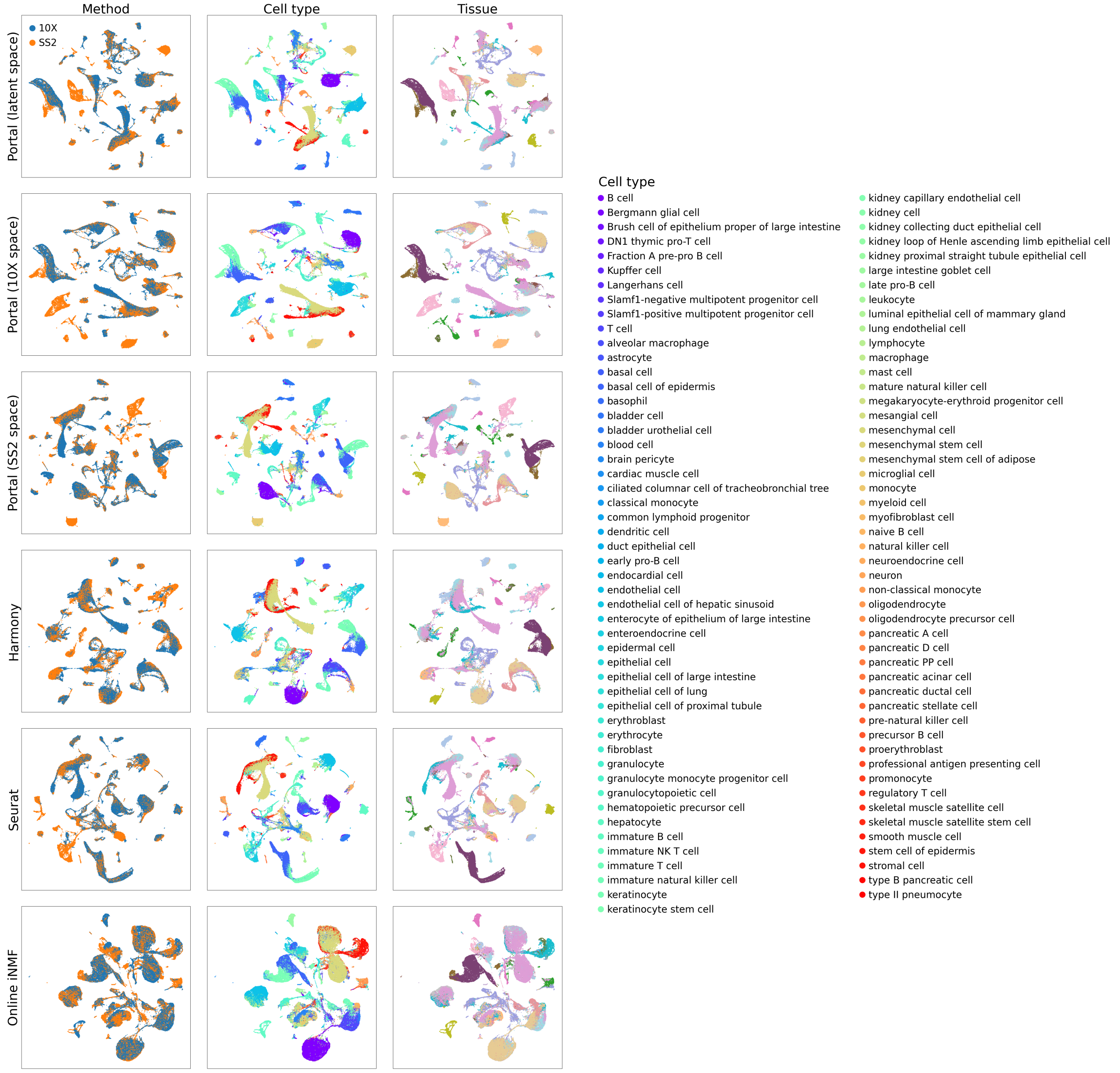


Figure S24: **Comparison of the capability of Portal, Harmony, Seurat and online iNMF to construct a comprehensive cell atlas across entire organism.** We applied the four integration approaches to harmonize the SS2 dataset and the 10X dataset from the Tabula Muris project, where mouse cells from 20 tissues were profiled. For a comprehensive investigation into Portal's performance, we visualized integration results of Portal in three spaces, namely shared latent space, 10X data space, and SS2 data space. Notably, among the 20 tissues, only 13 of them were included in the 10X data, while all of them were included in the SS2 data. In such a integration task, Portal preserved unique cell types contained in SS2 data, e.g., microglial cells, cell types in large intestine, and pancreatic islets cells. In contrast, Harmony, Seurat and online iNMF provided less accurate results, e.g., they incorrectly mixed microglial cells in brain myeloid with macrophage cells.

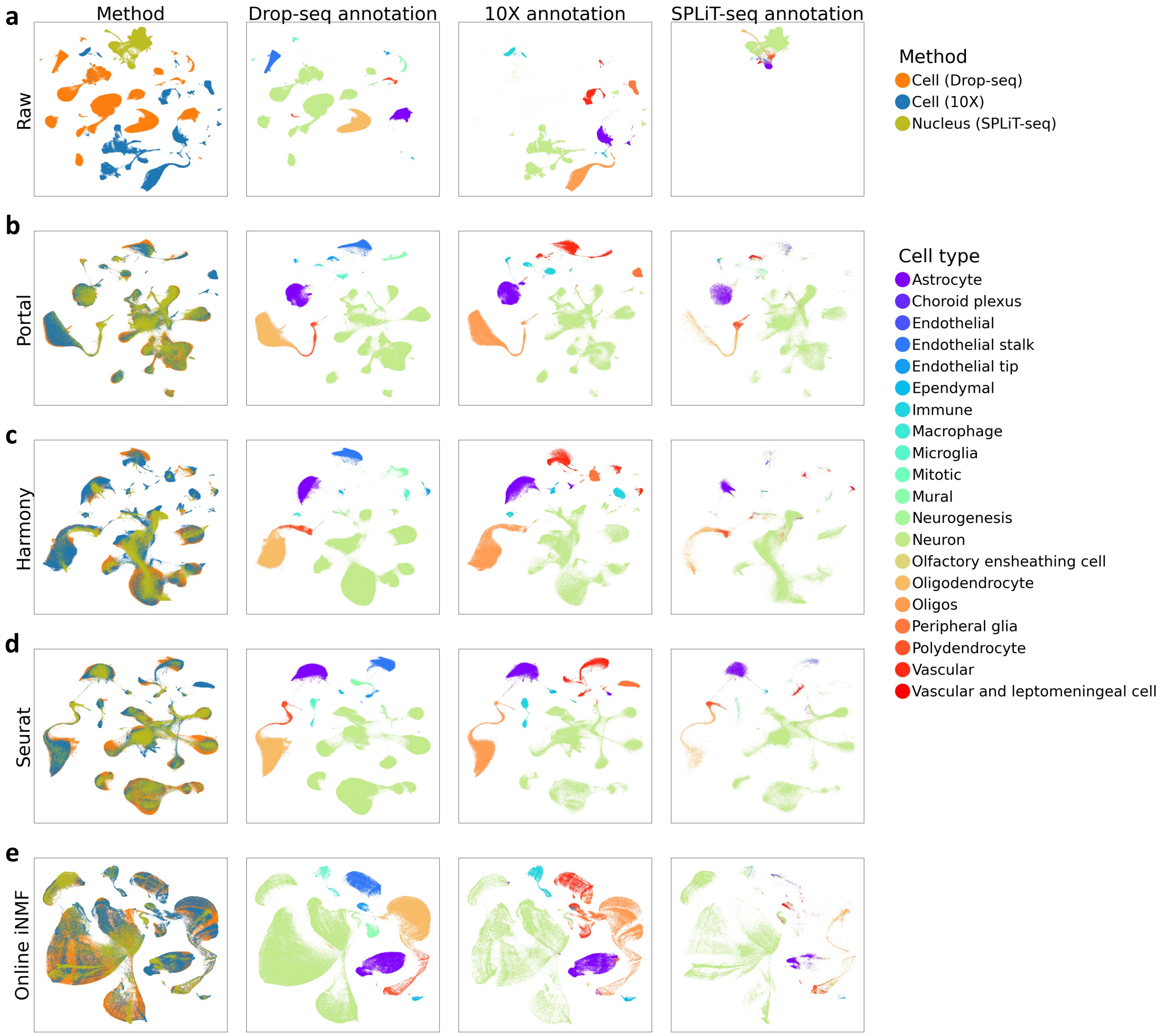


Figure S25: **Comparison of the ability of Portal, Harmony, Seurat and online iNMF to build alignment across one snRNA-seq dataset and two scRNA-seq datasets.** We applied the four integration approaches to align one snRNA-seq dataset profiled by SPLiT-seq [6], and two scRNA-seq datasets profiled by Drop-seq and 10X [7, 8]. We combined the cell type annotations provided by the three datasets together, although they contained slightly different annotations for non-neuron cells, e.g. immune cells and endothelial cells. **a-e**, UMAP visualizations of combined raw data (**a**), integration results of Portal (**b**), Harmony (**c**), Seurat (**d**) and online iNMF (**e**). UMAP plots were colored by profiling methods and cell types, respectively.

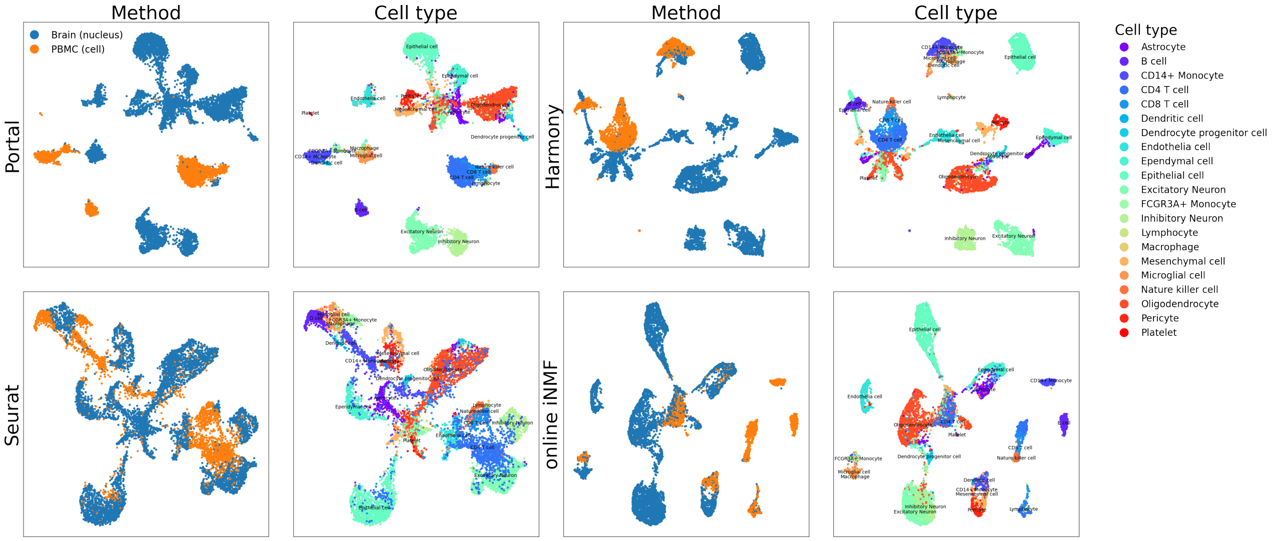


Figure S26: **Integration of human brain snRNA-seq dataset [9] and human blood scRNA-seq dataset [10] (example one).** We visualized the results with UMAP plots colored by datasets (profiling methods) and cell types, respectively. In this example, Portal preserved similarities among immune cells, and also prevented distinct cell types from being mixed. However, incorrect mixing was found in the other three methods: Harmony mixed B cells from blood with epithelial cells from brain; Seurat mixed CD4 T cells from blood with neurons from brain; online iNMF mixed dendritic cells and CD14+ monocytes from blood with mesenchymal cells from brain, and mixed CD4 T cells from blood with oligodendrocytes from brain.

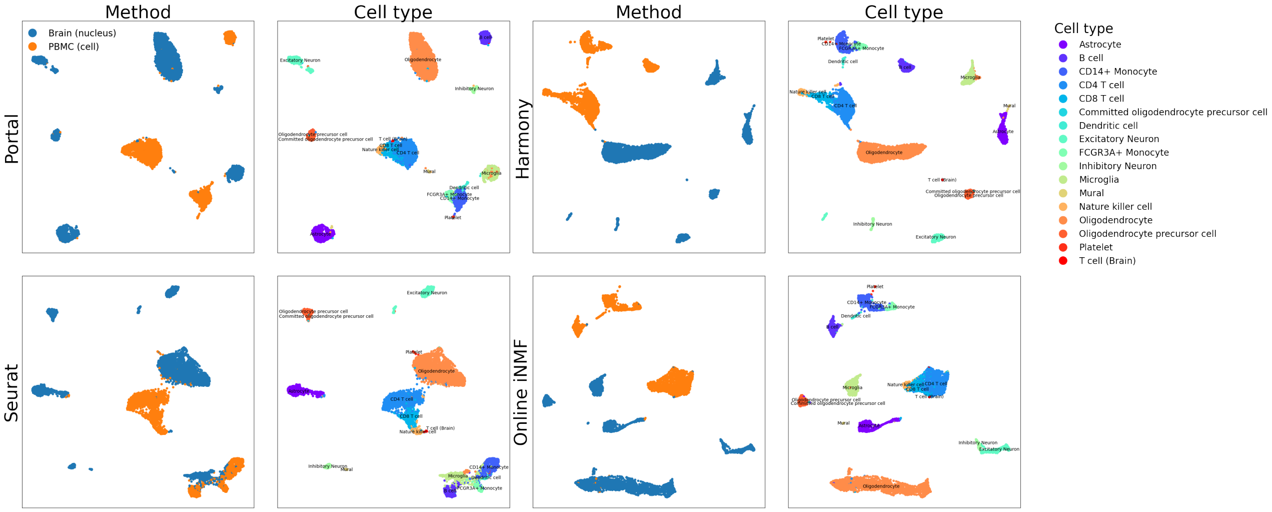


Figure S27: **Integration of human brain snRNA-seq dataset [11] and human blood scRNA-seq dataset [10] (example two).** We visualized the results with UMAP plots colored by datasets (profiling methods) and cell types, respectively. Portal aligned T cells from the two tissue types properly, while it also prevented distinct cell types from being mixed.

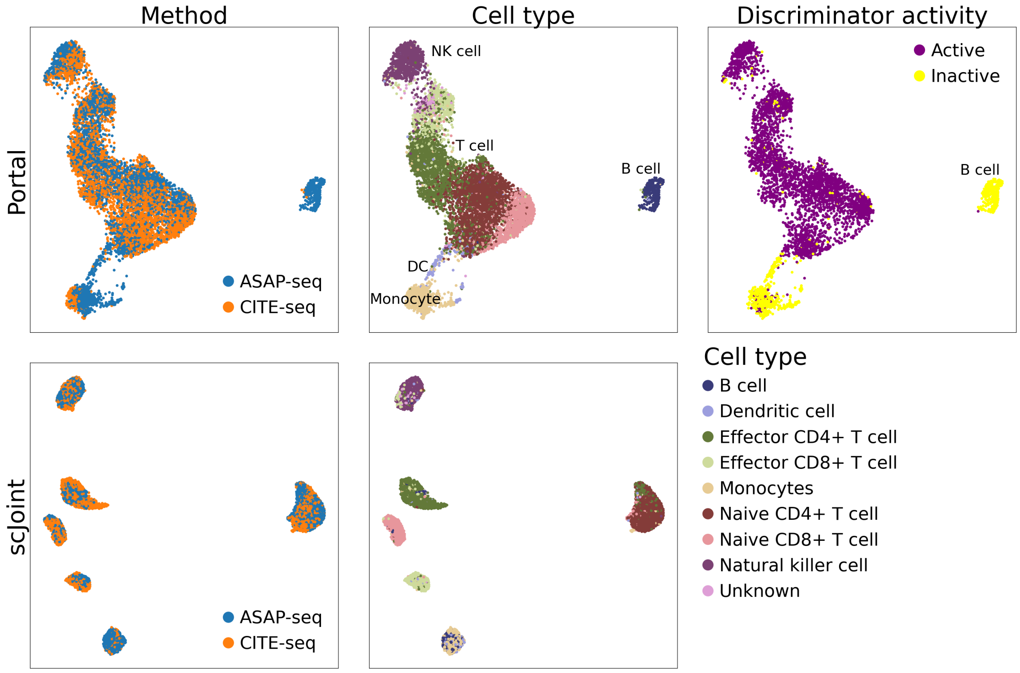


Figure S28: **Comparison of Portal and scJoint on the integration of scRNA-seq and scATAC-seq data from peripheral blood mononuclear cells (PBMCs) [12] where B cells were manually removed from the scRNA-seq dataset.** The scRNA-seq dataset and the scATAC-seq dataset were profiled by CITE-seq and ASAP-seq, respectively. We visualized the results with UMAP plots colored by profiling methods and cell types, respectively. For Portal's result, we also visualized the discriminator activity on the ASAP-seq cells. Portal protected B cells (unique in the ASAP-seq dataset) from being overcorrected by making the discriminator inactive on them.

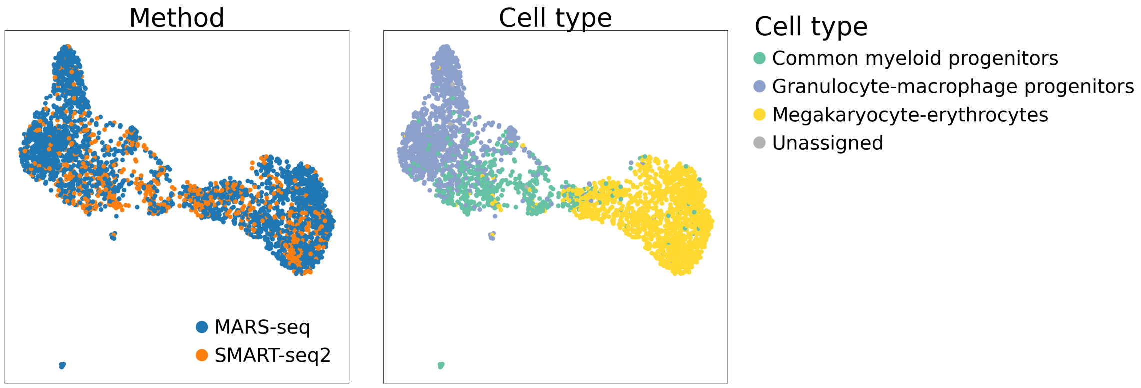


Figure S29: **Portal's integration of hematopoietic stem cells (HSC) differentiation datasets [13], [14].** The result was visualized by UMAP plots colored by profiling methods and cell types, respectively. The derivation of granulocyte-macrophage progenitors (GMP) and megakaryocyte-erythrocytes (MEP) from common myeloid progenitors (CMP) was still well preserved in Portal's integration result.

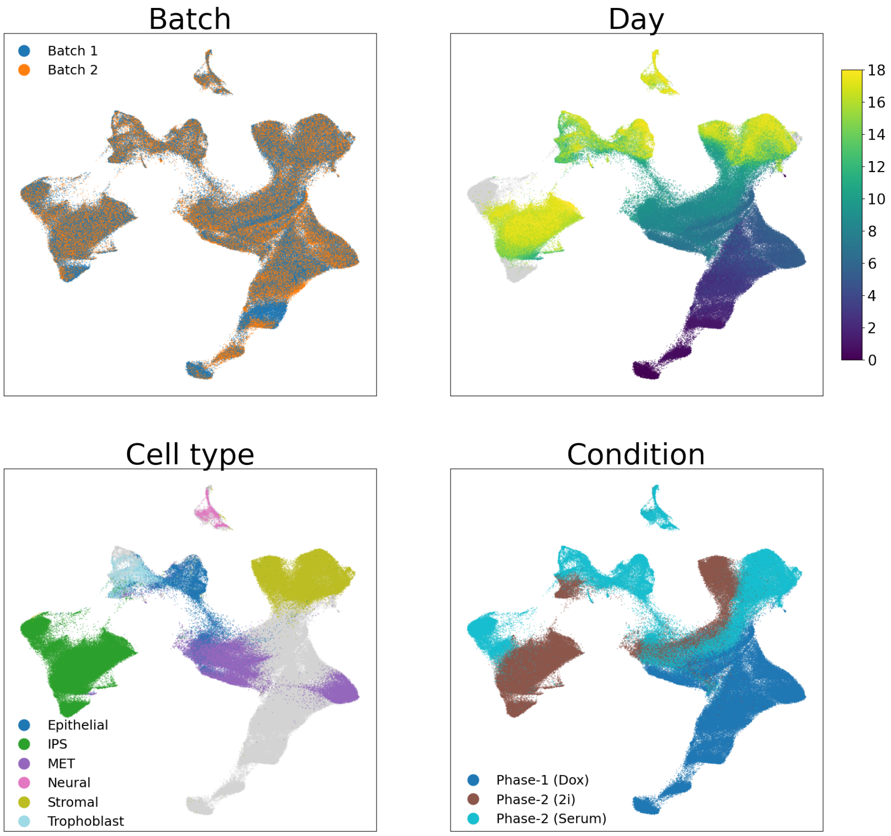


Figure S30: **Portal' s integration of induced pluripotent stem cells (iPSC) reprogramming datasets [15].** The result was visualized by UMAP plots colored by batches, time points (day), cell types, and conditions, respectively. In the UMAP plots colored by time point and cell type, cells in gray color were not assigned with the corresponding labels in the original publication. The gradual change of cell identities during the time period and the gradual changes in cellular identities that are associated with culture/reprogramming conditions were preserved in Portal's result.

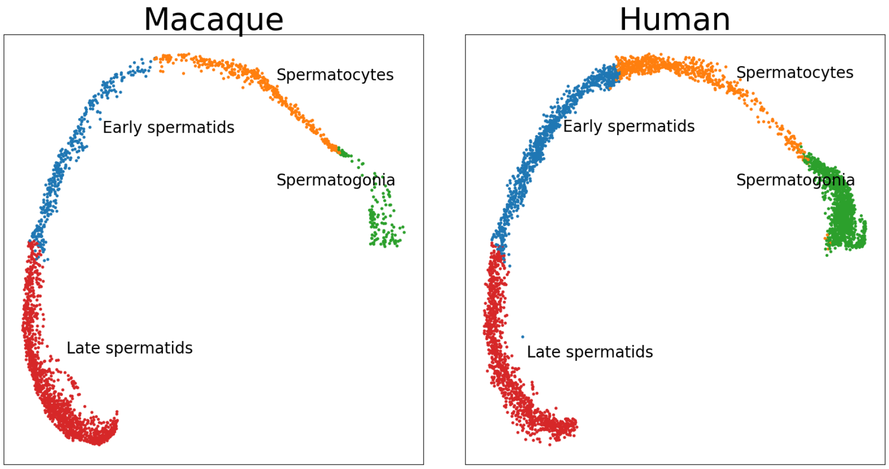


Figure S31: **UMAP visualization of the original published annotations for spermatogenesis data of macaque and human [16].**

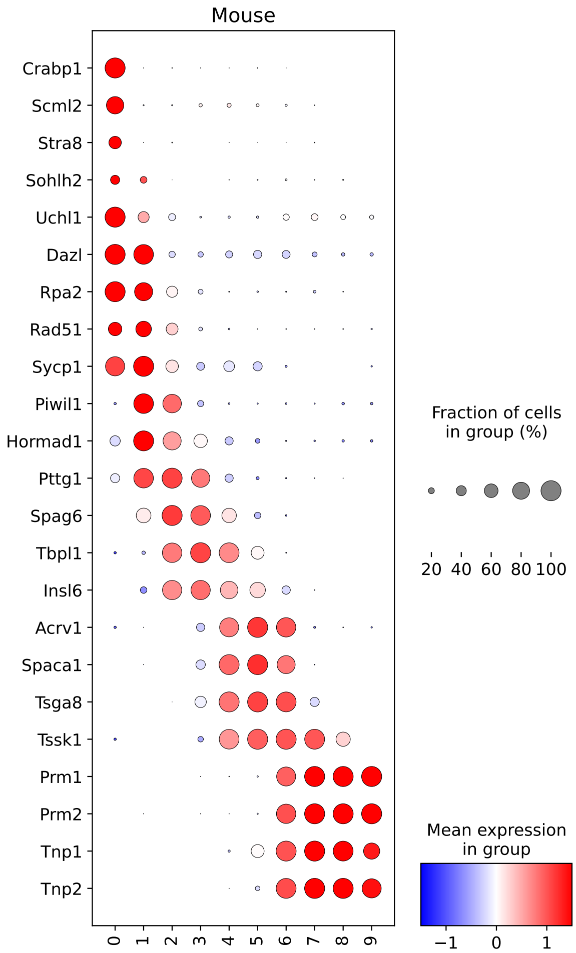


Figure S32: **Marker gene patterns of mouse in Louvain clusters in cross-species integration of spermatogenesis differentiation process.** For mouse, *Crabp1*, *Scml2*, *Stra8*, *Sohlh2*, *Uchl*, *Dazl*, *Rpa2*, *Rad51*, *Sycp1* are markers of spermatogonia, *Piwil1*, *Hormad1*, *Pttg1*, *Spag6*, *Tbpl1*, *Insl6* are markers of spermatocytes, *Acrv1*, *Spaca1*, *Tsga8*, *Tssk1* are markers of early spermatids, and *Prm1*, *Prm2*, *Tnp1*, *Tnp2* are markers of late spermatids.

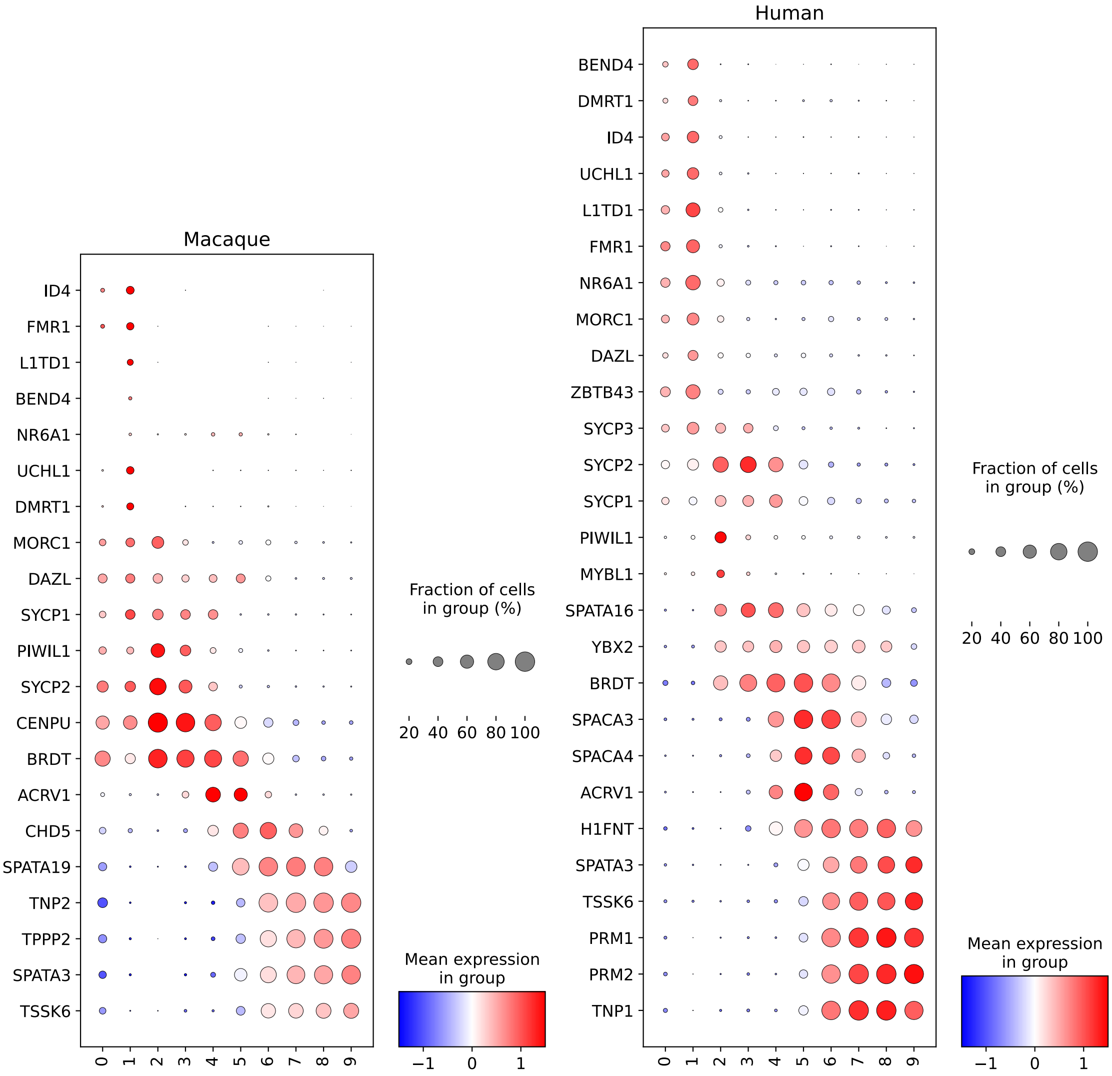


Figure S33: **Marker gene patterns of macaque and human in Louvain clusters in cross-species integration of spermatogenesis differentiation process.** For macaque, *ID4*, *FMR1*, *L1TD1*, *BEND4*, *NR6A1*, *UCHL1*, *DMRT1*, *MORC1*, *DAZL* are markers of spermatogonia, *SYCP1*, *PIWIL1*, *SYCP2*, *CENPU* are markers of spermatocytes, *BRDT*, *ACRV1*, *CHD5*, *SPATA19* are markers of early spermatids, and *TNP2*, *TPPP2*, *SPATA3*, *TSSK6* are markers of late spermatids. For human, *BEND4*, *DMRT1*, *ID4*, *UCHL1*, *L1TD1*, *FMR1*, *NR6A1*, *MORC1*, *DAZL*, *ZBTB43*, *SYCP3* are markers of spermatogonia, *SYCP2*, *SYCP1*, *PIWIL1*, *MYBL1*, *SPATA16*, *YBX2* are markers of spermatocytes, *BRDT*, *SPACA3*, *SPACA4*, *ACRV1*, *H1FNT* are markers of early spermatids, and *TSSK6*, *PRM1*, *PRM2*, *TNP1*, *SPATA3* are markers of late spermatids. The marker gene patterns validated the label transfer results given by Portal.

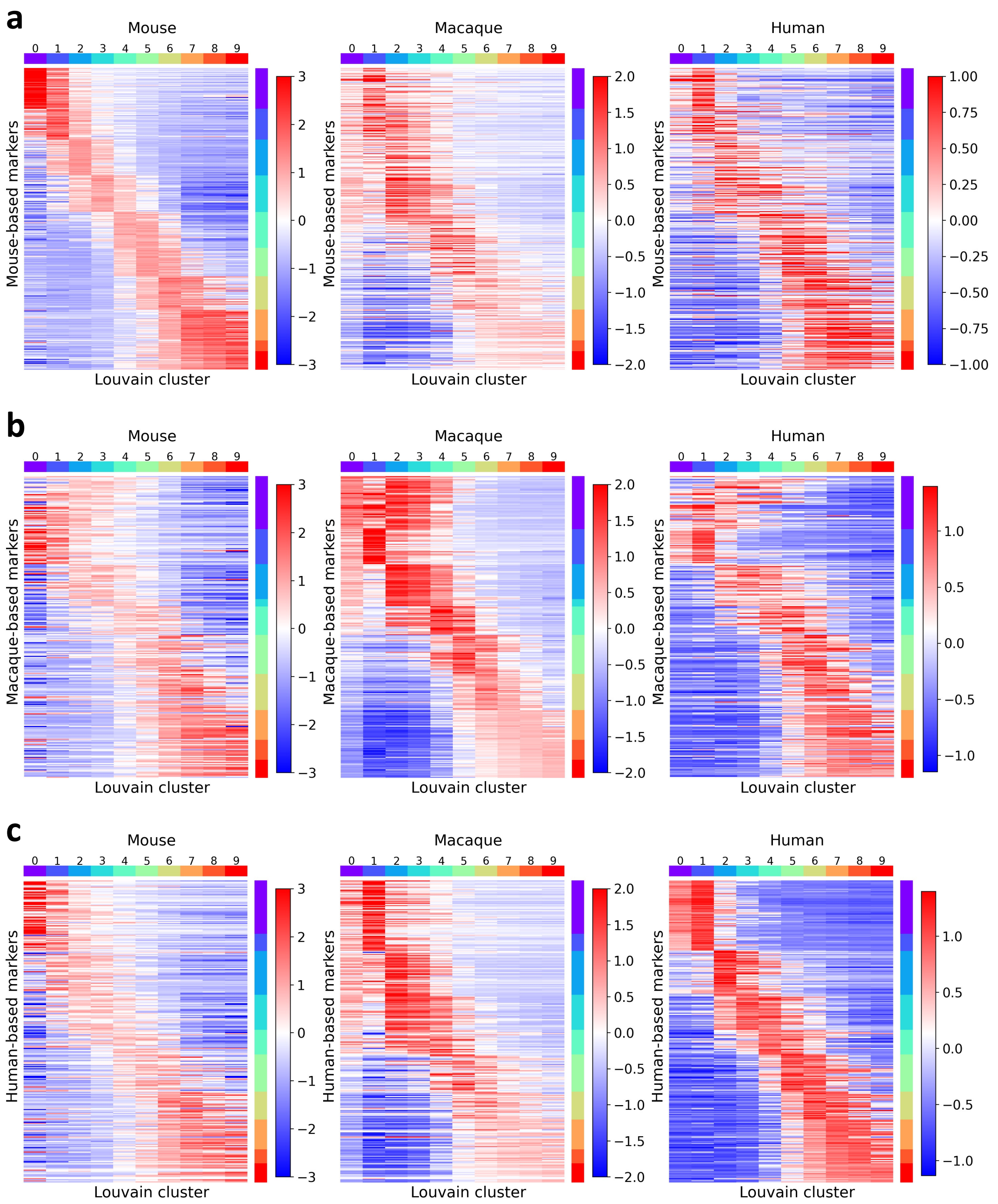


Figure S34: **Gene expression heatmaps in Louvain clusters in cross-species integration of spermatogenesis differentiation process.** For each species, we selected highly-expressed genes for each cluster and combined them together. Gene expression patterns on genes selected based on mouse (**a**), macaque (**b**) and human (**c**) showed connection and distinction among spermatogenesis differentiation processes of different species.

### Supplementary Information: Tables

#### Network structures in Portal

| Layer | Detail | Input Size | Output Size | Number of Parameters |
|---|---|---|---|---|
| Fully Connected | Linear | 30 | 512 | $30 \times 512 + 512 = 15,872$ |
| | RELU | 512 | 512 | 0 |
| Fully Connected | Linear | 512 | 20 | $512 \times 20 + 20 = 10,260$ |

Table S1: **Structure of Portal's encoder networks.**

| Layer | Detail | Input Size | Output Size | Number of Parameters |
|---|---|---|---|---|
| Fully Connected | Linear | 20 | 512 | $20 \times 512 + 512 = 10,752$ |
| | RELU | 512 | 512 | 0 |
| Fully Connected | Linear | 512 | 30 | $512 \times 30 + 30 = 15,390$ |

Table S2: **Structure of Portal's generator networks.**

| Layer | Detail | Input Size | Output Size | Number of Parameters |
|---|---|---|---|---|
| Fully Connected | Linear | 30 | 512 | $30 \times 512 + 512 = 15,872$ |
| | RELU | 512 | 512 | 0 |
| Fully Connected | Linear | 512 | 512 | $512 \times 512 + 512 = 262,656$ |
| | RELU | 512 | 512 | 0 |
| Fully Connected | Linear | 512 | 1 | $512 \times 1 + 1 = 513$ |

Table S3: **Structure of Portal's discriminator networks.**

[8] Amit Zeisel, Hannah Hochgerner, Peter Lönnerberg, Anna Johnsson, Fatima Memic, Job

Van Der Zwan, Martin Häring, Emelie Braun, Lars E Borm, Gioele La Manno, et al. Molecular architecture of the mouse nervous system. *Cell*, 174(4):999–1014, 2018.

[9] John F Fullard, Hao-Chih Lee, Georgios Voloudakis, Shengbao Suo, Behnam Javidfar, Zhiping Shao, Cyril Peter, Wen Zhang, Shan Jiang, André Corvelo, et al. Single-nucleus transcriptome analysis of human brain immune response in patients with severe covid-19. *Genome medicine*, 13(1):1–13, 2021.

[10] 3k peripheral blood mononuclear cells (PBMCs) from a healthy donor from 10X Genomics. https://support.10xgenomics.com/single-cell-gene-expression/datasets/1.1.0/pbmc3k.
